# The G-protein coupled receptor GPR34 maintains the homeostatic state of microglia and restrains the disease-associated microglial response in an Alzheimer’s disease model

**DOI:** 10.1101/2025.11.26.690816

**Authors:** Ana Geller, Min Jee Kwon, Sean K. Simmons, Qihong Xu, William E. Martenis, Jordan Doman, Sahana Natarajan, Katherine J. Stalnaker, Jixiang Zhang, Xian Adiconis, Kwanho Kim, Trang Nguyen, Matthew Demers, Donovan Batzli, Antia Valle-Tojeiro, Christy Biji, Sameer Aryal, Horia Pribiag, Constanze Depp, Nader Morshed, Deeksha Misri, Diana Bohannon, Elena Longhi, Hasmik Keshishian, Steven A. Carr, David McKinney, Alexandra E. Gould, Evan Lebois, Michel Weïwer, Beth Stevens, Matthew Johnson, Joshua Z. Levin, Yan-Ling Zhang, Morgan Sheng, Prabhat S. Kunwar

**Affiliations:** The Stanley Center for Psychiatric Research, The Broad Institute of MIT and Harvard, Cambridge, MA, USA; Department of Medicine, University of Massachusetts Chan Medical School, Worcester, MA, USA; F.M. Kirby Neurobiology Center, Boston Children’s Hospital, Boston, MA, USA; Society of Fellows, Harvard University, Cambridge, MA 02138, USA; Neuroscience Program, University of California San Francisco; The Proteomics Platform, The Broad Institute of MIT and Harvard, Cambridge, MA, USA; The Center for the Development of Therapeutics, The Broad Institute of MIT and Harvard; Violet Therapeutics, Cambridge, MA, USA; Howard Hughes Medical Institute, Harvard Medical School, Boston, MA, USA; Department of Brain and Cognitive Sciences, Massachusetts Institute of Technology, Cambridge, MA, USA

## Abstract

Expression of the G protein coupled receptor GPR34 is highly enriched in microglia and is downregulated in various brain diseases, including Alzheimer’s disease (AD) and multiple sclerosis (MS). GPR34 function is poorly understood, as is its role in regulation of microglial states. Using RNA-sequencing, we find that microglia from *Gpr34* knockout (KO) mouse brains exhibited a transcriptomic shift toward disease-associated microglia (DAM) and inflammatory profiles, partially resembling the microglial phenotype seen in 5xFAD AD model mice. Moreover, when *Gpr34* KO mice were crossed with 5xFAD mice, the DAM transcriptional profile of microglia and glial pathology were further exacerbated beyond the already robust DAM signature driven by 5xFAD alone. This occurred without affecting amyloid plaque burden. Human stem cell-derived microglia (iMGLs) lacking *GPR34* showed reduced calcium (Ca²⁺) and phosphorylated ERK (pERK) signaling in response to stimulation with known GPR34 agonists (lyso-phosphatidylserine (lysoPS) and myelin), as well as transcriptomic changes in immune regulation and cell proliferation related pathways. Interestingly, *GPR34* KO iMGLs were selectively impaired in phagocytosis of myelin but not amyloid-β (Aβ) or *E. coli,* and showed a diminished transcriptional response elicited by myelin. Together, these findings suggest that GPR34 is important for maintaining microglia in a homeostatic state, promotes phagocytosis of and transcriptional response to myelin, and limits microglial reactivity in neurodegenerative disease conditions.

## Introduction

Microglia are resident immune cells of the brain that play a key role in maintaining brain homeostasis and health. They perform diverse functions such as synaptic elimination, debris clearance, immune surveillance, and sculpting neuronal activity and plasticity^1–7^. Microglial dysfunction is a key feature of neurodegenerative and myelin-related diseases such as Alzheimer’s disease (AD) and multiple sclerosis (MS)^4–6,8,9^. For example, in AD, microglia can be maladaptive, exhibiting changes in gene expression and function such as impaired clearance of myelin debris and amyloid*-β*, chronic pro-inflammatory responses, and excessive elimination of neuronal synapses. Notably, the expression of many risk genes for AD are highly enriched in microglia^2,10,11^; however, the signals and mechanisms that regulate the transition of microglia between homeostatic, adaptive, and pathological states remain poorly understood^1,4,8,9^.

G-protein coupled receptors (GPCRs) mediate various cell-environment interactions and represent the largest set of currently druggable therapeutic targets. Several GPCRs are selectively expressed in microglia, but their functions are poorly understood, despite their potential relevance for therapeutic intervention^12,13^. One such receptor is GPR34, which is a member of the rhodopsin-like family of GPCRs. *GPR34* is highly and specifically expressed in microglia and macrophages, but its biological role remains largely undefined^13,14^.

*GPR34* expression in microglia is downregulated under various disease conditions such as AD and MS^14,15^. Further, GPR34 is reported to play a role in maintaining microglial homeostasis, regulating lipid responses, and modulating microglia states and disease pathology^16–23^. However, the identity of its endogenous ligand(s) remains uncertain; lysoPS has been proposed as a candidate ligand, but because it activates multiple receptors, its biological effects cannot be specifically attributed to GPR34^24–28^. Studies report conflicting functions for GPR34: some protective, others detrimental against AD pathology^16,18,20–23,29^. Several reports show that GPR34 inhibition alleviates neuroinflammation and promotes amyloid clearance in mice^16,20,21^, whereas another reported that GPR34 agonism can enhance *in vivo* amyloid clearance^23^. These discrepancies highlight the limited understanding of the upstream signals and downstream pathways regulating GPR34 activity in microglia, as well as its **context-dependent functions**.

To address the *in vivo* function of GPR34, we first studied transcriptomic changes in *Gpr34* KO mouse brains by RNA sequencing (RNA-seq). To investigate GPR34’s role in an Alzheimer’s disease model, we studied the brain and microglia of *Gpr34* KO mice crossed to the 5xFAD mouse model of pathology^30,31^. Human iPSC-derived induced microglia-like cells (iMGLs) null for *GPR34* were generated and used to dissect downstream signaling of GPR34 in response to various ligands, and to evaluate the resulting effects on gene expression and phagocytosis. We find that *Gpr34* deletion in mice shifts microglia toward inflammation- and DAM-related gene expression signatures, with some resemblance to 5xFAD microglia. In 5xFAD mice, *Gpr34* KO exacerbates the disease-associated features, including heightened glial reactivity. In iMGLs, *GPR34* deletion abolishes downstream signaling in response to both pharmacological agonists and myelin, and diminishes functional responses, including myelin-induced transcriptional effects and phagocytosis. Thus, by utilizing both *in vivo* and *in vitro* models, we were able to clarify how GPR34 influences microglial transcriptional states, signaling, and function in both normal physiologic and chronic disease settings.

## Results

### *Gpr34* knockout in mice shifts microglial transcriptional profiles toward DAM- and inflammation-associated signatures

GPR34 is expressed highly and selectively in microglia within the brain in humans and mice (**Supplemental Figure 1)**^13,14^. To determine how GPR34 loss affects microglial transcriptional programs, we performed single-cell RNA sequencing (scRNA-seq) of isolated microglia and brain border-associated macrophages (BAMs) from hippocampal tissue from 6-month-old wild-type (WT) and *Gpr34* KO mice, on either a normal C57BL/6 background or in 5xFAD transgenic mice, resulting in 4 conditions: WT, *Gpr34* KO (which refers to *Gpr34 -/Y,* as *Gpr34* gene is X-linked), 5xFAD, and *Gpr34* KO;5xFAD (**Figure 1A-B; see Methods**). We also performed single-nucleus RNA sequencing (snRNA-seq) of hippocampal tissue from the same mice **(contralateral hemisphere)** to study the effect of GPR34 loss on the non-microglial cells. Clustering, quality control, and cell type annotation were performed on both the single-cell and single-nucleus data (**Figure 1B, Supplemental Figure 2 and 3**). After standard quality control across samples of the microglia scRNA-seq data, we identified distinct subpopulations as described from previous studies^9,32^, including a homeostatic state, DAM (which, more specifically, Amyloid-DAMs, as associated with amyloid-disease in 5xFAD mice)^1^, interferon-responsive microglia, and a *Ms4a7+* macrophage population representing brain-border associated macrophages^1^, and cycling microglia (**Supplemental Figure 3A-B**). We then first focused on the comparison between WT and *Gpr34* KO mice to define the transcriptional consequences of Gpr34 loss in the absence of 5xFAD pathology.

**Figure 1:**
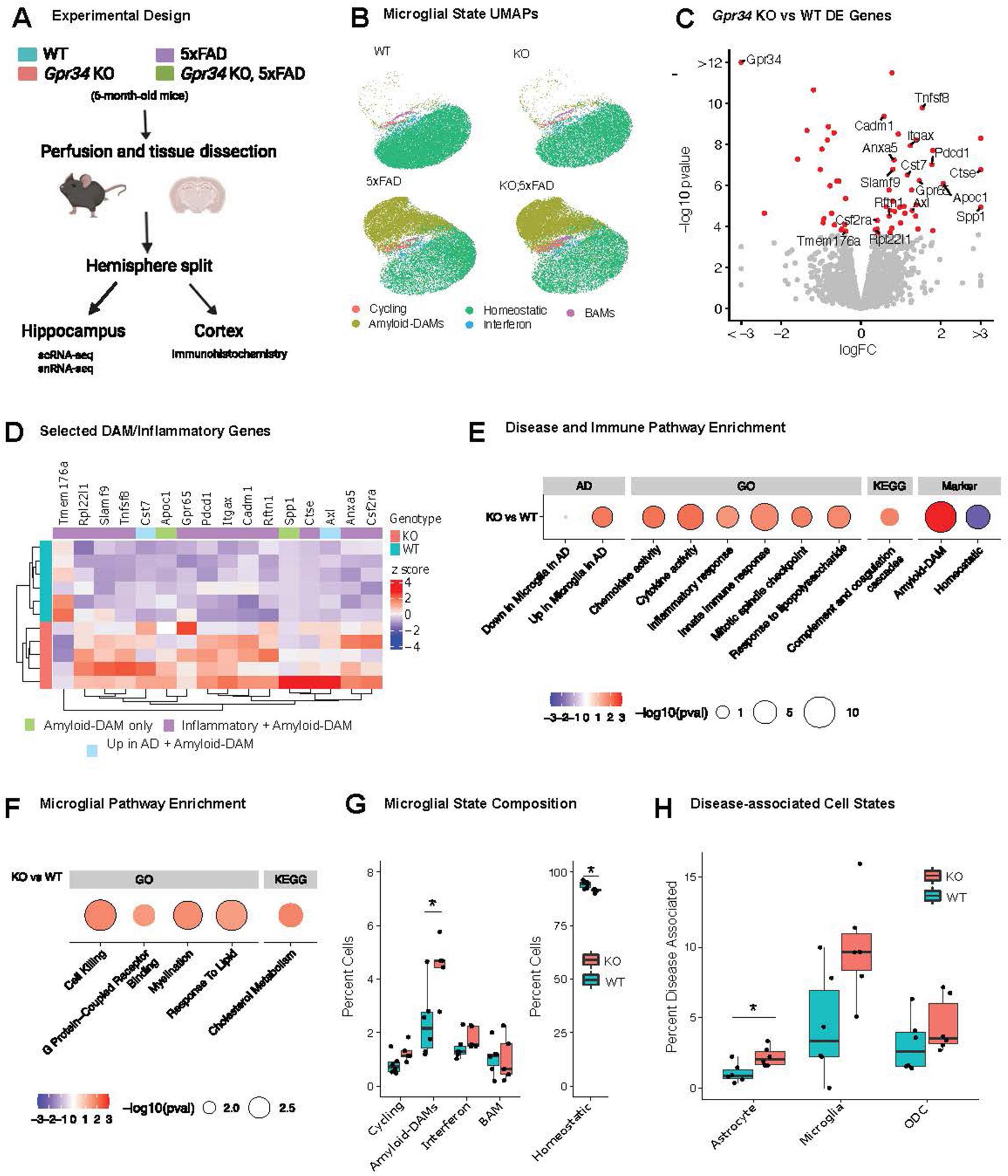
*Gpr34* knockout in mice shifts microglial transcriptional profiles toward DAM-and inflammation-associated signatures, partially resembling 5xFAD Alzheimer’s model transcriptomic profile. **A:** Overview of the experimental design. |**B:** UMAP of single-cell microglia data, colored by microglial states. Each condition (*Gpr34* KO, WT, 5xFAD, and *Gpr34* KO;5xFAD) is shown separately. **C:** Volcano plot of differentially expressed (DE) genes between *Gpr34* KO and WT mice in the single-cell data, calculated with pseudobulk. Those colored red are significant. Labelled genes are known to be DAM specific based on previous publications. **D:** Heatmap of selected DE genes. The rows are mice, columns are genes, the color is the z score of the pseudobulked TPM per mouse. The bars on the left represent *Gpr34* KO vs WT, the bars on the top represent which gene set each gene belongs to. All are significant at an FDR of 0.05. **E:** Enrichment analysis of chosen terms related to disease and the immune processes. The terms (see Methods) are on the x-axis, color is Normalized Enrichment Score (NES), size is –log_10_ pvalue, those outlined in black have FDR<0.1. The dataset of origin for each term is annotated on the top. **F:** Enrichment analysis of chosen terms related to microglia function. The terms are on the x-axis, color is NES, size is –log_10_ pvalue, those outlined in black have FDR<0.1. The dataset of origin for each term is annotated on the top. **G:** Percent of cells (y-axis) in each microglia cell type (x-axis) in each *Gpr34* KO and WT mouse (color). There is a significant (FDR<0.05) reduction in homeostatic cells and increase in DAMs in the *Gpr34* KO mice, represented by the asterisks. **H:** Percent of disease associated cells (y-axis) in astrocyte, microglia and oligodendrocyte (ODC) populations (x-axis) in each KO and WT mouse (color). We see an increase in all of them, though it only reaches significance in the astrocytes (FDR<0.05), represented by the asterisk. **Sample sizes for sequencing analyses:** For single-cell RNA-seq, WT, n = 6 mice; *Gpr34* KO, n = 5 mice; and 5xFAD, n = 5 mice. For single-nucleus RNA-seq, n = 6 mice per condition.

In *Gpr34* KO mice, we verified data quality across samples (**Supplemental Figure 2B and Supplemental Table 1**) and confirmed the absence of *Gpr34* transcripts in *Gpr34* KO microglia (**Supplemental Figure 3C**). Differential expression analysis of scRNA-seq from all isolated microglia comparing *Gpr34* KO to WT revealed 60 differentially expressed genes (DEGs) with an FDR cutoff of 0.05 (**Figure 1C and Supplemental Table 2**), with many of the DEGs persisting even when performing differential expression analysis between *Gpr34* KO and WT mice within a given subtype of microglia (**Supplemental Table 2**). Among the DEGs from the analysis with cells from all microglial subtypes (**Figure 1C and 1D**), we observed upregulation of known DAM and inflammatory genes, such as *Anxa5*, *Axl*, *Itgax*, *Cst7*, *Cadm1*, and *Rftn1*. Gene set enrichment analyses (GSEA) of *Gpr34*-deficient microglia revealed transcriptional changes across immune, Alzheimer’s disease-related, myelination, cell death, and cholesterol metabolism related modules (**Figure 1E-F**). DAM-related gene sets (more specifically Amyloid-DAM gene set) based on previously published data^1,9^ (see Methods for details) were significantly enriched among upregulated genes of *Gpr34* KO microglia, including genes such as *Spp1* and *Cst7*, while “homeostatic” microglia gene sets were enriched among the downregulated genes (**Figure 1E**). Consistent with these changes, *Gpr34* KO microglia showed increased enrichment of immune and inflammatory gene sets among upregulated genes, particularly cytokine and chemokine signaling, complement activation, and responses to lipopolysaccharide and interferon gamma, along with modest enrichment of Alzheimer’s disease-related genes (**Figure 1E**). Functional modules associated with microglial activity such as response to lipids, myelination, cell killing, and GPCR pathways were also enriched among upregulated genes (**Figure 1F**), suggesting altered lipid handling and signaling capacity.

Quantitative analysis of subcluster proportions in the microglia single cell data (**Figure 1G**) revealed a slight but significant reduction of homeostatic microglia and a corresponding expansion of DAM microglia in *Gpr34* KO mice, with other subclasses not significantly changed.

Additionally, we used snRNA-seq to investigate the impacts of *Gpr34* loss on other cell types in the hippocampus (**Supplemental Figure 4A)**. Cell type composition analysis indicated a significant increase in the proportion of astrocytes that are disease-associated astrocytes (expressing previously defined markers genes such as *Osmr1*, *Ggta1*, and *Gsn*) and an increase (not reaching statistical significance) in the proportions of disease-associated microglia (DAMs; p-value=.083) and disease-associated oligodendrocytes (with previously reported markers such as *C4b* and *Serpina3n;* p-value=.29) (**Figure 1H**)^33–35^. Notably, these changes occur despite the absence of *Gpr34* expression in non-microglial cells, suggesting that the cell state changes in astrocytes and oligodendrocytes (ODCs) are secondary to the loss of *Gpr34* in microglia. No significant changes in neuronal abundance or other major cell classes were observed, though we were able to identify differentially expressed (DE) genes in numerous cell types, most notably *C4b* in ODCs (**Supplemental Figure 4B-G and Supplementary Table 3**). Note, unlike in the single-cell data, we do not see any DEGs in the microglia in the single-nucleus data. This is likely due to the rarity of microglia in the single nucleus data, leading to low power to detect DEGs^7,36^.

Together, the transcriptomics data demonstrate that the deletion of *Gpr34* results in a small but significant shift in microglia from a homeostatic state towards a more DAM-like and inflammatory state, correlated with an expansion of DAM proportions and accompanied by (presumably secondary) transcriptomic changes in astrocytes and ODCs. These findings suggest that GPR34 functions to maintain microglia in a homeostatic state *in vivo* and that loss of *Gpr34* may prime microglia toward reactivity, inflammation, and disease-like states.

### *Gpr34* KO microglia partially resemble microglia of the 5xFAD Alzheimer’s model in transcriptomic profile

To further evaluate the relationship between GPR34 loss and AD-related microglial transcriptomic profiles, we directly compared DE results of *Gpr34* KO vs WT microglia to those for the 5XFAD mouse model of AD vs WT at 6 months. 5xFAD is a well-established mouse model for amyloidosis that exhibits early and aggressive βθamyloid accumulation and robust microgliosis^31^. The original characterization of DAM and homeostatic transcriptional programs of microglia came from a study of the 5xFAD mouse line^9^. Changes in gene expression in *Gpr34* KO microglia were positively correlated with those observed in 5xFAD microglia. 26 of the 60 genes that were significantly altered in *Gpr34* KO vs WT microglia (**Figure 1C**) were also significantly altered in 5xFAD vs WT microglia, including many upregulated genes (*Cst7*, *Itgax*, *Anxa5*) and downregulated genes (including *Gimap1*, *Gimap6*, and *Gimap9*) all changing in the same direction (20 upregulated genes, 6 downregulated genes) (**Figure 2A**). Directly comparing the long fold changes (logFCs) from the DE analysis (**Figure 2B**) demonstrated positive correlation across the datasets (R = 0.53 among genes that reach significance in either comparison), particularly for DAMθassociated genes such as *Apoe, Spp1, Cst7,* and *Axl*. Although well correlated between *Gpr34* KO and 5xFAD, the effect size (log_2_FC) of the transcriptomic changes was generally larger for the DEGs in 5xFAD than in *Gpr34* KO microglia (**Figure 2B**). Since both DE gene analyses used the same set of WT samples as controls, overestimation of correlation values can occur. To address this, we applied a permutation-based approach which found the correlation to be significant, confirming that the observed correlation was not driven artifactually by shared controls. As expected, analysis of scRNAθseq from purified microglia showed a reduction in *Gpr34* expressio**n** in 5xFAD mice (**Figure 2C)**, reaching significance based on the raw p-values reported by EdgeR^37^ but not significant after applying fdrtools^38^ as described in the methods.

**Figure 2:**
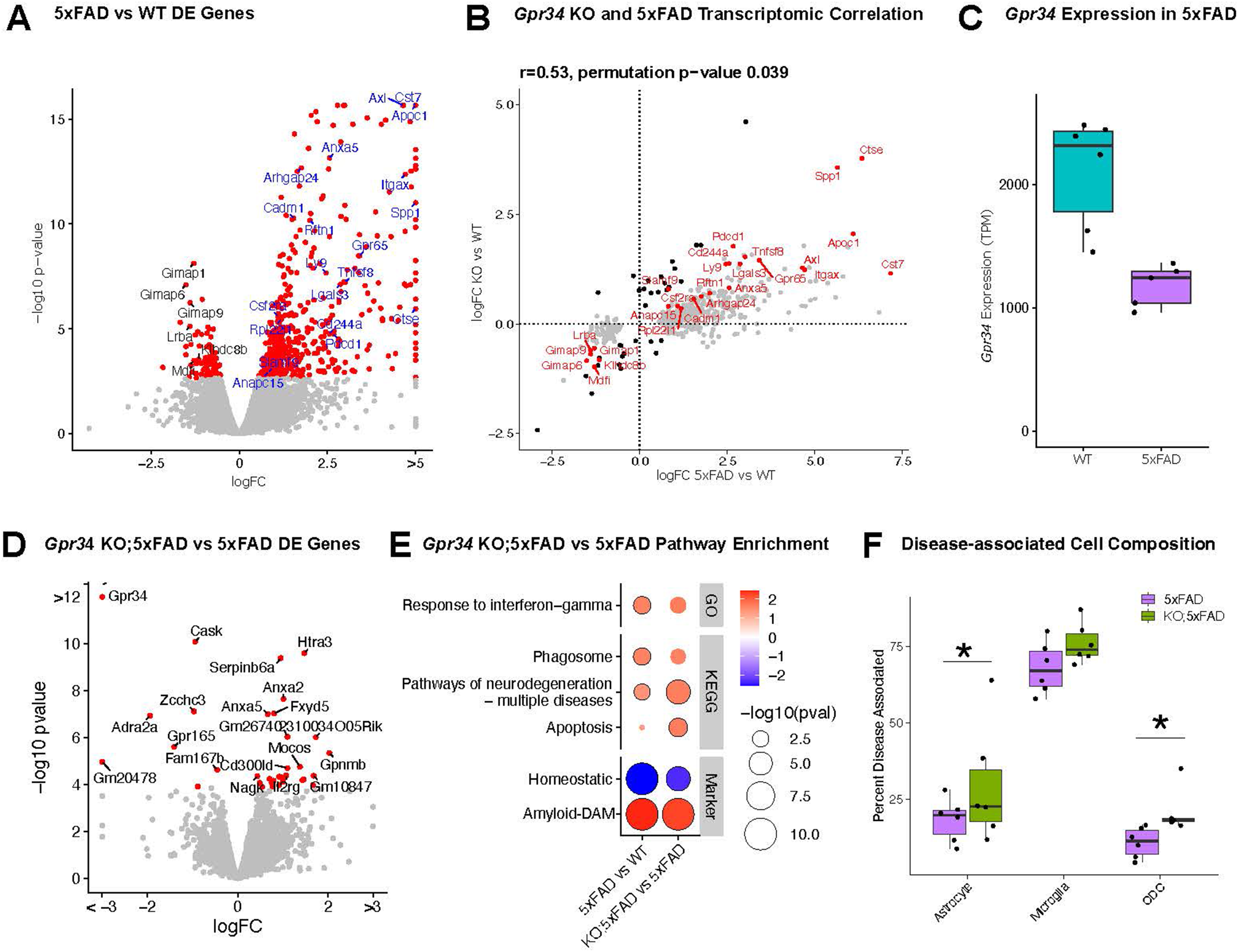
*Gpr34* deletion partially resembles 5xFAD Alzheimer’s model transcriptomic profile and potentiates DAM and inflammatory programs in the 5xFAD Alzheimer’s model. **A:** Volcano plot of DE genes for 5xFAD vs WT microglia in the single-cell data; those colored red are significant. Those gene names colored blue are upregulated in *Gpr34* KO vs WT mice and those in black are downregulated. **B:** Correlation between the logFC of genes in 5xFAD vs WT (x-axis) versus those in *Gpr34* KO vs WT (y-axis) in the single-cell data. We only show those genes that are significant in at least one comparison, the color of each dot represents which comparison each gene is significant in—black dots are significant in KO vs WT, grey dots in 5xFAD vs WT, and red are significant in both. Those that are significant in both are labelled. The correlation and the associated permutation-based p-value (the p-value is calculated so as to overcome the issue of shared controls which can lead to inflated correlations) are shown at the top. C: *Gpr34* expression for each mouse (y-axis, measured in TPM) in WT vs 5xFAD microglia (x-axis). **D:** Volcano plot of DE genes in KO;5xFAD vs 5xFAD mice in the single-cell data, calculated with a pseudobulk based method. Those colored red are significant. Those labelled genes are the top 20 most significant genes. **E:** Enrichment analysis of chosen terms related to disease, the immune system, and microglia function in the single cell data. The terms are on the y-axis, color is NES, size is –log_10_ pvalue, those outlined in black have FDR<0.1. The comparison is on the x-axis, either 5xFAD vs WT or KO;5xFAD vs 5xFAD. The dataset of origin for each term is annotated on the right side. **F:** Percent of disease associated cells (y-axis) in each *Gpr34* KO;5xFAD and 5xFAD mouse (color) for each cell type (x-axis) in the single-nucleus data. We see an increase in all of them, though it only reaches significance in the astrocytes and oligodendrocytes (FDR<.05), represented by the asterisks. **Sample sizes for sequencing analyses:** For single-cell RNA-seq, WT, n = 6 mice; *Gpr34* KO, n = 5 mice; and 5xFAD, n = 5 mice. For single-nucleus RNA-seq, n = 6 mice per condition.

### *Gpr34* deletion potentiates DAM and inflammatory programs in the 5xFAD Alzheimer’s model

Given that *Gpr34* KO by itself induces DAM-like and inflammation-related transcriptomic changes that partially overlap with that of 5xFAD microglia, we hypothesized that *Gpr34* KO would further exacerbate the microglial DAM phenotype in the 5xFAD mouse model of amyloidosis. To address this, we crossed *Gpr34* KO mice with 5xFAD mice and performed scRNA-seq of isolated microglia and snRNA-seq of hippocampal brain tissue from 6-month-old animals to assess gene expression and cell-type-specific changes across genotypes **(Figure 1A).**

Comparing *Gpr34* KO;5xFAD to *Gpr34* WT; 5xFAD by scRNA-seq revealed amplification of DAM and inflammatory-associated transcriptional responses in the *Gpr34*-deficient 5xFAD microglia. Differential expression analysis (**Figure 2D**) identified 27 DEGs that were upregulated in *Gpr34* KO;5xFAD microglia relative to 5xFAD microglia, including *Anxa5* and *Gpnmb*, reflecting an exacerbation of the DAM gene signature. There were only 8 significantly downregulated genes comparing *Gpr34* KO;5xFAD to *Gpr34* WT; 5xFAD microglia. Gene set enrichment analysis **(Figure 2E)** showed enrichment among upregulated genes of immune and microglial functional pathways, apoptosis-related terms, and gene sets associated with DAM and neurodegeneration, along with further downregulation of homeostatic gene sets. Cell-type composition analysis showed an increased proportion of DAMs in *Gpr34* KO;5xFAD microglia relative to 5xFAD, though this did not reach significance (**Supplemental Figure 5A**).

snRNAseq of hippocampal tissues (**Figure 2F**) confirmed that *Gpr34* deletion in the 5xFAD model significantly increased the proportion of disease-associated astrocytes and disease-associated oligodendrocytes with a non-significant increase in DAMs^33–35^. As with the *Gpr34* KO vs WT analysis, we were able to identify many DEGs in various cell types other than microglia (**Supplemental Figure 5B-F and Supplemental Table 3**).

Taken together, *Gpr34* loss alone induces DAM and inflammation signatures in microglia (**Figure 1**) and its deletion in 5xFAD mice further amplifies, rather than reverses, the DAM and inflammatory transcriptomic profiles of microglia in 6-month-old 5xFAD mice (**Figure 2**). These findings suggest that GPR34 plays a modulatory role, restraining disease-associated and inflammatory pathways in response to βθamyloid pathology.

### *Gpr34* loss exacerbates microglial reactivity and increases amyloid beta engulfment in 5xFAD mice

We examined the effects of *Gpr34* knockout on other aspects of AD pathology (amyloid-beta (Aβ) and glial reactivity) by immunohistochemistry (IHC) of cortical sections of 6-month-old 5xFAD mice with or without *Gpr34* (*Gpr34* KO;5xFAD). Using confocal z-stack imaging and Imaris 3D surface analysis, we observed a non-significant increase in total Aβ-positive burden in the cortical ROIs in *Gpr34* KO;5xFAD mice compared to 5xFAD (**Figure 3A and 3B**). The total Iba1+ immunoreactivity within cortical ROIs was unaltered between these genotypes (**Figure 3C**). In addition, we analyzed Iba1+ immunoreactivity in the same cortical regions in WT and *Gpr34* KO mice and did not observe significant differences (data not shown). However, we found that the CD68 signal (a lysosome marker) within Iba1+ microglial compartments, normalized to total Iba1 signal, was significantly increased in *Gpr34* KO;5xFAD mice (**Figure 3D-E)**. Notably, Aβ signal within CD68+ structures (phagosomes/lysosomes) inside Iba1+ microglia was significantly increased in *Gpr34* KO;5xFAD mice (**Figure 3F**). We also measured GFAP (a marker of astrocytes, whose expression increases upon astrocyte activation) in the cortex whole-slide immunostaining and 2D image analysis. We found a 45.7% increase in GFAP+ area in *Gpr34* KO;5xFAD mice relative to 5xFAD, although that did not reach significance (p =0.08) (**Figure 3G and 3H**). Together, these immunohistochemical findings align with our transcriptomic results and suggest that *Gpr34* loss in 5xFAD mice enhances microglial reactivity, amyloid beta engulfment, with a trend towards increased astrocyte reactivity.

**Figure 3:**
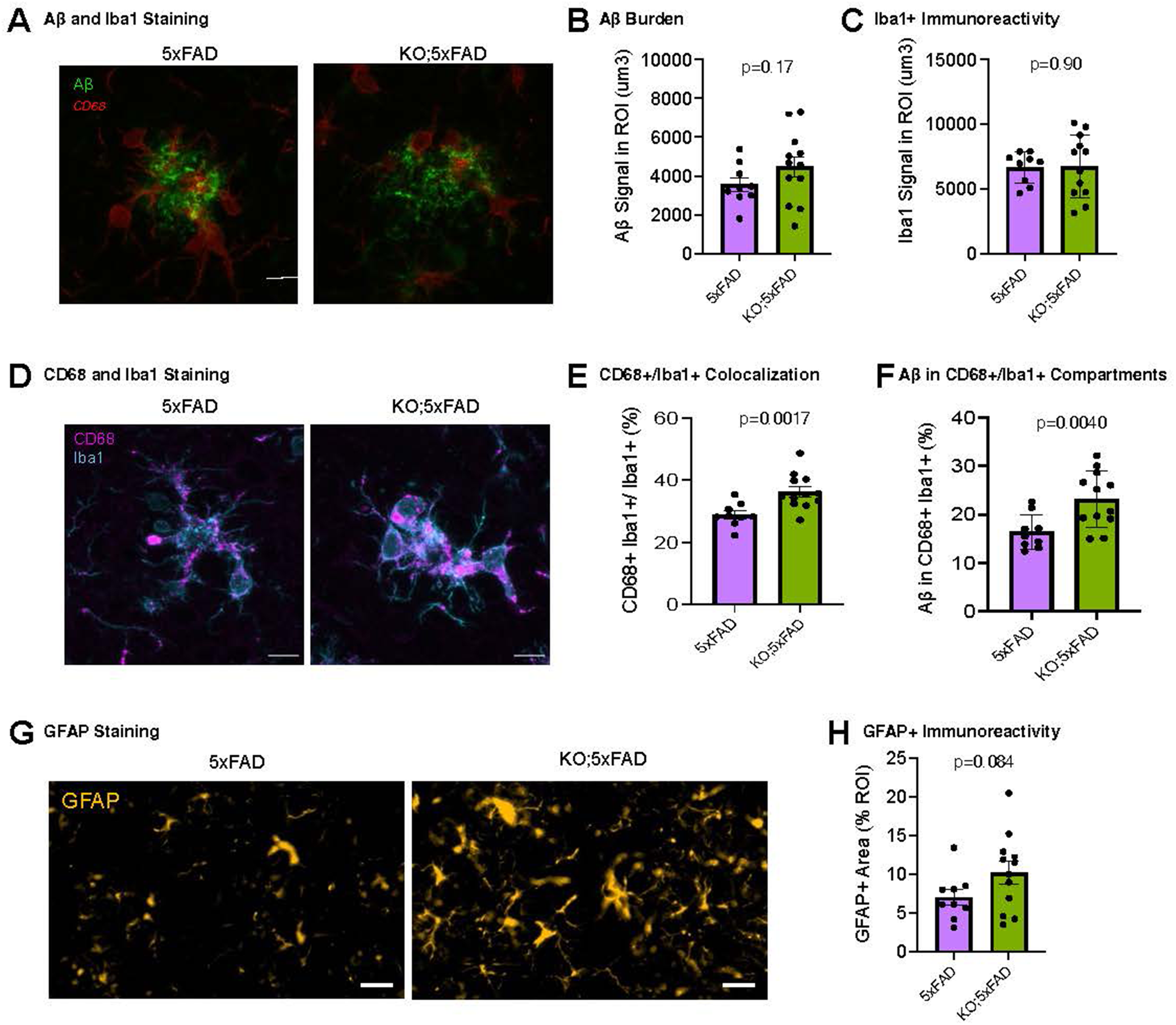
*Gpr34* deletion exacerbates microglial reactivity and increases amyloid beta engulfment in 5xFAD mice. **A:** Representative image of Aβ (shown in green) and Iba1 (shown in red) immunofluorescence staining in the cortex of 5xFAD and *Gpr34* KO;5xFAD mice. Scale bar indicates 10µm. **B:** Quantification of total Aβ signa in the region of interest (ROI) of the cortex of 5xFAD and *Gpr34* KO;5xFAD mice using confocal z-stack imaging and Imaris 3D surface analysis. Values represent summed Aβ-positive signal within each ROI. P = 0.17. **C:** Quantification of total Iba1 immunoreactivity in the ROI of the cortex of 5xFAD and *Gpr34* KO;5xFAD mice using confocal z-stack imaging and Imaris 3D surface analysis. P = 0.90. **G:** Representative image of Iba1 (shown in cyan) and CD68 (shown in magenta) immunofluorescence staining in the cortex of 5xFAD and *Gpr34* KO;5xFAD mice. Scale bar indicates 10µm. **D:** Representative image of CD68 (shown in pink) and Iba1 (shown in blue) immunofluorescence staining in the cortex of 5xFAD and *Gpr34* KO;5xFAD mice. Scale bar indicates 10µm. **E.** Quantification of CD68 signal within Iba1+ microglial compartments, normalized to total Iba1 signal, in cortical ROIs from 5xFAD and *Gpr34* KO;5xFAD mice using Imaris 3D surface analysis. Values are normalized to total Iba1 signal. P = 0.0017. **F:** Quantification of Aβ signal within CD68+/Iba1+ compartments in cortical ROIs using Imaris 3D surface analysis. Values are normalized to total Aβ signal. P = 0.0040. **G:** Representative whole-slide immunofluorescence image of GFAP (shown in yellow) staining in the cortex of 5xFAD and *Gpr34* KO;5xFAD mice. Scale bar indicates 20µm. **H:** Quantification of GFAP-positive immunoreactivity within cortical ROIs from 5xFAD and *Gpr34* KO;5xFAD mice, using 2D whole-slide image analysis, measured as percent area. P = 0.084. **Imaging quantification:** 5xFAD, N=9 mice and *Gpr34* KO;5xFAD, N=12 mice. 4-5 brain slices were analyzed for each animal for Aβ, Iba1 and CD68 measurements. 2-3 brain slices were analyzed for each animal for GFAP measurement. Data represent mean ±s.e.m. Welch’s t test. **Sample sizes for sequencing analyses:** For single-cell RNA-seq, 5xFAD, n = 5 mice, and *Gpr34* KO;5xFAD, n = 6 mice. For single-nucleus RNA-seq, n = 6 mice per condition.

### Generation and pharmacological validation of *GPR34*-deficient human iPSC-derived microglia (iMGLs)

We next sought to establish a robust platform for mechanistic and pharmacological studies at the cellular level. However, cultured primary murine microglia were inappropriate for these purposes, as *Gpr34* expression is swiftly downregulated upon removal from the CNS^12,39^(data not shown). We therefore utilized human iPSC-derived microglia-like cells (iMGLs), following a previously established 40-day cytokine-based protocol^32,40^. We used a CRISPR/Cas9 strategy to introduce precise 2-base-pair deletions into the *GPR34* coding sequence of WTC-11 iPSCs and confirmed an indel mutation in two different *GPR34* KO clones (designated KO-1 and KO-2) by Sanger sequencing and RNA-sequencing (**Figure 4A; Supplemental Figure 6A).** For each KO clone, a matched isogenic control clone (WT-1 and WT-2) that underwent the identical CRISPR editing and clonal expansion workflow but retained the wild-type *GPR34* sequence was used as a paired control. After differentiation into iMGLs, both KO clones responded similarly to a known GPR34 agonist, Compound 4B (**Supplemental Figure 9A**)^41^, so we focused on one pair (KO-1 and WT-1) for subsequent experiments. *GPR34* KO-1 iMGLs displayed ∼2-fold elevated *GPR34* transcript level relative to WT-1 iMGLs (**Supplemental Figure 6B).** However, quantitative mass spectrometry showed that *GPR34* KO-1 iMGLs indeed lacked detectable GPR34 protein (**Figure 4B**).

**Figure 4:**
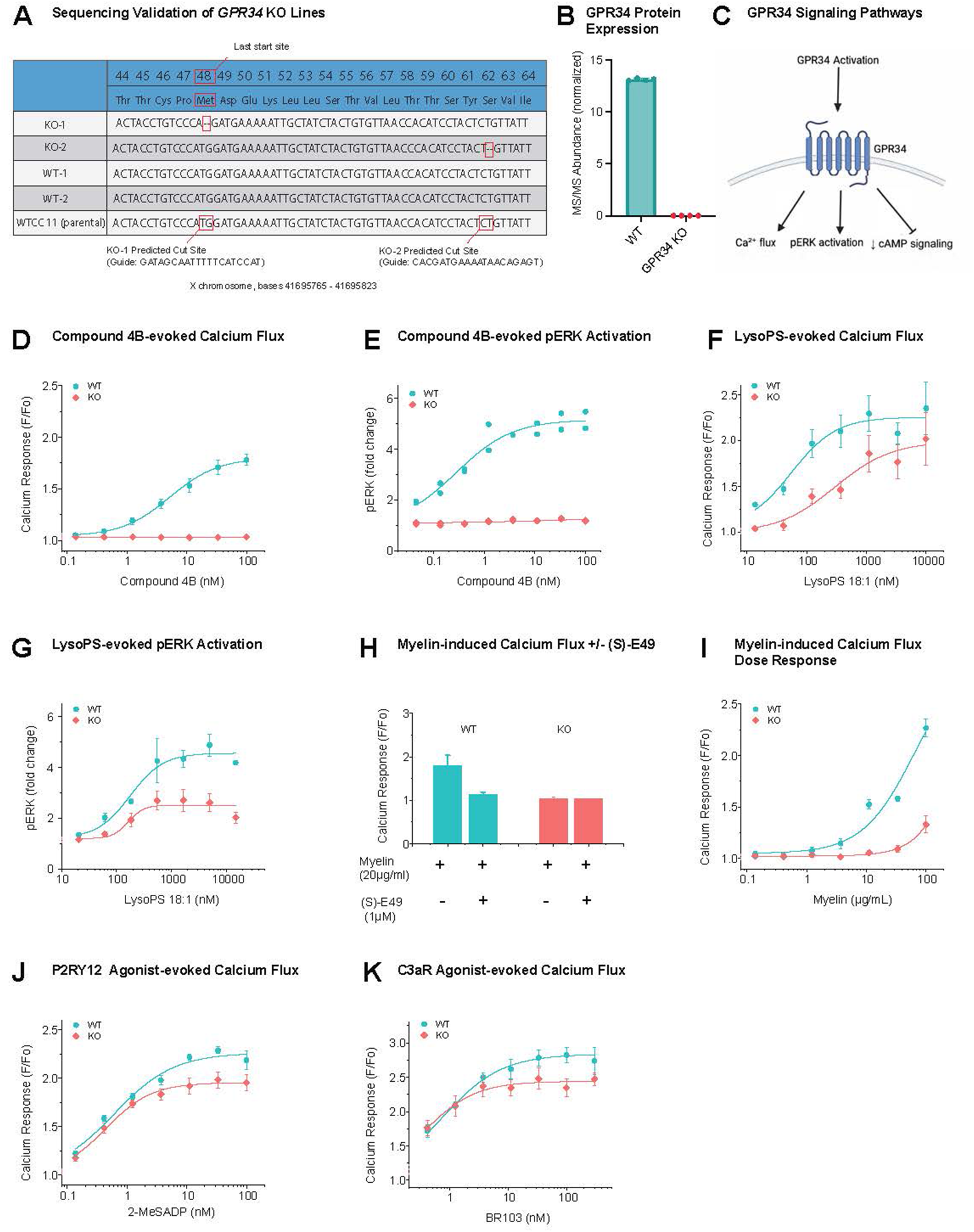
Generation and pharmacological validation of *GPR34*-deficient human iPSC-derived microglia (iMGLs) **A:** Sequencing validation of two independent *GPR34* iPSC knockout (KO) clones (KO-1 and KO-2) and their respective WT controls. Guide RNA sequences for KO-1 and KO-2 are indicated below the alignment, with predicted Cas9 cut sites marked. Red boxes indicate cut sites, confirming editing of *GPR34* in the KO clones relative to WT and parental controls **B:** MS/MS analysis shows the loss of GPR34 protein in *GPR34* KO-1 iMGLs **C:** Schematic illustrating GPR34 activation and downstream signaling pathways. **D:** Dose-dependent calcium response to the GPR34 agonist Compound 4B in WT and *GPR34* KO iMGLs. **E:** Dose-dependent pERK activation in response to the GPR34 agonist Compound 4B in WT and *GPR34* KO iMGLs. **F:** Dose-dependent calcium response to the GPR34 endogenous ligand lysoPS in WT and *GPR34* KO iMGLs. The representative curve was shown from the average data from triplicates. **G:** Dose-dependent pERK activation in response to the GPR34 endogenous ligand lysoPS in WT and *GPR34* KO iMGLs. **H:** Myelin-induced calcium signaling in WT and *GPR34* KO iMGLs treated with (S)-E49. **I:** Dose-response of myelin-induced calcium flux in WT and *GPR34* KO iMGLs. **J-K:** Dose-dependent calcium response to GPR34 unrelated agonists (P2Y12/2-MeSADP, C3aR/BR103) in WT and *GPR34* KO iMGLs. **Sample sizes and stats:** *GPR34* KO line 1 (KO-1) and its control (WT-1) were used for this study. The representative curve for D, F, G, H and I was shown from the average data are from triplicates and for E is from duplicates.

To evaluate any gross morphological effects of *GPR34* loss, we performed immunostaining on iMGLs that were dissociated on day 42 and replated at equal cell density. Nuclear staining confirmed comparable cell numbers across all wells and genotypes (**Supplemental Figures 7A-B)**. Cell area was measured by immunostaining using a panel of cell-surface and intracellular microglial markers consisting of: P2RY12, IBA1, C1q, C3aR, and CD45. All markers exhibited robust staining, validating that these were fully differentiated iMGLs^42,43^ (**Supplemental Figure 7A**). *GPR34* KO-1 iMGLs showed similar immunostaining intensities for all tested markers and no obvious differences in cell morphology were observed between KO and WT iMGLs (**Supplemental Figures 7C-D).** Additionally, flow cytometry conducted on Day 40, upon completion of the microglial differentiation protocol, showed comparable expression of CD45, CD11b, P2RY12, and CX3CR1 microglial markers in WT-1 and *GPR34* KO-1 cells. (**Supplemental Figures 8A-B**).

To determine the functional consequences of *GPR34* loss on downstream pathways and ligand responses, we used a set of pharmacologically validated tool compounds in our iMGL platform and examined downstream GPCR signaling pathways associated with GPR34^44^. **(Figure 4C)**. GPR34 activation by a validated agonist, Compound 4B, elevated intracellular calcium levels with an EC_50_ of approximately 3 nM (**Figure 4D and Supplemental Figure 9A**) and increased ERK1/2 phosphorylation with an EC_50_ of approximately 0.5 nM (**Figure 4E**). Both responses were abolished in *GPR34* knockout iMGLs (**Figure 4D-E**) and by the GPR34 antagonist YL-365^45^. (**Supplement Figure 9B-C**). GPR34 is thought to be a G*α*i-coupled receptor^44^. However, upon treatment with Compound 4B, the level of forskolin-induced cAMP was unchanged, suggesting that GPR34 does not significantly modulate cAMP signaling under these conditions (**Supplemental Figure 9D**). These data validate our iMGL platform for assessing GPCR signaling, demonstrating robust and receptor-dependent activation of canonical downstream signaling pathways.

Lysophosphatidylserine (LysoPS), which has been proposed as an endogenous GPR34 ligand, stimulated calcium and pERK responses in wildtype iMGLs^16^. Responses to LysoPS were reduced in the GPR34 knockout iMGLs, albeit incompletely and with a rightward shift in the dose response curve (**Figure 4F-G**), suggesting the presence of additional LysoPS-responsive receptors in iMGLs besides GPR34. Myelin debris, which is lysoPS-rich, has recently been reported to promote microglial dysfunction, neuroinflammation, and AD-related pathology via GPR34^18,21^. Consistent with this, we found that myelin also induced calcium elevations in microglia, which was blocked in *GPR34* KO-1 iMGLs (**Figure 4H-I**), or by a GPR34 antagonist, (S)-E49 (**Figure 4H**) ((S)-E49 - Itoh et. al. Patent application publication US2010/0130737). By contrast, calcium signals elicited by agonists of other microglia GPCRs (P2Y12 and C3aR), remained largely intact in *GPR34* KO-1 iMGLs (**Figure 4J-K**).

Together, these findings show that GPR34 primarily regulates calcium and pERK signaling but not cAMP in human iPSC-derived microglia and that LysoPS, the proposed endogenous ligand, acts, at least in part, in a GPR34-dependent manner. Importantly, myelin debris also stimulates calcium signaling in microglia in a GPR34-dependent manner.

### *GPR34* deficiency selectively impairs myelin phagocytosis in iPSC-derived microglia

Phagocytosis is a core microglial function essential for homeostatic maintenance of the central nervous system (CNS)^2,5,6^. Our data indicate that myelin, a key *in vivo* phagocytic substrate, can signal through GPR34 (**See Figure 4H-I**), leading us to examine whether GPR34’s interaction with myelin also regulates its engulfment. To directly investigate the role of GPR34 in this fundamental process, we performed *in vitro* phagocytosis assays using iMGLs from WT-1 and *GPR34* KO-1 iMGLs, along with the GPR34 antagonist YL-365^45^. These experiments build on prior findings suggesting that GPR34 contributes to microglial phagocytosis *in vivo*^19^ and substrate selectivity *in vitro*^23^.

We assessed the phagocytic uptake of three substrates: myelin debris, amyloid fibrils, and *E. coli* (**Figure 5A**, schematic). Each substrate was conjugated to pHrodo Deep Red, a pH-sensitive fluorophore that becomes fluorescent in acidic compartments such as the phagolysosome. After exposure of WT-1 and *GPR34* KO-1 iMGLs to each pHrodo-labeled substrate, we measured median fluorescence intensity (MFI) using two complementary approaches: live confocal imaging 5 and 24 hours after substrate addition (**Figure 5A**), and flow cytometry at the 25-hour terminal time point (**Supplemental Figure 10**). Both methods showed a significant 30% reduction of myelin uptake in *GPR34* KO-1 microglia compared to WT-1 cells (95% CI: 12-49%) (**Figures 5B and Supplemental Figures 10A-B**). In contrast, phagocytosis of amyloid fibrils and *E. coli* bioparticles was not significantly affected (**Figure 5C-D and Supplemental Figures 10C-D**). To test whether pharmacological inhibition of GPR34 could replicate this selective impairment, iMGLs were pre-treated with the GPR34 antagonist YL-365 (10 µM) for 30 minutes prior to substrate exposure. Antagonist-treated cells showed a similar reduction in myelin phagocytosis, with no effect on uptake of the other substrates (**Figures 5E and Supplemental Figures 10E-H**). Together, these results indicate that, at least *in vitro*, GPR34 plays a selective and critical role in microglial phagocytosis of myelin debris. However, it is unclear whether the reduced myelin content in *GPR34* KO microglia reflects reduced engulfment or increased degradation of myelin, and future studies are needed to distinguish these possibilities (please also see the Public Review Comment #3 for additional discussion).

**Figure 5:**
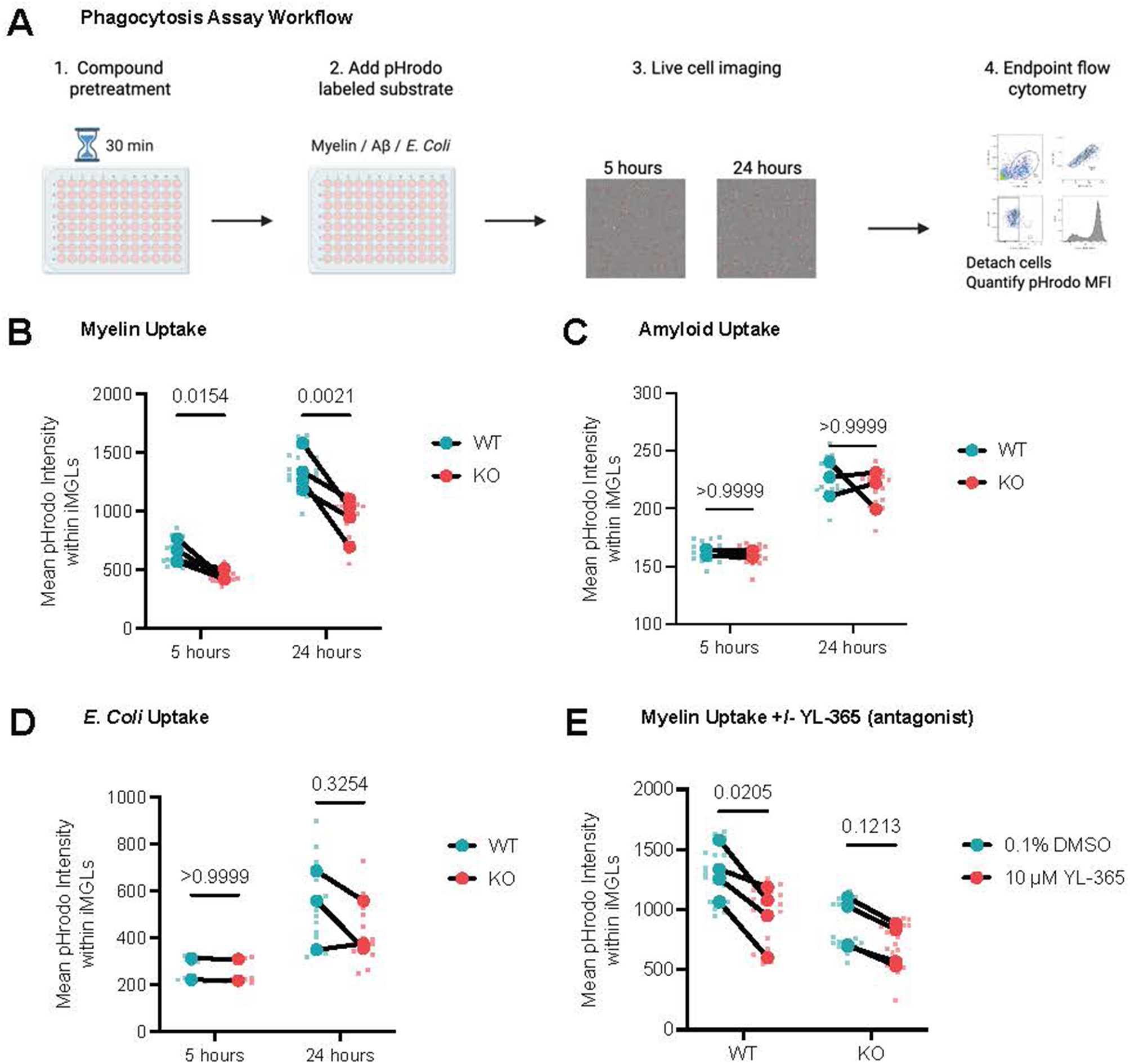
*GPR34* deficiency selectively impairs myelin phagocytosis in iPSC-derived microglia. **A:** Experimental workflow for live-cell imaging and flow cytometry-based assessment of phagocytosis. iMGLs were incubated with pHrodo-labeled substrates and imaged following 5 and 24 hours of substrate incubation using the Opera Phenix live-imaging system. Cells were then detached from imaging plates onto PhenoPlates and processed by flow cytometry on the CytoFLEX LX for assessment of median fluorescence intensity (MFI). **B-D:** WT and *GPR34* KO iMGLs were exposed to pHrodo-labeled myelin (**B)**, amyloid fibrils (**C)**, or *E. coli* bioparticles (**D)** and assessed by live cell imaging at 5 and 24 hours. Lines connect paired WT and KO biological replicates derived from the same experimental batch. **E:** WT and *GPR34* KO iMGLs were exposed to GPR34 antagonist YL-365 (10 µM) and assessed for phagocytosis of pHrodo labeled myelin at 24 hours by live cell imaging. **Sample sizes and stats:** Data are represented as mean values with +/- SEM. Smaller circles represent technical replicates, and larger circles represent biological replicates. Statistical tests were undertaken at 5h, 24h, and 25h (n=2-4 independent biological replicates with 5-6 technical replicates per condition). Two-way ANOVA with Bonferroni correction, multiple comparisons test, paired t-test with Welch’s correction, as appropriate.

### Multiomic analysis of *GPR34*-deficient iMGLs highlights dysregulation of immune, replicative, and lysosomal programs

To investigate the molecular consequences of GPR34 loss in human iMGLs, we performed bulk RNA sequencing and quantitative mass spectrometry (MS) proteomics on WT-1 versus *GPR34* KO-1 iMGLs. Principal component analysis (PCA) of RNA-seq data demonstrated clear separation of WT-1 and KO-1 iMGLs on PC1 across multiple rounds of differentiation **(Supplemental Figure 11A).**

Differential expression analysis revealed broad transcriptional differences between *GPR34* KO-1 and WT-1 iMGLs (**Figure 6A, Supplement Table 4**). Gene set enrichment analysis (GSEA) highlighted enrichment of DAM-related genes, and of pathways related to phagocytosis, endocytosis, and lysosomal function among the upregulated genes, consistent with increased expression of genes such as *PLIN2, CD36, LAMP2 and SYT11*^46–49^. Conversely, pathways associated with DNA replication, cell cycle, and proliferation were downregulated with genes such as *MCM4*, and *TOP2A*, and *CBX5* contributing to these changes^50–52^ (**Figure 6B-C**).

**Figure 6:**
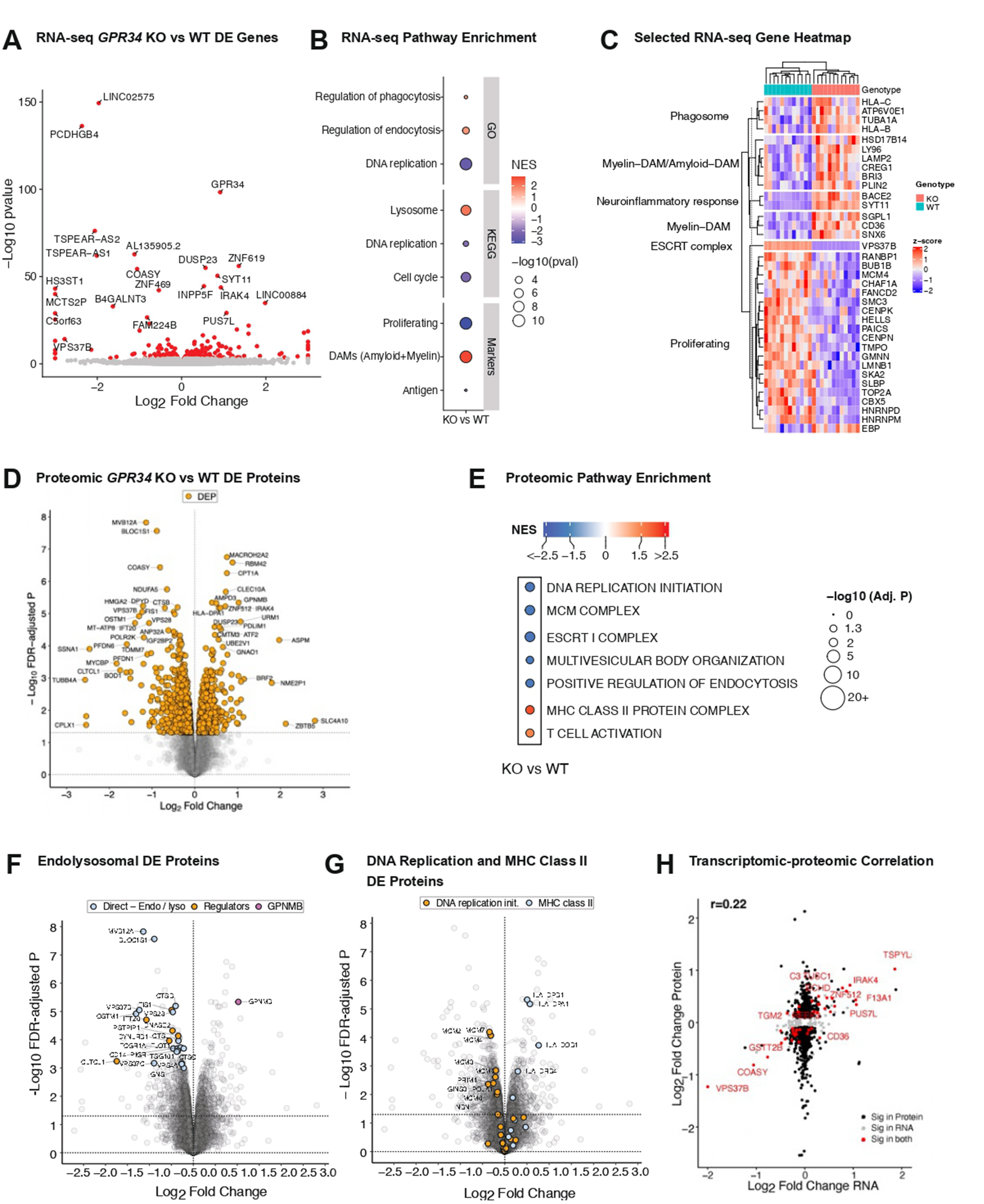
Multiomic analysis of *GPR34*-deficient iMGLs highlights dysregulation of immune, replicative, and lysosomal programs. **A:** Volcano plot of RNA-seq analysis in *GPR34* KO vs WT iMGL samples. Red dots are significant, top genes (those with smallest p-values) are labelled. Based on the meta-analysis of two experimental batches. **B:** Plot of selected gene sets (y-axis) from gene set enrichment analysis for the *GPR34* KO vs WT RNA-Seq comparison. Color is NES, size is -log_10_ p-value, those circled in black are significant (FDR<.1). **C:** Heatmap of selected genes from pathways altered in *GPR34* KO vs WT iMGL samples. The columns are samples, rows are genes, the color is the z score of the TPM expression values. The bars on the top represent KO vs WT. Note the separation between genes from different gene sets (y-axis). **D:** Volcano plot of differentially expressed proteins (DEPs) in MS/MS quantitative proteomics analysis of *GPR34* KO vs WT iMGLs. Top DEPs are labelled. **E:** Gene set enrichment analysis of quantitative proteomics data highlighting prominently changed molecular pathways in *GPR34* KO vs WT iMGLs. Color indicates normalized enrichment scores (NES) and bubble size indicates FDR-adjusted *P*-value. All shown pathways are significantly altered (FDR-adjusted P < 0.05) in KO iMGLs (compared to WT). **F:** Volcano plot of quantitative proteomics analysis of *GPR34* KO vs. WT iMGLs highlighting endosomal and lysosomal pathways. Proteins directly involved in endosome/lysosome function (“direct - endo/lyso”) and known regulators of these pathways (“regulators”) are indicated. GPNMB, a putative biomarker of lysosomal dysfunction, is also highlighted. **G:** Volcano plot of quantitative proteomics analysis of *GPR34* KO vs WT iMGLs highlighting DNA replication and MHC class II pathways **H:** Comparison of transcriptomic (x-axis) and proteomic (y-axis) changes (log_2_FC) in *GPR34* KO vs WT iMGLs. Color indicates DE in RNA, protein, or both. Only genes that are DE in either or both assays are shown.

Consistent with the transcriptomic analysis, PCA analysis of the proteomics dataset demonstrated distinct clustering of WT-1 and KO-1 samples (**Supplemental Figure 11B**). *GPR34* deletion produced a pronounced effect on the iMGL proteome, with thousands of differentially expressed proteins (DEPs) surpassing the significance threshold (FDR-adjusted *P* < 0.05) (**Figure 6D**). GSEA indicated enrichment of proteins related to T-cell activation and MHC Class II complex formation (FDR adjusted P < 0.05) among the upregulated DEPs, whereas pathways related to the MCM complex, DNA replication initiation, multivesicular body organization, endocytic regulation, and ESCRT I complex were downregulated (**Figure 6E)**.

Within the vesicular trafficking pathways, VPS37B, a core ESCRT component required for endosomal sorting^53^, was strongly downregulated at both the transcript and protein levels (log_2_FC = -2.77, adjusted p = 6.8 x10^-15^ for RNA; log_2_FC = -1.23, adjusted p = 8.9 x10^-6^ for protein) **(Figure 6A**, **Figure 6D)**. The broader proteomic landscape also emphasized endosomal and lysosomal dysfunction, evidenced by reduction of ESCRT complex components (MVB12A, VPS37B, VPS37C, VPS28), lysosomal enzymes (CTSL, CTSB, CTSC, GNS), lysosomal membrane proteins (OSTM1), and regulators of endosomal trafficking (BLOC1S1, FLOT1, VPS13C, DYNLRB1, CLTCL1) **(Figure 6F).** Notably, these reductions were accompanied by a striking increase in GPNMB at the protein level, a putative biomarker of lysosomal impairment, and a canonical DAM marker^54,55^ (**Figure 6F**). Together, these data highlight vesicular trafficking and lysosomal biology as microglial processes likely disturbed by GPR34 loss of function.

Consistent with the transcriptional downregulation of proliferative pathways in RNAseq data, we observed reduced abundance of DNA replication proteins, including multiple MCM family members (MCM2-7) in *GPR34* KO-1 iMGLs (**Figure 6G**). These molecular phenotypes suggest reduced proliferation of *GPR34*-deficient cells, which is supported by our findings that KO-1 cultures yielded fewer cells than WT-1 during the iMGL differentiation protocol (**Supplemental Figure 11C)**. In parallel, antigen presentation pathways were strongly upregulated, driven by increased protein levels of MHC class II molecules such as HLA-DPB1, HLA-DPA1, HLA-DQB1, and HLA-DRB4, highlighting a shift from proliferative to immune-related programs^56–58^.

Finally, integration of transcriptomic and proteomic datasets demonstrated significant albeit modest correlation between respective log_2_FCs of RNAs and proteins (r = 0.22, *P*=2.96e-17) (**Figure 6H**). In general, the magnitude of change was much greater on the proteome than on the transcriptome, implying that many of the changes driven by loss of GPR34 occur through post-transcriptional mechanisms, which suggest that proteomics can detect biologically relevant changes that are not present at the mRNA level. Nonetheless, several prominently altered DEGs exhibited directionally consistent and pronounced changes in RNA and protein levels, including downregulation of VPS37B, a core ESCRT-I component required for lysosomal degradation, and COASY, which encodes coenzyme A synthase^59^.

Together, our omics data in human iMGLs indicate that loss of *GPR34* impacts broadly microglial RNA and protein expression, with upregulation of pathways related to immune activation, and suppression of pathways related to DNA replication, phagocytosis, lysosomes and vesicle trafficking.

### *GPR34* loss blunts microglial transcriptional responses to myelin challenge

Myelin is a potent stimulus for microglia, inducing DAM and inflammatory gene expression signatures^32^. Since *GPR34*-deficient microglia show impaired calcium signaling in response to myelin (**Figure 4H-I**) and selective deficits in myelin phagocytosis (**Figure 5B-E**), we next investigated whether GPR34 alters microglial transcriptional responses to myelin. To capture both early and sustained responses, WT-1 and *GPR34* KO-1 iMGLs were exposed to myelin in culture for 2 hours or 24 hours, followed by bulk RNA sequencing.

Principal component analysis (PCA) revealed that the transcriptome of 24-hour myelin-treated samples clustered distinctly from the 2-hour and untreated (no-myelin) groups, reflecting transcriptional changes induced by prolonged myelin exposure (**Figure 7A**). The 2-hour and no-myelin samples clustered more closely, consistent with modest early transcriptional responses. In all treatment conditions, WT-1 and KO-1 samples clustered separately from each other, in line with our previous analyses of *GPR34* KO-1 microglia (**Figure 7A**). Differential expression analysis and transcriptome-wide impact (TWI) scores^60^ from KO-1 vs WT-1 iMGL samples confirmed that myelin induced a substantial transcriptomic shift, with KO-1 samples showing attenuated responses at both time points (231 DEGs in KO-1 vs 921 DEGs in WT-1 at 2 hours, 5648 DE genes in KO-1 vs 8370 genes in WT-1 at 24 hours, (**Figure 7B and Supplemental Figure 12A**). Despite fewer DEGs in KO-1 samples treated by myelin, many of the DEGs at 24 hours were shared with WT-1 iMGLs treated with myelin (4,729 shared DEGs, r = 0.89, based on genes DE at an FDR cutoff of 0.05 in at least one of the two comparisons) (**Figure 7C**; **Supplemental Figure 12B).** Overall, the loss of GPR34 attenuated the broader transcriptomic response induced by prolonged myelin exposure but did not abolish or profoundly alter the nature of the response, implying that additional myelin-responsive pathways act alongside GPR34. This contrasts with acute calcium signaling response to myelin, which appears to be largely mediated by GPR34 (**Figure 4H-I**).

**Figure 7:**
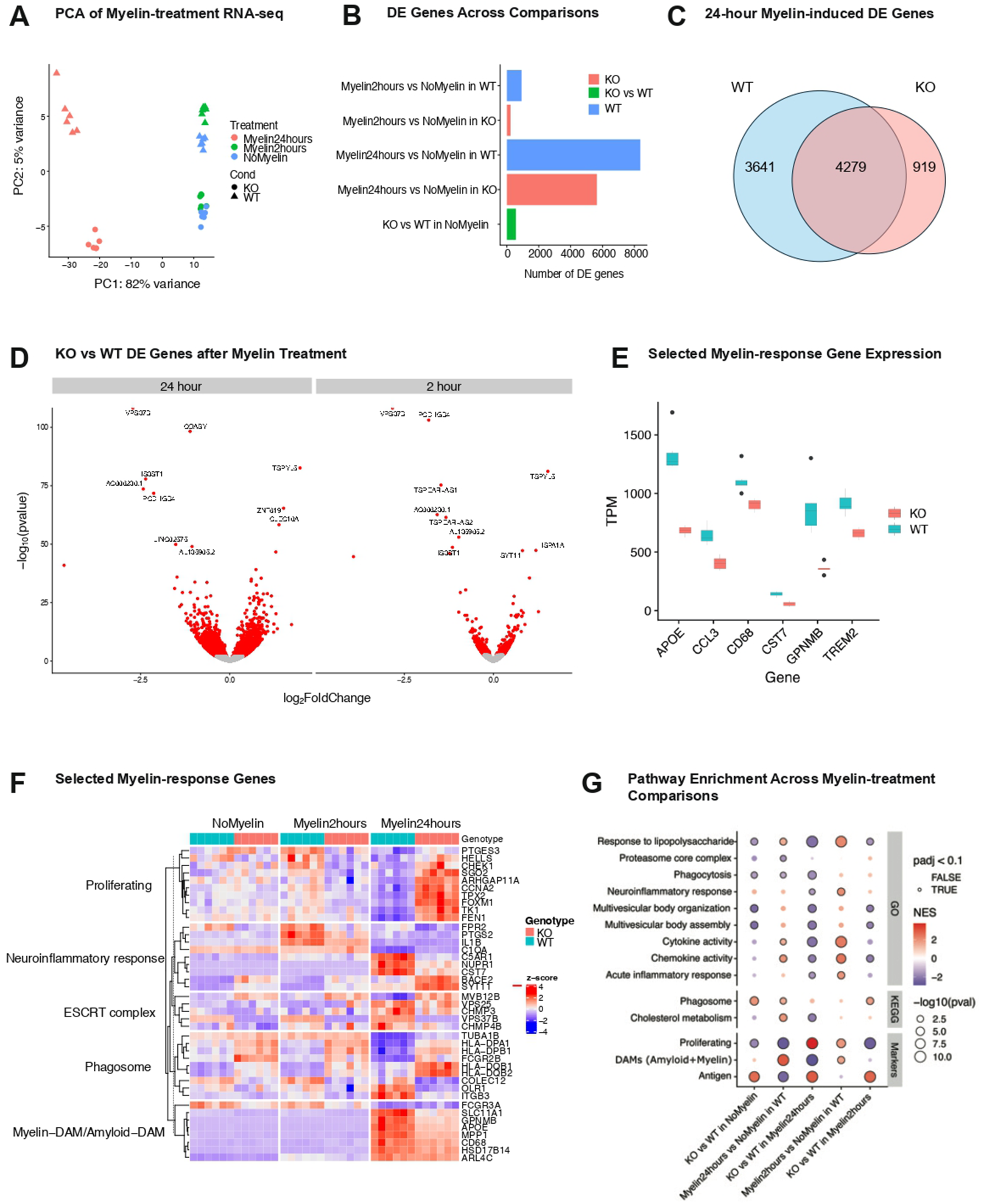
*GPR34* loss blunts microglial transcriptional responses to myelin challenge. **A:** PCA plot of iMGL bulk RNA-seq samples, colored by myelin treatment. The genotype is represented by the shape. **B:** Bar plots of the number of DE genes (x-axis) for each of the myelin-treatment comparisons (y-axis). **C:** Venn diagram of the number of DE genes after 24-hour myelin treatment versus no-myelin in WT and *GPR34* KO iMGLs. Note overlaps are only counted if they have the same direction of effect. **D:** Volcano plots comparing *GPR34* KO versus WT iMGLs after 24-hour or 2-hour myelin treatment. Significant hits are colored red (FDR<.05), genes with the smallest p-values are labelled. **E:** Expression of selected microglial response genes *APOE*, *CCL3*, *CD68*, *CST7*, *GPNMB*, *TREM2* (TPM, y-axis) in WT and *GPR34* KO iMGLs (x-axis) after 24-hour myelin treatment. In the box plots the center line is the median, the boxes span from 25% to 75%, the length of the whiskers is based off the smallest/largest value within 1.5 of the interquartile range from the median, and everything outside that range is plotted as an outlier. **F:** Heatmap of selected myelin-response genes across genotypes and myelin-treatment conditions. The columns are samples, rows are genes, the color is the z score of the TPM expression values. The bars on the top represent KO vs WT. Note the separation between samples with different treatments (x-axis) and genes from different gene sets (y-axis). **G:** Gene set enrichment analysis across genotypes and myelin-treatment comparisons. The terms are on the y-axis, color is NES, size is –log_10_ p-value, those outlined in black have FDR<0.1. The comparison is on the x-axis. The dataset of origin for each term is annotated on the right side.

At 2 hours, myelin induced a significant transcriptional response in WT-1 iMGLs with roughly 1,500 DEGs, characterized by rapid upregulation of immediate-early (IEGs) and cytokine-responsive genes such as *TNF, DUSP1*, and *FOS* consistent with acute activation of microglia (**Supplemental Figure 12C**). Caution must always be used with IEGs since such genes are known to also be related to certain activation artifacts, as has been reported previously^61^, By 24 hours, the transcriptional response to myelin expanded to roughly 8,000 DEGs and shifted toward lipid-related gene sets. Specifically, myelin exposure triggered downregulation of cholesterol biosynthesis genes (*HMGCS1, MVK, FDFT1, DHCR24*), upregulation of LDLR negative regulator (*MYLIP*), downregulation of cholesterol uptake gene (*LDLR*), and upregulation of cholesterol export genes (*ABCG1* and *PLTP*), collectively suggesting changes in handling and export of cholesterol (itself a component of myelin)^62–65^ **(Supplemental Figure 12C)**.

To address whether *GPR34* deficiency alters the microglial response to myelin, we compared the transcriptional changes between WT-1 and KO-1 iMGLs. At 24 hours following myelin treatment, *VPS37B* (a core ESCRT-I component required for lysosomal degradation) and *COASY* (which catalyzes the final steps of coenzyme A biosynthesis) were both reduced in KO-1 iMGLs relative to WT-1 samples (**Figure 7D**). Relative to untreated controls, VPS37B was upregulated by myelin in both genotypes. In contrast, *COASY* was downregulated in WT-1s after myelin treatment but was not significantly different in KO-1 cells. These changes suggest impaired vesicle trafficking and altered metabolic-related programs in KO-1 iMGLs. Upregulation of DAM-related and lysosomal-related genes such as *APOE, CCL3, CD68, CST7, GPNMB*, and *TREM2* was also blunted in KO-1 iMGLs after 24-hour myelin exposure (**Figure 7E, F, Supplement Table 4)**^9^.

At the pathway level, myelin exposure in WT-1 iMGLs at 2 hours led to enrichment among upregulated genes of pathways related to chemokine and cytokine activity, neuroinflammatory response, and proliferation (**Figure 7G**). In the KO-1 versus WT-1 comparison, many of these immune-related and proliferation pathways were instead downregulated, reflecting a blunted early transcriptional response of KO-1 iMGLs to myelin. By 24 hours, WT-1 samples showed enrichment of upregulated genes in DAM signatures, cholesterol metabolism, and maintenance of immune and inflammatory responses, alongside a concurrent decrease in proliferation. In the KO-1 versus WT-1 comparison, these responses were altered: DAM- and immune-related programs, as well as phagocytosis pathways, were downregulated, while proliferation pathways were upregulated in the KO-1, suggesting an attenuated or dampened transcriptional response relative to WT-1 controls (**Figure 7G**).

We observed that myelin treatment (2 hours and 24 hours) elicits similar gene-set-level transcriptional responses in WT-1 and KO-1 iMGLs (**Supplemental Figure 12D; see also Figure 7B, C, G and Supplemental Figure 12B**). Altogether, our data suggest that GPR34 is not required for microglial transcriptomic response to myelin stimulus but may modulate the magnitude of the response. We hypothesize that there are other receptors on microglia, besides GPR34, that can recognize myelin or the components of myelin.

## Discussion

Although GPR34 has long been recognized as a microglia-enriched GPCR whose expression is mis-regulated in various brain disease conditions, its normal physiological functions in microglia and its relevance to human neurodegenerative diseases have remained unclear^14,17,19,20,26,66,67^. In the mouse brain, loss of *Gpr34* shifted microglia toward DAM and inflammatory-related transcriptional states (**Figure 1**). The transcriptional modules upregulated in *Gpr34*-deficient microglia overlapped with those upregulated in the 5xFAD model of AD. *Gpr34* KO in 5xFAD mice further potentiated DAM-and inflammation-related features of microglia and exacerbated glial reactivity with no change in amyloid burden in this amyloidosis mouse model (**Figures 2-3**). Together these data suggest that GPR34 plays a role in preventing activation of microglial pathways that are associated with disease and inflammation. Our findings build on previous studies showing that *Gpr3*4 deficiency increases immune and cytokine gene expression^17,19^ and accelerates transition to DAM microglia^20^. Overall, these data support a model in which GPR34 helps maintain microglial homeostasis, consistent with its high expression in homeostatic microglia and suppression in disease-associated microglia, including in our reanalysis of a published DAM dataset^9^. The generation of the DAM and homeostatic gene sets is described in the Methods.

In 5xFAD brains, we found reduced *Gpr34* expression, consistent with reports that *Gpr34* levels are reduced in human brain tissue from neurodegenerative patient contexts^23^, although another recent paper reported the opposite trend^21^. Our RNA-seq and histopathological results from *Gpr34* KO;5xFAD mice appear to contrast with recent reports that *Gpr34* genetic disruption or inhibition with drugs alleviates neuroinflammation and promotes amyloid clearance in the APP/PS1 and 5xFAD models^16,20,21^ and with a separate report in which *Gpr34* agonist treatment promotes *in vivo* clearance of amyloid plaque in the APP/NLGF model^23^. These apparent discrepancies may reflect differences in model severity, experimental approach, disease stage, or timing of intervention. Notably, our study focuses on an earlier timepoint (6 months) in 5xFAD mice compared to 9 months of age in Zhou et al., 2025 when inflammatory pathways in microglia may be differently or more strongly engaged^21^. Lin et al., 2023 studied *Gpr34* knockdown in the hippocampus of APP/PS1 mice, introducing another layer of variability^16^. In addition, our study utilized a constitutive *Gpr34* KO, which can introduce developmental effects on microglia prior to the appearance of amyloidosis, in contrast to a pharmacological perturbation^21^ or genetic intervention after the onset of disease pathology^16,18^. Furthermore, the existing pharmacological compounds (GPR34 antagonists: YL-365, S-E49 and agonist compound 4B) are inadequate for in vivo studies due to their poor blood-brain barrier permeability^23,41,45^. This limitation restricts their utility for directly investigating GPR34 function in the brain and prevents straightforward comparison with our genetic intervention. Future systematic studies that compare the temporal and mechanistic dimensions of GPR34 function will be essential to clarify whether, when, and how its functions change during development or disease progression.

*GPR34*-deficient iMGLs displayed blunted Ca²⁺ and pERK signaling in response to LysoPS-derived agonists and myelin (**Figure 4**). Our findings align with prior reports showing that GPR34 mediates LysoPS- and myelin-derived signaling and regulates microglial lipid homeostasis^18,20,21^. Interestingly, while LysoPS-derived agonists of GPR34 such as Compound 4B activated robust signaling in iMGLs, LysoPS itself induced responses that were only partially dependent on GPR34. This suggests that there are additional receptors that can sense Lyso-PS and related lipids^14,26,28,45,68^. Interestingly, we found that *GPR34*-deficient iMGLs were only deficient in phagocytosis of myelin with no impairment in β-amyloid or E. coli uptake. This supports the idea that myelin uptake involves LysoPS/GPR34 interactions, since synthetic amyloid fibrils do not contain Lyso-PS and bacteria are very low in Lyso-PS. Our findings contrast with prior studies, which themselves reported variable outcomes^18,19,21,23^. Lin et al., 2024 found no effect of *Gpr34* loss on myelin uptake^18^. Etani et al., 2024 reported increased β-amyloid phagocytosis with GPR34 activation without affecting myelin^23^, and Preissler et al., 2015 described reduced uptake of zymosan and FBS-coated particles in *Gpr34*-KO mouse microglia^19^. These discrepancies likely reflect differences in model systems, assay conditions and other experimental factors. Two of these studies evaluated myelin uptake in primary mouse microglia^18,23^. However, primary mouse microglia rapidly downregulate GPCR expression upon removal from the CNS and during extended culture, unlike physiologically maintained iPSC-derived microglia, in which *GPR34* expression remains relatively stable^13^ (our unpublished data). Further, acute pharmacological activation to interrogate GPR34-dependent uptake^23^, may not correlate with the impacts of long-term genetic loss in a knockout model^19^.

Our in vivo and in vitro findings nonetheless suggest a potential link between GPR34’s role in regulating microglial states in vivo and myelin uptake in vitro cell culture. Recent studies indicate that myelin pathology, specifically myelin loss and increased myelin debris, is a feature of AD and may contribute to disease development and cognitive decline through mechanisms partly independent of other pathologies, such as amyloid accumulation^1,69–74^. In our transcriptomic dataset, we observed an overlap between the myelin-induced response of microglia in vitro and disease-associated microglia states in vivo, particularly in the 5xFAD model (**see public review comment#4 and Figures PublicReview#2 and 3**). Given this, we hypothesize that the abnormal myelin handling due to *GPR34* loss may, at least in part, activate disease-associated pathology and eventually affect cognitive function^71,72^. However, we did not test cognitive function in *Gpr34* mutants, which remains an important direction for future research. GPR34 signaling can modulate microglial state and appears to favor a homeostatic rather than inflammatory function of microglia. Therefore, we hypothesize that pharmacologic approaches that activate or inhibit GPR34 activity could be used to alter microglial state in neuroinflammatory and neurodegenerative disorders^23,41,45,75–77^. Whether stimulation or inhibition of GPR34 would be beneficial is likely to depend on the disease pathology, the stage of the disease, and the specific functions of microglia that are being targeted.

Our study is limited to a single disease model and relies primarily on genetic loss-of-function approaches in vivo. Although we used lysoPS and Compound 4B to prove GPR34 signaling pharmacologically in vitro, future studies should carefully test the impact of pharmacological modulation of GPR34 signaling in vivo through both agonist and antagonist approaches. Future research should also address sex-dependent variability and expand to additional disease contexts and examine how GPR34 interacts with other microglial receptors and lipid-sensing mechanisms^12,13,32,78–82^. These comparisons will help clarify whether GPR34 acts in parallel or synergistically with pathways mediating microglial homeostasis, lipid metabolism, and the inflammatory response.

## Acknowledgements

We thank Kris Dickson and Berengere Sauvagnat for their contributions to project strategy and project management, respectively. We are grateful to the members of the Sheng and Stevens labs for their valuable input, and to Jen Pan and Oivin Guicherit for their project discussions. We also thank Leiby Vargas, Matthew Beck, Nathan Chan, and Tyler Caron for assistance with mouse colony management; the Broad Institute Genomics Platform for support with RNA sequencing; the Broad Proteomics Platform for assistance with proteomics analyses; and the Flow Cytometry Core for access to instrumentation and technical support. We are especially grateful to Laurence Marie Daheron and Caroline E. Becker from the Harvard iPS Core Facility for generating the *GPR34* KO iPSCs. We thank Jeffrey Howard for administrative support, Elvira Dzhura and Abigail Desrochers for laboratory management, and Laney DiGiovanni for budgeting support.

## Author Contributions

M.S. conceived the project and supervised all aspects of the work. P.K. initiated and oversaw the iMGL project and supervised in vivo data analysis involving *Gpr34* KO and 5xFAD experiments. A.G. designed and conducted iMGL experiments: phagocytosis, RNA-seq, and myelin RNA-seq. M.K. conducted in vivo experiments and contributed to data analysis. Y.Z. co-initiated the iMGL platform, and Y.Z. and Q.X. performed pharmacological validation. W.E.M. generated the *GPR34* knockout clones, co-initiated and co-validated the iMGL platform, performed early iMGL RNA-seq experiment and contributed to in vivo experiments. S.S. analyzed RNA-seq data and contributed to experimental design under the supervision of J.L. J.D., C.D., and N.M. advised and contributed to iMGL experiments under the mentorship of B.S. S.N. performed in vivo immunohistochemistry and analysis under the supervision of H.B. E.Le. initiated in vivo experiments. X.A. conducted the *in vivo* RNA-seq experiments under the supervision of J.L. T.N. and M.D. contributed to iMGL preparation. K.J.S. and D.B. maintained and characterized the *Gpr34* knockout animals. J.Z. contributed to iMGL-associated RNA-seq experiments. A. V. and D.V. performed mass spectrometry proteomics analysis under the supervision of H.K. and S.A.C. E.Lo. and C.B. developed pharmacological compounds under the supervision of D.Mc., S.G., and M.W. D.Mi. contributed to RNA-seq data analysis. M.J. advised on all aspects of projects involving iMGL work. P.S.K., A.G., S.S., Y.L.Z., and M.K. wrote the manuscript with assistance from M.S., and input from all authors

## Disclosures, Funding statement

### Data and Code Availability

Code used for this manuscript is available on GitHub (see https://github.com/seanken/GPR34_DAM). Raw sequencing data have been uploaded to the Gene Expression Omnibus (GEO) under accession numbers **GSE327711** for the iMGL RNAseq data and **GSE327428** for the mouse RNAseq datasets. The processed data will be put in the Broad Single Cell Portal.

## GPR34 MS Methods Section + Resources Table

### A: Methods

#### Animals

*Gpr34* mutant mice were acquired from Taconic Biosciences (TF1866). *Gpr34* mutant mice were crossed with 5xFAD mice (The Jackson Laboratory, #034848) to produce wild-type, *Gpr34* KO, *Gpr34* KO; 5xFAD, and 5XFAD littermates used in scRNA-seq, snRNA-seq, and IHC experiments. To minimize variability, only 6-month-old wild-type, *Gpr34* KO, *Gpr34* KO; 5XFAD, and 5XFAD male mice were used in this study. All animals were housed at AAALAC-approved facilities on a 12-hour light/dark cycle, with food and water available *ad libitum*. All procedures involving these mice were approved by the Broad Institute IACUC (Institutional Animal Care and Use Committee) and conducted in accordance with the NIH Guide for the Care and Use of Laboratory Animals. Ear punched tissues from each mouse were sent out to the commercial vendor (Transnetyx, Cordova, TN) for genotyping using real time PCR with specific probes designed for each gene.

#### Euthanasia and Necropsy

6-month-old wild-type, *Gpr34* KO, *Gpr34* KO; 5XFAD, and 5XFAD male mice were anesthetized by administration of isoflurane in a gas chamber. While under anesthesia, transcardial perfusions were performed with ice-cold Phosphate-buffered saline (PBS). After perfusion, hippocampus from one hemibrain of each animal was quickly dissected out and placed in a pre-chilled 1.5mL Eppendorf containing ice-cold Hank’s Balanced Salt Solution (HBSS). Collected hippocampus samples were used for snRNA-seq and scRNA-seq (see below).

The other hemibrain from each animal were drop-fixed in 10X the brain volume of ice-cold, freshly made paraformaldehyde (PFA) for 48 hr at 4°C in a Falcon tube with mild agitation using a tube rocker. After drop fixation, hemibrains were moved to a Falcon tube containing ice-cold post-fixation PBS buffer solution (PBS with 0.01% sodium azide) and stored at 4°C until shipped to NeuroScience Associates, Inc. (NSA, Knoxville, TN) for processing. NSA used MultiBrain® technology to embed 40 hemibrains together in a single gelatin matrix and freeze sectioned at 30 μm in the sagittal plane.

#### Immunohistochemistry and Whole Slide Imaging

Immunohistochemistry was performed by NSA using free-floating sections. Primary antibodies against GFAP (chicken; EnCor Biotechnology, CPCA-GFAP) and Iba1 (goat; Novus Biologicals, NB100-1028), together with the corresponding secondary antibodies anti-chicken-Cy3 (Jackson ImmunoResearch, 703-165-155) and anti-goat-Cy3 (Jackson ImmunoResearch, 706-165-147), were used at a dilution of 1:500. Immunofluorescent slides were imaged by NSA at 40x magnification. Three ROIs were acquired from anterior, middle, and posterior regions of the cortex per section per brain. They were selected consistently across animals. All whole slide image analysis was performed in a blinded manner using FIJI (version 1.54f). All images were evaluated in 8-bit grayscale. GFAP was analyzed using a local threshold and measured for percent-area. All thresholds were kept consistent between all slices.

#### Immunohistochemistry and *In vivo* microglia engulfment of plaque

Free-floating sections were blocked in blocking solution (10% normal donkey serum in PBS with 0.5 % Triton X-100 and 0.5% Bovine-serum albumin (BSA)) and incubated overnight at 4°C with the primary antibody in a carrier solution (1% normal donkey serum in PBS with 0.5 % Triton X-100 and 0.5% Bovine-serum albumin (BSA)). Sections were washed with 1XPBS (3 x 5 min each) before being placed in secondary antibodies in carrier solution for 2-3 hours, covered, and at room temperature. Sections were washed in 1XPBS (3 x 5 min each) and coverslipped with DAPI Fluoromount (SouthernBiotech 0100-20).

Primary antibodies: Iba1 (1:1000, chicken, Synaptic Systems, 234009), β-amyloid (6E10, 1:1000, mouse, BioLegend, 803001), and CD68 (1:1000, rat, Bio-Rad, MCA1957). Secondary antibodies were used at 1:750.

Images were acquired on a Zeiss LSM 900 confocal microscope using 63× magnification. Frame size of 0.043 μm x 0.043 μm was used for all images. 40-50 Z-stacks were acquired with an interval size of 0.3 μm. 4-5 ROI were acquired in the cortex per section per brain. Those ROIs were consistently selected from anterior, middle, and posterior regions of the cortex across animals rather than based on local plaque density. Imaris 10.2 3D surface rendering was used to first create a surface on the Iba1 channel. Subsequently, the CD68 channel was masked to the Iba1 surface to obtain only CD68 within the analyzed cell. A surface was created on the CD68 masked channel, and a number of voxels > 10 filter was applied. 6E10 channel was then masked to the CD68 surface to obtain 6E10 inside CD68+ vesicles and a surface was created on this masked 6E10 channel. Total 6E10 was measured using surface function. Imaris 3D surface function algorithm settings were the following for all channels: “Absolute intensity” and grain size of 0.1 μm were used. The threshold values were adjusted to optimize visualization of individual channels and varied between channels but were kept constant within a channel for all images analyzed.

#### Microglia isolation and single-cell RNA-seq

These experiments started with the mouse brain hippocampus samples harvested in 1x HBSS buffer (Thermo Fisher Scientific, 14185052). The tissue was transferred to a petri dish and finely diced on ice. A small portion of the diced sample was reserved in RNAlater (Thermo Fisher Scientific, AM7020) for snRNA-seq^1^. The remaining sample was rinsed with 2 ml ice cold HBSS and transferred to a douncing device vial (Sigma, D8938). The tissue was dounced with pestle A 15 times and followed by 15 times with pestle B. The 2 ml homogenized tissue was then passed through a pre-wetted 70-micron filter (VWR, 21008-952) and collected into a 15 ml tube. The douncing vial was also rinsed with another 2 ml ice cold HBSS and passed through the same filter. The sample was centrifuged at 300 g at 4°C for 5 minutes. After supernatant removal, the cell pellet was resuspended in 1550 µl of ice-cold D-PBS (Thermo Fisher Scientific, 14287080). 450 µl prechilled Debris Removal Buffer (Miltenyi Biotech, 130-109-398) was then added and mixed by pipetting. Two ml of cold D-PBS were gently overlayed on top of the sample mix and centrifuged at 3000 g and 4°C for 10 minutes. The upper layer and second cloudy debris layer were removed, and the volume was brought back to 7.5 ml with cold D-PBS and the suspension was mixed by gently inverting three times. The solution was centrifuged at 1000 g and 4°C for 10 minutes and the supernatant was removed. The cell pellet was then resuspended in 90 µl ice cold PB buffer (0.5% BSA, 0.2 U/µl Recombinant RNase Inhibitor (Takara, 2313A), 1x PBS) and 10 µl CD11B microbeads (Miltenyi Biotech, 130-126-725) were added. The solution was mixed by gently pipetting and then incubated in the dark for 15 minutes. Another 500 µl ice cold PB buffer was added at the end of the incubation. During the incubation time, the MS column (Miltenyi Biotech, 130-042-201) was prepared by placing it on the magnet (Miltenyi Biotech, 130-042-108) and prewetting with 500 µl ice cold PB. The sample and microbead mix were then pipetted into the column collection chamber and let drip through via gravity. After all the sample solution passed through, the column was washed with 1.5 ml cold PB. The column was then removed from the magnet and the CD11B labeled microglia were flushed out with 1 ml of ice cold PB. The cell suspension was spun down at 300 g and 4°C for 10 minutes and the supernatant was removed. The pellet was resuspended in 50 µl PB. Only 5 µl in a 1/4x dilution was used and stained with AOPI (Nexcelom Bioscience, CS2-0106-5ML) to count and assess viability using the Cellometer K2 cell counter (Nexcelom Bioscience).

Depending on the yield, most of the isolated microglia were loaded onto a Chromium Next GEM Chip G (10x Genomics, PN-1000120) aiming to recover up to 10,000 cells. The single-cell RNA-seq libraries were generated using the Chromium™ Single Cell 3’ Library & Gel Bead Kit v3.1 (10x Genomics, PN-1000121). Twenty-four libraries from different animals were pooled based on molar concentrations and sequenced on one NextSeq 500 flow cell and two NovaSeq 6000 SP flow cells (Illumina) with 28 bases for read 1, 55 bases for read 2 and 8 bases for Index read 1.

#### Nuclei isolation and single-nucleus RNA-seq

Nuclei were isolated from RNAlater preserved hippocampus samples harvested earlier. The tissue was placed into a douncing vial containing 2 ml cold EZ-prep lysis buffer (Sigma, NUC101-1KT) and dounced with pestle A and followed by pestle B with 15 times each. A 30 µm filter (Sysmex, 04-004-2326) was pre-wet with 0.5 ml MST buffer (containing 10 mM Tris, 146 mM NaCl, 1 mM CaCl_2_, 21 mM MgCl_2_, 1% BSA (Miltenyi Biotech, 130-091-376), 0.02% Tween-20 (Roche, 11332465001), 40 U/ml of Recombinant RNase Inhibitor (Takara, 2313A)) with the lysate being passed through the filter and collected into a 15 ml tube. The douncing vial was rinsed with 1.5 ml of the MST buffer and passed through the same filter. The lysate was spun down in a swinging-bucket centrifuge at 500 g and 4°C for 5 minutes. After supernatant removal, the nuclei were washed by resuspending in 1 ml MST buffer and transferring to a 1.5 ml lo-bind tube (Eppendorf, 22431021). The solution was spun down in a benchtop fixed-angle centrifuge at 500 g and 4°C for 5 minutes and the supernatant was removed. This wash step was repeated one more time before the final resuspension in 100 µl RB buffer (containing 1x PBS, 1% BSA and 0.2 U/µl RNaseIn) and passing through a 20 µm filter (Sysmex, 04-004-2325). The AO (Nexcelom Bioscience, CS1-0108-5ML) stained nuclei were counted using a Cellometer K2 cell counter with a gate setting of 5-20 µm.

The snRNA-seq libraries were prepared with the same process as for the scRNA-seq libraries (described above), to recover up to 10,000 nuclei. Twenty-four libraries were pooled and sequenced on one NextSeq 500 flow cell and four NovaSeq 6000 SP flow cells (Illumina) with 28 bases for read 1, 55 bases for read 2 and 8 bases for Index read 1.

#### Processing of single-cell data

FASTQ files were processed with Cell Ranger v6.1.2, with the refdata-gex-mm10-2020-A Cell Ranger reference (10x Genomics) and with the expected number of cells set based on estimated loading^2^. The downsampleReads command in DropletUtils v1.14.2 was used to downsample all samples to the same number of reads per cell, then the emptyDropsCellRanger command was used to remove empty droplets^3^. This count matrix was then loaded in R v4.0.3. A Seurat v4.0.0 object was created, cells with less than 200 genes were removed, and the data was LogNormalized with NormalizeData^4^. Variable genes were extracted with FindVariableFeatures, data was scaled with ScaleData, and PCA was run with RunPCA with npcs=60. Dimensionality reduction was performed on the count data with scGBM v0.1.0, using the gbm.sc command with M=25 and subset=50,000, using just the variable genes^5^. Clusters were calculated with the FindNeighbors and FindClusters commands using dims=1:25 and the scGBM reduction. The UMAP was generated with the RunUMAP command with the same arguments used for clustering. We calculated the percent of UMIs coming from the mitochondria, ribosomal proteins, and a doublet score using scds v1.6.0, run separately on each 10x channel^6^. We used Azimuth v0.3.2 and an existing reference from the Allen Brain Institute^4^ to annotate the data and identify which clusters were microglia versus not. We removed two outlier samples (s1 and s16), both of which had >50% non-microglia cells and removed non microglia cells from the other samples. Variable genes were calculated again, and *Gpr34* was removed from the variable gene list to avoid bias. scGBM was run again, this time with M=10 and subset=30,000, followed by clustering and UMAP using the same commands as previously, except with dims=1:10. Clustering was run at both the resolution=0.8 and 1.5 level, with the 1.5 level being used for downstream annotation. Clusters were annotated with known microglial cell types with known marker genes^7^, and possible doublet clusters (identified based on expression of known markers for non-microglia cell types) were removed.

To test for cell type composition changes between conditions, we used the propeller.ttest test in speckle v0.0.2^8^ with robust=TRUE, trend=FALSE, sort=TRUE, using the asin transform and adding batch as an additional command. For differential expression analysis, sample level pseudobulks were made with the AggregateExpression command in Seurat with slot=”counts”. We then used edgeR v3.32.1, first to remove lowly expressed genes with the filterByExpr argument, then to test for DE with the LRT test (glmFit command followed by glmLRT, both in the edgeR package)^9^. Note batch was included as an extra covariate in DE testing. For enrichment analysis, fgsea v1.16.0 was used in all cases using the score calculated as a sign of the logFC time negative log pvalue^10^. KEGG gene sets were downloaded with the download_KEGG tool from clusterProfiler v3.18.1, GO gene sets were extracted with the AnnotationDbi v1.52.0 tool ((Pagès H, Carlson M, Falcon S, Li N (2025). *AnnotationDbi: Manipulation of SQLite-based annotations in Bioconductor*. doi:10.18129/B9.bioc.AnnotationDbi)), while other gene sets (marker genes for DAMs and homeostatic cells, up and down regulated genes in AD were extracted from the literature^11^.

#### Processing of snRNA-seq data

The processing of snRNA-seq data was similar to that of scRNA-seq data, except Cell Ranger was run with --include-introns and emptyDropsCellRanger was run with umi.min=200. Processing and downsampling were performed as described for the scRNA-seq data, except scGBM was run with M=25 and subset=30,000, and nuclei with <150 genes were removed. QC metrics were calculated and added to the Seurat object. Cells were annotated with Azimuth and doublets scores were calculated as for the single-cell data. We removed one large cluster (cluster 0) that didn’t seem to correspond to any cell type and had high doublet scores, then recalculated with UMAP and clusters with resolution=1.5. We annotated the clusters by cell type using the Azimuth labels, known markers (such as *Slc17a6*/Slc17a7 for Excitatory, *Gad1*/*Gad2* for inhibitory, *Plp1*/*Mobp*/*Mbp* for ODCs, *Pdgfra* for OPC, *Csf1r* for microglia, *Flt1* for Endothelial, *Slc1a3*/*Gfap* for astrocytes), and doublet scores. We removed contaminating cell types from the cortex and doublets. We jointly subclustered the oligodendrocyte and oligodendrocyte precursor (OPC) clusters (running gbm with M=15, followed by clustering with resolution=1.2) to identify a doublet subcluster which was then removed from the overall Seurat object. We then subclustered the oligodendrocytes, microglia, and astrocytes with scGBM followed by RunUMAP and FindNeighbors/FindClusters as described previously, annotating the clusters using known markers (*Gfap* for activated vs not activated astrocytes, *Plp1* for ODC, *Pdgfra* for OPC) and presence/absence in 5xFAD mice vs wild types (cell type present in 5xFAD but not in WT are labelled as disease associated, consistent with the literature^11^.

Cell type composition analysis, differential expression analysis, and enrichment analysis were performed as described for the scRNA-seq data. For differential expression we excluded nuclei with >1% mitochondrial UMIs or >10,000 UMIs.

#### Processing of bulk RNA-seq data

To extract RNA-seq quantification we used an in-house pipeline as reported previously^12^. Reads were mapped to the GrCH38 genome with STAR v2.7.9a^13^ and QC was calculated on the resulting BAM file with PICARDTool CollectRNASeqMetrics v2.27(https://broadinstitute.github.io/picard/). Quantification was performed with Salmon (v1.6.0)^14^ with the arguments -l A --posBias --seqBias --gcBias --validate Mapping, with the full genome as decoys. These data were loaded into R with tximport v1.18.0^15^. The count data was loaded into a DESeqDataSet object using DESeq2 v1.30.1^16^. For each analysis, the DESeq object was downsampled to samples of interest. PCA was performed by using the vst and plotPCA functions in DESeq2. Differential expression was performed with the DESeq function in DESeq2. More specifically, genes with <10 reads total were excluded, and the DESeq command was used, followed by the lfcShrink command with type= “normal”. We also extracted the results without shrinkage using the results command, which was fed into TRADE v0.1.0 to extract the TWI score^17^. Enrichment analysis was performed as described for the scRNA-seq data, except the “stat” statistic from DESeq2 was used and the KEGG gene sets were extracted using the limma v3.46.0 commands getGeneKEGGLinks and getKEGGPathwayNames^18^. To identify the iMGL marker genes, we downloaded the iMGL scRNA-seq data from the following publication^11^. We used the wilcoxauc command in presto v1.0.0 to extract DE genes^19^. We then used these DE genes to create marker lists for each iMGL cell type. For KO vs WT differential expression analysis with no treatment, we performed DE separately on each batch (two batches) without shrinkage, then combined the results from the two batches with a meta-analysis random effect approach. This was performed with the rma function in the metafor v4.8 package^20^, with the argument method=”REML”, using the log fold changes and standard errors reported by DESeq2 as inputs.

#### Generation of *GPR34* KO iPSC line

*GPR34* KO clones were generated from WTC11^21^ cells by the Harvard University Stem Cell Institute by introducing a CRISPR construct and sgRNAs targeting two nucleotides for deletion (X chromosome, bases 41695777-41695778 [KO-1] and bases 41695816-41695817 [KO-2]) into WTC11 iPSCs to introduce early stop codons. Guide RNA for KO-1 was GATAGCAATTTTTCATCCAT and guide RNA for KO-2 was CACGATGAAAATAACAGAGT. As antibodies proved ineffective at detecting GPR34 protein in iPSCs or iMGLs, KO was confirmed by DNA sequencing. Cell lines were sequenced through Genewiz (Azenta) using primers AGACAATGAGAAGTCATACC and CAGTGTTCCCACAACCTTGC. DNA was extracted from iPSCs via DNeasy kit (Qiagen #69506) and sequences were aligned using SnapGene (GSL Biotech). Protein was shown to be non-functional in reporter assays.

#### iPSC Culture

iPSC-derived microglia (iMGL) were differentiated from a WTC11 line^21^. iPSCs were maintained on Matrigel-coated 6 well plates (Corning) according to the manufacturer’s specifications in Essential 8 (E8, Thermo Fisher Scientific) culture medium. Culture medium was replenished every day with fresh medium. Cells were passaged every 4-6 days using ReLeSR (STEMCELL Technologies) according to manufacturer’s specifications. Unless otherwise stated, for all in vitro experiments, iPSCs were cultured in 5% O2, 5% CO2 at 37 C.

#### iMGL differentiation

iPSC lines were maintained in E8 media until seeding for differentiation. At ∼80% confluence, hematopoietic differentiation was started as previously described^22^. Cells were dissociated using Accutase (Stem Cell Technologies), centrifuged for 5 minutes at 300g and counted using trypan blue (Thermo Fisher Scientific). The pellet resuspended at 100,000 cells per mL in E8 media containing 10 µM Y-27632 ROCK Inhibitor (Selleckchem) and were plated at 200,000 cells per well in a low-adherence six-well plate (Corning). For the first 10 days, HPC media was used (50% IMDM (Thermo Fisher Scientific), 50% F-12 (Thermo Fisher Scientific), ITSG-X 2% v/v (Thermo Fisher Scientific), Glutamax (1x, Thermo Fisher Scientific), chemically defined lipid concentrate (1x, Thermo Fisher Scientific), nonessential amino acids (Thermo Fisher Scientific), l-ascorbic acid 2-phosphate (64 µg/ml, Sigma), poly(vinyl) alcohol (10 µg/ml, Sigma), and monothioglycerol (400 µM, Sigma). On day 0, embryoid bodies were gently collected and centrifuged at 100 g for 5 minutes. They were then resuspended in HPC media supplemented with 1 µM ROCK inhibitor, FGF2 (50 ng/ml, Thermo Fisher Scientific), BMP4 (50 ng/ml, Thermo Fisher Scientific), Activin-A (12.5 ng/ml, Thermo Fisher Scientific) and LiCl (2mM, Sigma), and incubated in a hypoxic incubator (5% O2, 5% CO2, 37 °C). On day 2, cells were gently collected, and the media was changed to HPC media supplemented with FGF2 (50 ng/ml, Thermo Fisher Scientific) and VEGF (50 ng/ml, PeproTech), before being returned to the hypoxic incubator. On day 4, cells were gently collected and the media was changed to HPC media supplemented with FGF2 (50 ng/ml, Thermo Fisher Scientific), VEGF (50 ng/ml, PeproTech), TPO (50 ng/ml, PeproTech), SCF (10 ng/ml, Thermo Fisher Scientific), IL6 (50 ng/ml, PeproTech) and IL3 (10 ng/ml, PeproTech); then, cells were incubated in a normoxic incubator (20% O2, 5% CO2, 37°C). On days 6 and 8, 1 ml of day 4 media was added to each well. On day 10, cells were collected, counted using trypan blue and frozen in Cryostor (Sigma Aldrich) in aliquots of 600,000-1,200,000 cells.

To start iMGL differentiation, cells were thawed, washed 1× with PBS and plated at 100,000 cells per well in a six-well plate coated with Matrigel in iMGL media ((DMEM/F12 (Thermo Fisher Scientific), ITSG (2% v/v, Thermo Fisher Scientific), B27 (2% v/v, Thermo Fisher Scientific), N2 (0.5% v/v, Thermo Fisher Scientific), monothioglycerol (200 µM, Sigma), Glutamax (1×, Thermo Fisher Scientific), nonessential amino acids (1×, Thermo Fisher Scientific)), human insulin (5 µg/ml, Merck Millipore/Sigma-Aldrich) supplemented with M-CSF (25 ng/ml, PeproTech), IL-34 (100 ng/ml, PeproTech) and TGFB-1 (50 ng/ml, PeproTech). Cells were fed every 2 days and replated at day 22. On day 30, cells were collected and replated in iMGL media supplemented with M-CSF (25 ng/ml, PeproTech), IL-34 (100 ng/ml, PeproTech), TGFB-1 (50 ng/ml, PeproTech), CD200 (100 ng/ml, VWR) and CX3CL1 (100 ng/ml, PeproTech). Cells were used at day 40 for functional, transcriptomic, and proteomic assays. iMGL differentiation was assessed at day 40 (expression of CD45, CD11b, P2RY12, CX3CR1 and LIVE/DEAD Fixable viability dye) by flow cytometry (see below for protocol). During the quality control on day 40, we observed similar protein expression among the replicates. At no point were iMGLs exposed to serum of any kind. Throughout the differentiation, cell viability was >90% as measured by trypan blue.

#### Validation of iMGL differentiation by flow cytometry

For iMGL differentiation quality control, iMGLs were detached using cold DPBS then resuspended in fluorescence-activated cell sorting (FACS) buffer (DPBS containing 1% bovine serum albumin and 0.5 mM EDTA). Samples were incubated for 15 minutes in human Fc block (BD Biosciences), followed by 1 hour staining with conjugated antibodies (see Resources Table) at 4 °C. Samples were washed three times with FACS buffer and resuspended in 200 µL of FACS buffer for flow cytometry. Samples were run on a CytoFLEX S analyzer (Beckman Coulter) until at least 2,000 cells were recorded. For analysis, cells were identified according to the following gating: (1) cells versus debris (FSC-A versus SSC-A), (2) singlets (FSC-A versus FSC-H), and antibody-specific gating based on a negative control sample. Antibodies for staining iMGLs were CD45-AF488 (Biolegend), CD11b-AF488 (BioLegend), P2RY12-PE (BioLegend), Cx3CR1-AF647 (BioLegend), LIVE/DEAD Fixable Near-IR 876nm (Thermo Fisher Scientific), 1:100 for all.

#### Mass spectrometry on iMGL lines

iMGLs on day 40 were detached using ice-cold DPBS, spun down twice for 5 minutes at 1,000 g, and then cell pellets were flash frozen over dry ice. A total of 4 technical replicates were analyzed per genotype, each from pools from 6-well plates seeded at 100,000 cells/well.

#### Lysis and Digestion

Microglia cells underwent denaturing lysis in SDS to prepare for S-Trap digestion. Samples were lysed on ice in ∼100 µL SDS lysis buffer (5% SDS, 50 mM TEAB pH 7.55, 2 mM MgCl_2_, 2 µg/ml Aprotinin, 10 µg/mL Leupeptin, 1 mM PSMF, 10 mM NaF) for 15 min. Samples were then treated with 1 µL 250 units/μL Benzonase (Thomas Scientific, E1014-25KU) to shear DNA, mixed again, and incubated on top of ice for another 15 min. The lysates were cleared by centrifugation for 10 min at 20,000 g at 4°C and the supernatant was prepared for S-Trap digestion. Protein concentration was estimated using a BCA protein assay. Disulfide bonds were reduced with 5 mM DTT for 1 hour at 25°C and 1000 rpm shaking, and cysteine residues alkylated in 10 mM IAA in the dark for 45 min at 25°C and 1000 rpm shaking. 12% phosphoric acid was added at a 1:10 ratio of lysate volume to acidify, and proteins were precipitated with 6× sample volume of ice-cold S-Trap buffer (90% methanol, 100 mM TEAB) in either S-Trap micro of S-Trap mini cartridges depending on the protein yield. The precipitated protein was mixed by pipetting then centrifugated at 4000 g for 1 minute to remove the buffer. The precipitated proteins were washed 3× 4000g for 1 min. To digest the deposited protein material, digestion buffer (50 mM TEAB) containing both trypsin and LysC, each at 1:50 enzyme: substrate, was passed through each S-Trap column with 1 min centrifugation at 3000g. The digestion buffer was then added back atop the S-Trap, and the cartridges were left capped overnight at 25 °C. Peptide digests were eluted from the S-Trap sequentially at 4000 g for 1min, first with 50 mM TEAB, next with 0.1% FA, and a final elution with 50% ACN/0.1% FA. Eluted samples were frozen at -80°C and vacuum-centrifuged until dry.

Dried down samples were reconstituted in 500 µL of 1% FA and desalted in 40mg Sep-Pak tC18 96-well Plate (Waters, SKU:186002320) using a positive pressure manifold. 80ug of digested sample was desalted from each sample. Briefly, C18 wells were conditioned sequentially with 1 mL of 100% ACN, 1mL of 50% ACN/0.1% FA, and 4x 1mL of 0.1% TFA. Next, acidified peptides were loaded onto the C18 wells and washed with 3x 1mL per uL of 0.1% TFA, followed by 1x 1mL 1% FA. Desalted peptides were then eluted from the C18 resin using 2x 500 µL of 50% ACN/ 0.1% FA, frozen at -80°C, and vacuum-centrifuged until completely dry. Dry desalted peptides were then reconstituted in 1% FA and quantified using a BCA assay. For mass spectrometry analysis 600 ng of peptides from each sample were loaded onto Evotips (Evosep) adhering to the manufacturer’s guidelines for LC/MS analysis.

#### Liquid Chromatography - Mass Spectrometry (LC/MS) Analysis

Samples were analyzed on timsTOF HT mass spectrometer (MS) coupled to Evosep One (Evosep) LC system with mobile phase A of 0.1% formic acid/water and mobile phase B of 0.1% formic acid/acetonitrile. 600 ng of each sample was injected onto 75 mm ID x 150 mm length Aurora Elite^TM^ CSI C18 (IonOptics) column and eluted with Whisper Zoom 20SPD (Evosep) LC method. DiaPassef analysis is performed with ion mobility range of 0.70 1/K0 [V-s/cm-2] to 1.45 1/K0 [V-s/cm-2], ramp and accumulation time of 70 ms. DIA windows range from 330 m/z to 1523 m/z with 36 isolation windows and a cycle time of 1.67 sec. The collision energy was ramped as a function of increasing mobility starting from 30 eV at 0.8 1/K0 [V-s/cm-2] to 55 eV at 1.45 1/K0 [V-s/cm-2].

#### Data analysis

Raw files were searched against the reviewed human proteome database obtained from UniProt, without isoforms (20,462 entries), with contaminants appended to the FASTA file. Data were searched using Spectronaut 19 in direct DIA mode using quantification settings with a precursor Q-value cutoff of 0.01, a precursor PEP cutoff of 0.2, protein Q-value cutoff (experiment) of 0.01, a protein Q-value cutoff (run) of 0.05, and a protein PEP cutoff of 0.75. Carbamidomethylation was set as fixed modification, and N-terminal acetylation and methionine oxidation were set as variable modifications. A spectronaut protein summary report was generated following the search and the PG.Quantity column was used for protein group quantification in downstream analyses. Non-human contaminants were filtered from the report. Statistical analysis was performed using the Proteomics Toolset for Integrative Data Analysis (Protigy, v1.0.7, Broad Institute, https://github.com/broadinstitute/protigy). The protein summary report was filtered with 70% missing values cut off, then log_2_ transformed and median (non-zero) normalized. A 2-sample t-test was performed comparing wild type vs *GPR34* KO samples. Adjusted p-values were calculated using the Benjamini–Hochberg FDR approach and identified proteins with an adjusted p-value<0.05 were considered statistically significant.

#### Gene set enrichment analysis (GSEA)

GSEA^23^ was performed using the R Bioconductor package fgsea^19^, with human C5 ontology sets (v7.5) obtained from the Molecular Signatures Database (MSigDB)^24^. Proteins were pre-ranked by their moderated t-statistics comparing *GPR34* KO versus WT. For proteins with multiple isoforms, the isoform with the largest effect size was selected for pre-ranking. The gene symbols of the proteins were used for fgsea. Because enrichment analyses were conducted using gene symbols, we use the terms “gene set” and “protein set” interchangeably throughout this study. Gene sets with FDR-adjusted p-values < 0.05 were considered significant.

#### Phagocytosis

Substrates were generated or acquired as follows: myelin was isolated as previously described^25^, while human synthetic amyloid beta 1-42 pre-formed fibrils (StressMarq Biosciences) and E. Coli Bioparticles (Thermo Fisher Scientific) were ordered for use. Myelin and amyloid fibrils were conjugated to pHrodo (Deep Red TFP Ester; Thermo Fisher Scientific), according to manufacturer’s protocol. Briefly, substrates were incubated with 1 mL pHrodo per 1 mg substrate for 2 hours at room temperature, protected from light, washed three times using HBSS and flash frozen in dry ice to be placed in -80°C until use. *E. Coli* Bioparticles were acquired preconjugated and ready for use upon HBSS reconstitution.

Phagocytosis of substrates was assessed over time using the Opera Phenix Live Cell Imaging System (Revvity), followed by flow cytometry. Briefly, iMGLs were seeded at 15,000 cells per well in 96-well PhenoPlates (Revvity), utilizing only the inner 60 wells to minimize edge effects. After a 30-minute pretreatment with the desired compound or 0.1% DMSO control, the substrate of interest was added. Plates were returned to the incubator and imaged at 5- and 24-hours post-treatment using Opera Phenix. At each time point, brightfield, digital phase contrast, and the AF647 fluorescent channel were captured. Image analysis was performed with Harmony software (Revvity), which applied cell delineation algorithms for accurate object recognition and segmentation. The mean pHrodo intensity within delineated iMGLs was quantified and averaged to assess phagocytic activity.

Following 24-hour imaging, cells were detached using ice-cold PBS containing DAPI to assess viability and transferred to a 96-well U-bottom flow cytometry plate (VWR International). Samples were acquired on a CytoFLEX flow cytometer (Beckman Coulter) until a minimum of 2,000 events were recorded per condition. Flow cytometry analysis involved sequential gating for (1) cell versus debris (FSC-A vs SSC-A), (2) singlets (FSC-A vs FSC-H), and (3) viability, based on exclusion of DAPI-positive cells. Phagocytosis was quantified as the median fluorescence intensity of pHrodo for each sample.

#### Myelin preparation and iMGL treatment for RNA sequencing

Myelin was isolated as previously described^25^. Briefly, C57BL/6J mice were transcardially perfused with cold HBSS, and whole brains were extracted to undergo sucrose-density centrifugation and osmotic shock to isolate a light-density myelin membrane fraction. Protein concentration was determined via bicinchoninic acid (BCA), and 1 mg/mL stocks were generated for experimental use. iMGLs were treated with myelin for 2 hours and 24 hours respectively before transcriptomic analysis.

#### Bulk RNA-seq library preparation and sequencing

On day 40, iMGLs were collected for RNA sequencing with 6 technical replicates per genotype. Each sample was harvested from independent wells seeded at 50,000 cells per well, originating from the same differentiation batch. Alternatively, for a second experiment, on day 40 iMGLs were treated with myelin (6 µg/ml) for 2 or 24 hours, or PBS. A total of 36 samples were collected, representing two genotypes, 3 treatments, and six replicates per condition. Again, each sample was harvested from independent wells seeded at 50,000 cells per well, originating from the same differentiation batch.

RNA extractions were performed using RNeasy Plus Mini Kit (Qiagen) according to manufacturer’s protocol. Briefly, samples were collected using RLT buffer (Qiagen) and homogenized through use of Qiashredders. Samples were then placed into columns and bound to RNeasy silica membranes. Contaminants were washed away, and columns were treated with DNase (Qiagen provided) to digest any residual DNA, and concentrated RNA was eluted in water. The RNA concentration was measured using a NanoDrop Spectrophotometer and RNA integrity (RIN) was measured with RNA pico chips (Agilent) using a 2100 Bioanalyzer instrument (Agilent). Purified RNA was stored at -80 C until library preparation for bulk RNA-sequencing analysis. Bulk RNA sequencing libraries were prepared using Illumina TruSeq Stranded mRNA Kit (Catalog #20020595) per manufacturer’s instructions. 200 ng of total RNA from each sample was used and the concentration of resulting cDNA library was measured with High Sensitivity DNA chips (Agilent) using a 2100 Bioanalyzer Instrument (Agilent). A pooled 5 nM normalized library was prepared, and sequencing was performed on a NovaSeq X (Illumina) with 50 bases each for reads 1 and 2 and 8 bases each for index reads 1 and 2.

#### Quantification and statistical analyses

Unless otherwise indicated, statistical analyses were performed using GraphPad Prism Software (Version 10.5.0 (673)) or R. Statistical details including specific tests used, exact value of n, e.g. number of animals, cells, or independent trials, and precision measures e.g., mean, median, SD, SEM, are indicated in the corresponding figure legend. All *in vitro* data are from at least two independent experiments. All statistical analyses performed were two-tailed.

#### Calcium Assay

One day prior to conducting the assay, iMGLs from div40 were seeded at 15K/well density in 50 µl fresh medium in 384-well clear-bottom plates coated with poly-D-lysine(Corning, BioCoat, product #356697). The cell plates were incubated in a 5% CO2 humidified incubator at 37°C overnight. On the day of the assay, media was aspirated using a plate washer, and 25 µl Fluo-4 dye at 2 µM in buffer was added to each well. The assay plates were incubated for 2 hours at room temperature in the dark. After two washes with 25 µl buffer, the plates containing 20 µl buffer in each well were placed in the FLIPR Penta™ instrument (Molecular Devices). The assay was performed at 37°C. Changes of fluorescence were measured over time with excitation 470–495 nm and emission 515–575 nm at a 1 Hz sampling rate. Ligands were added using the Flipr inline liquid dispenser. The assay buffer is HBSS with 20 mM HEPES and 0.1% BSA (fat acid free sigma A8806), pH=7.3. The peak response over baseline from FLIPR after each ligand addition was used for data analysis. The medium was the same as used in maintaining iMGLs after DIV30.

#### ERK Phosphorylation Assay

iMGLs from div40 were seeded at 15K/well density in 100 µl fresh medium in a 96-well plate (Costar 8795BC). The cell plates were incubated in a 5% CO2 humidified incubator at 37°C overnight. On the day of the assay, 50 µL of medium were aspirated from each well. 25 µl of medium or antagonist were added to each well and preincubated for 10 minutes at 37°C before 25 µl agonist addition. Cells were then lysed after 5 minutes at room temperature and assayed for phosphorylated ERK1/2 (pT202/Tyr204) and total ERK1/2 using an ELISA kit (Thermo Fisher Scientific 85-86013). Absorbance was read on a SpectraMax iD5 at 450 nm. The medium was the same as used in maintaining iMGLs after DIV30.

#### cAMP Assay

The iMGL control or *GPR34* KO cells at div40 dissociated from 6-well plates were replated at a density of 10,000 cells/well in 30 µl of fresh medium in 384-well plates coated with poly-D-lysine. The cell plates were incubated overnight at 37°C in a 5% CO2 humidified incubator. On the day of the assay, cells were washed three times with the buffer (1xHBSS, 5 mM HEPES, 0.5 mM IBMX, 0.1% BSA, pH 7.4) before adding 10 µl of ligand or buffer control. After a 10-minute incubation at room temperature, 5 µl of Forskolin at different doses was added to each well, and the plate was incubated with Forskolin for an additional 20 minutes at room temperature. The cAMP level was detected by a Time-resolved Fluorescence Resonance Energy Transfer (TR-FRET) competition assay using the Lance Ultra cAMP kit from Perkin Elmer (TRF0263). The FRET signal was measured using a PHERAstar FSX multi-mode microplate reader (BMG Labtech) with a TR-FRET protocol, employing excitation at 337 nm, emission filter at 665 nm, and a second emission filter at 620 nm.

#### Immunocytochemistry and Analysis

The iMGLs of *GPR34* WT control or *GPR34* KO at div42 were replated at a density of 10,000 cells/well in 384-well PhenoPlates (Revvity Product #6057302). After two days of recovery in 37°C, 5% CO_2_, the iMGLs were fixed in 4% paraformaldehyde (PFA) in media for 15 minutes, followed by permeabilization and blocking (5% normal goat serum and 0.2% Triton-100 in PBS) for 15 minutes. iMGLs were then incubated with primary antibodies overnight at 4°C, followed by three washes with 0.025% Triton-100/PBS. Cells were then incubated with a secondary antibody for 40 minutes at room temperature. This was followed by four washes with 0.025% Triton-100/PBS before imaging. The primary antibodies and their concentrations were: IBA1 (Fujifilm Wako, 019-19741) 1/2000; P2RY12 (Sigma, HPA014518) 1/200; C3aR (Santa Cruz SC-133172) 1/200; CD45 (BioLegend, 304017) 1/100; C1q (Labome, A10136) 1/1000. The secondary antibodies and their concentrations were: Goat anti-mouse 488 (Thermo Fisher Scientific, A11029) 1/2000 and Goat anti-rabbit 555 (Thermo Fisher Scientific, A32732) 1/500. Following four washes with 0.025% Triton-100/PBS, imaging was performed on an Opera Phenix Imaging System (Revvity) with a 20x water objective. All images shown are representative images. The cell number, total fluorescence and cell morphology from individual protein maker staining were analyzed using Harmony™ software (Revvity) for high-content screening systems, and graphs show the relative protein staining in *GPR34* KO compared to WT. Values represent the mean ratio(+/-S.E.), calculated from at least 6 replicated wells.

### B: Resources Table

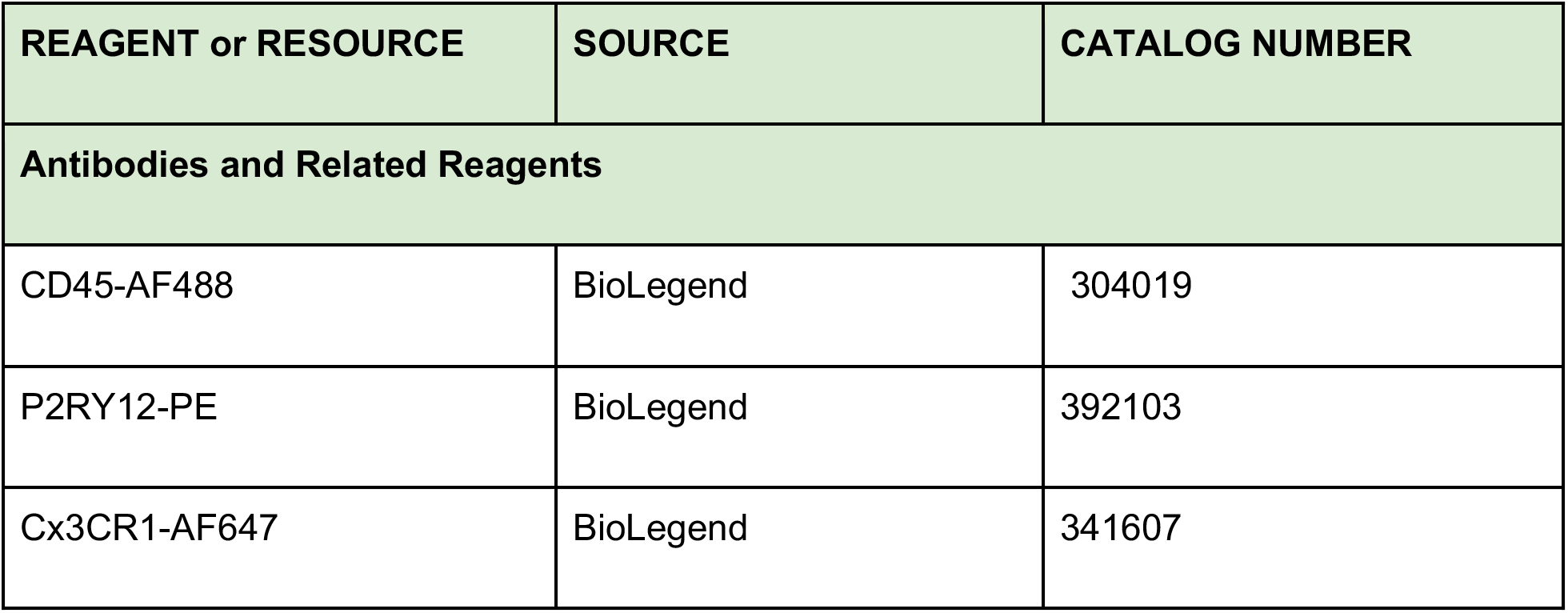
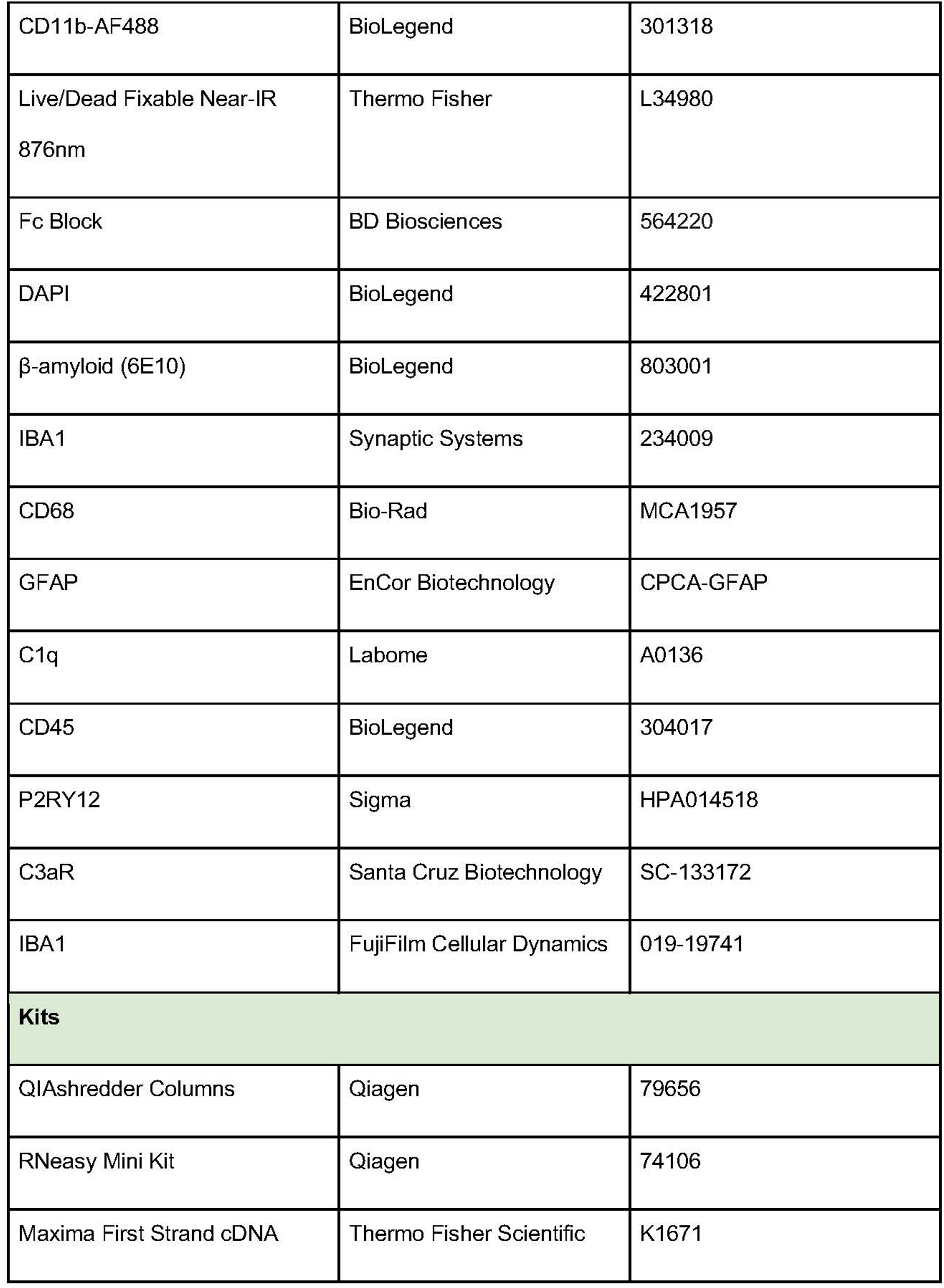
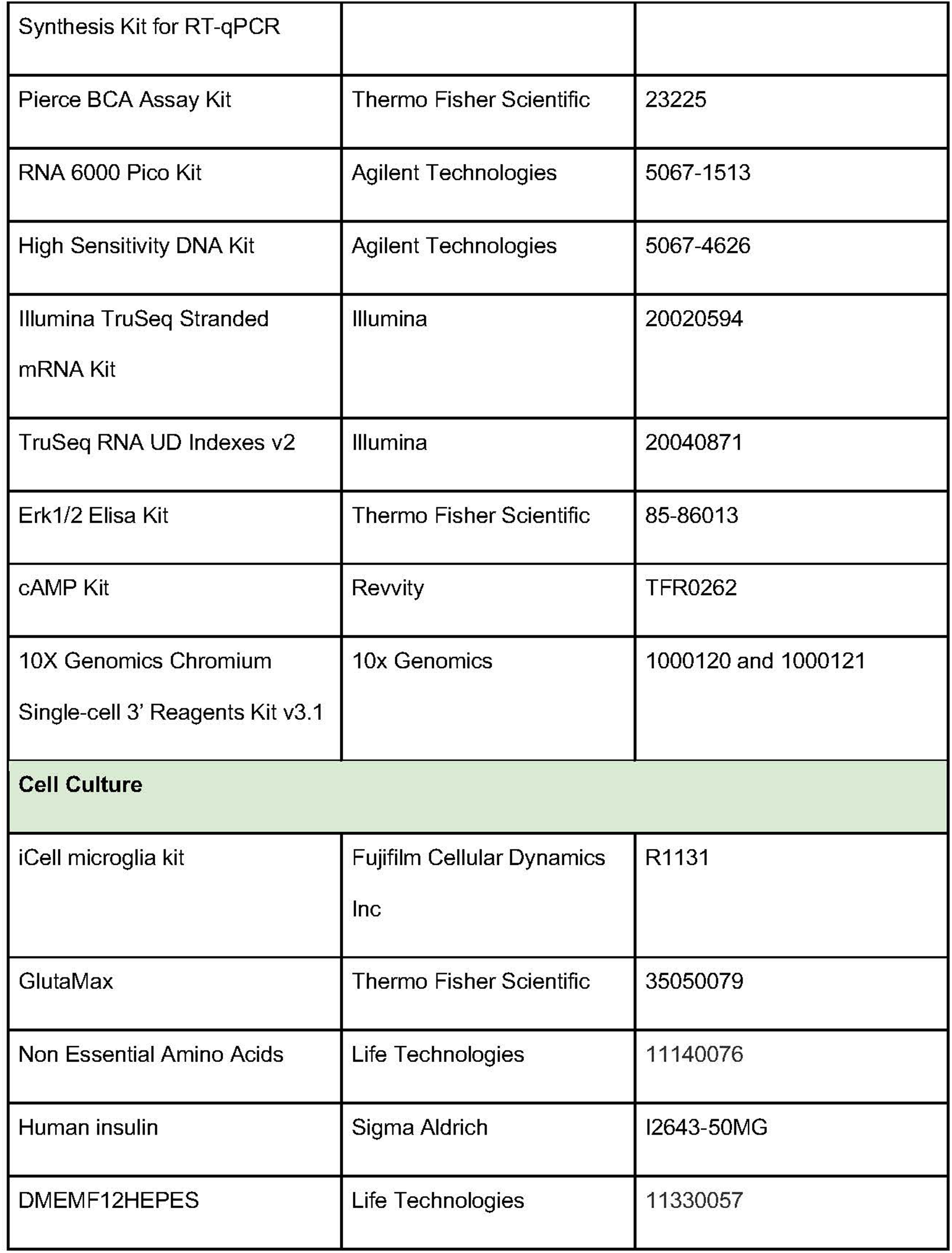
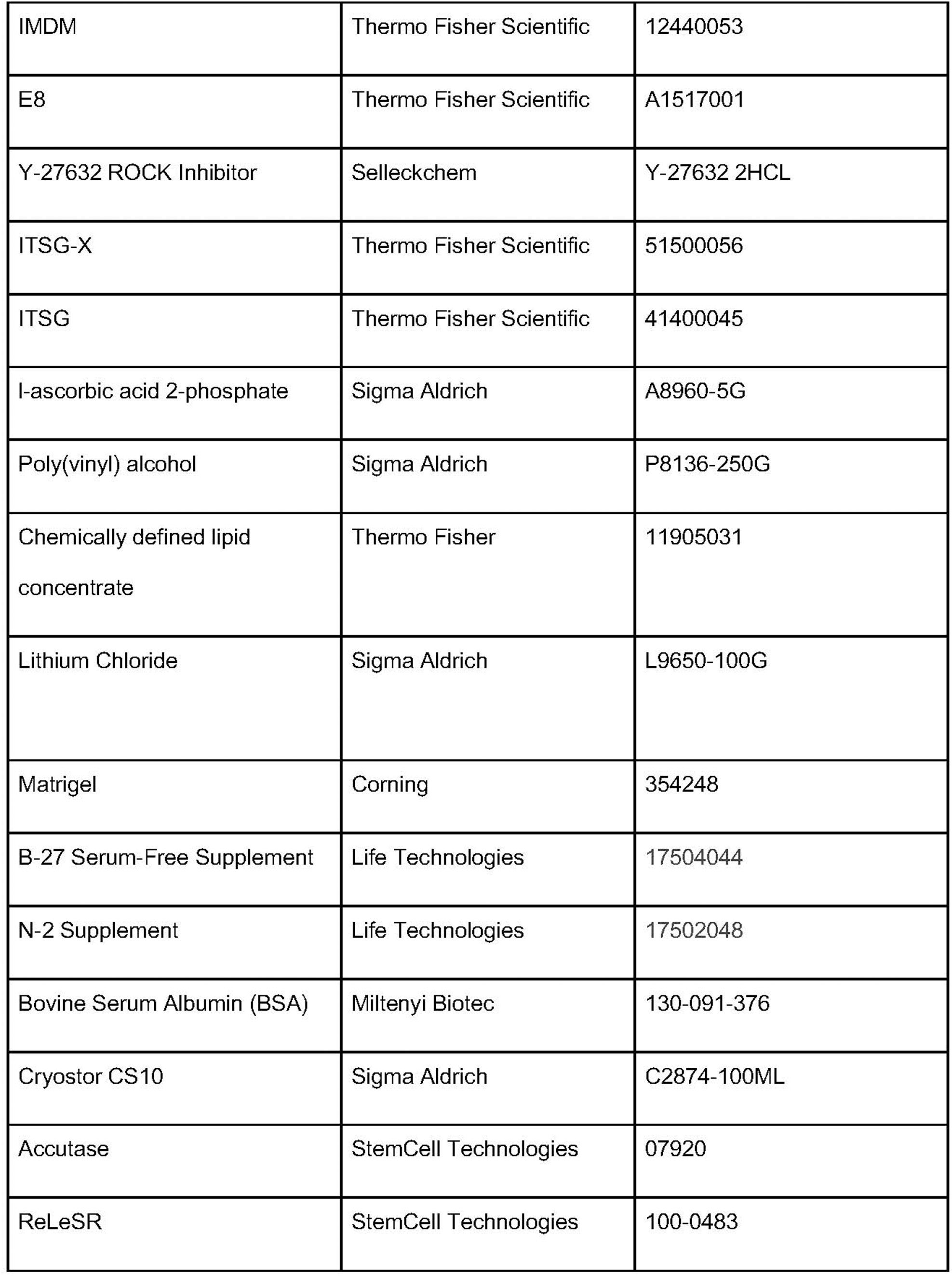
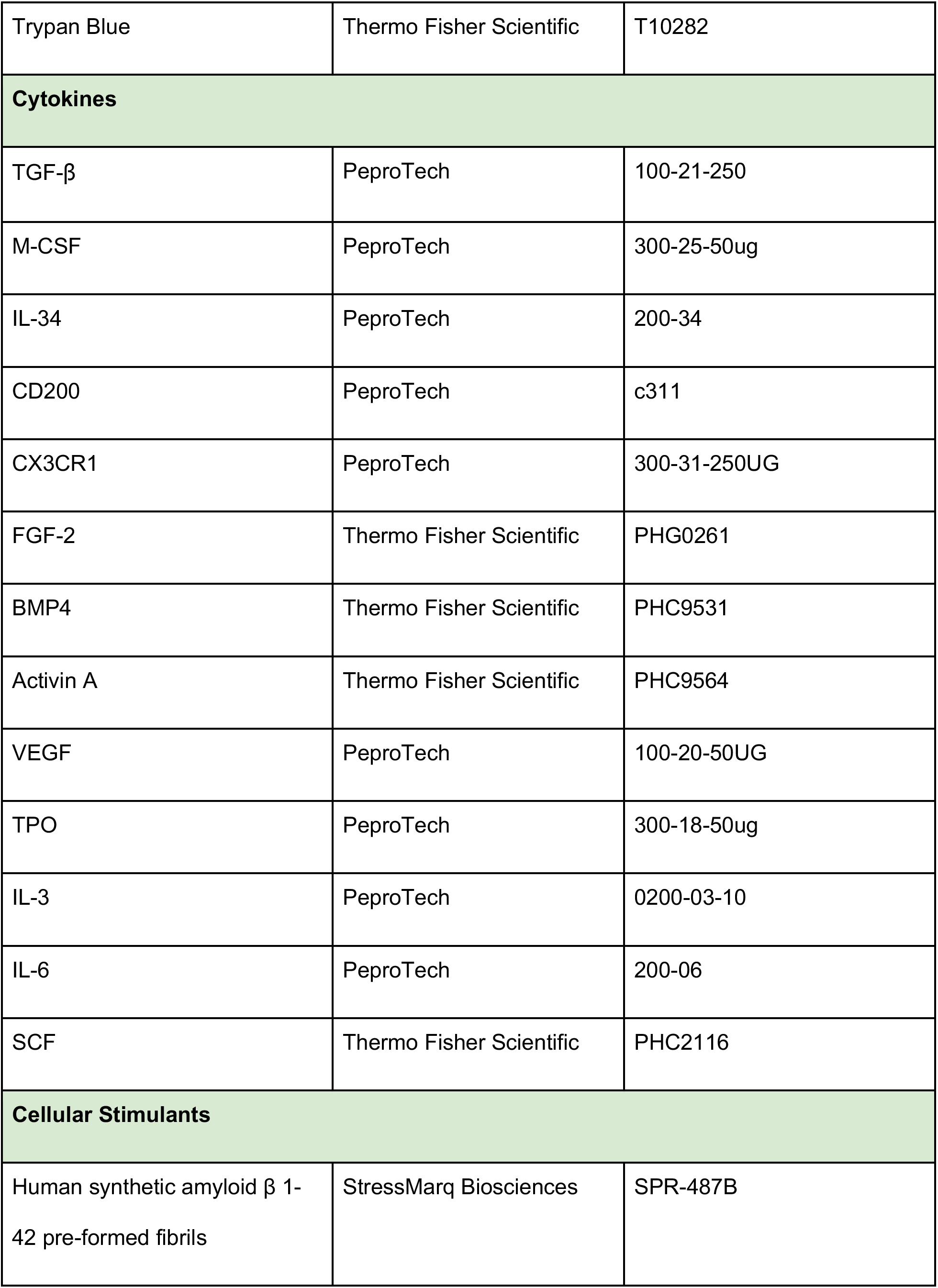
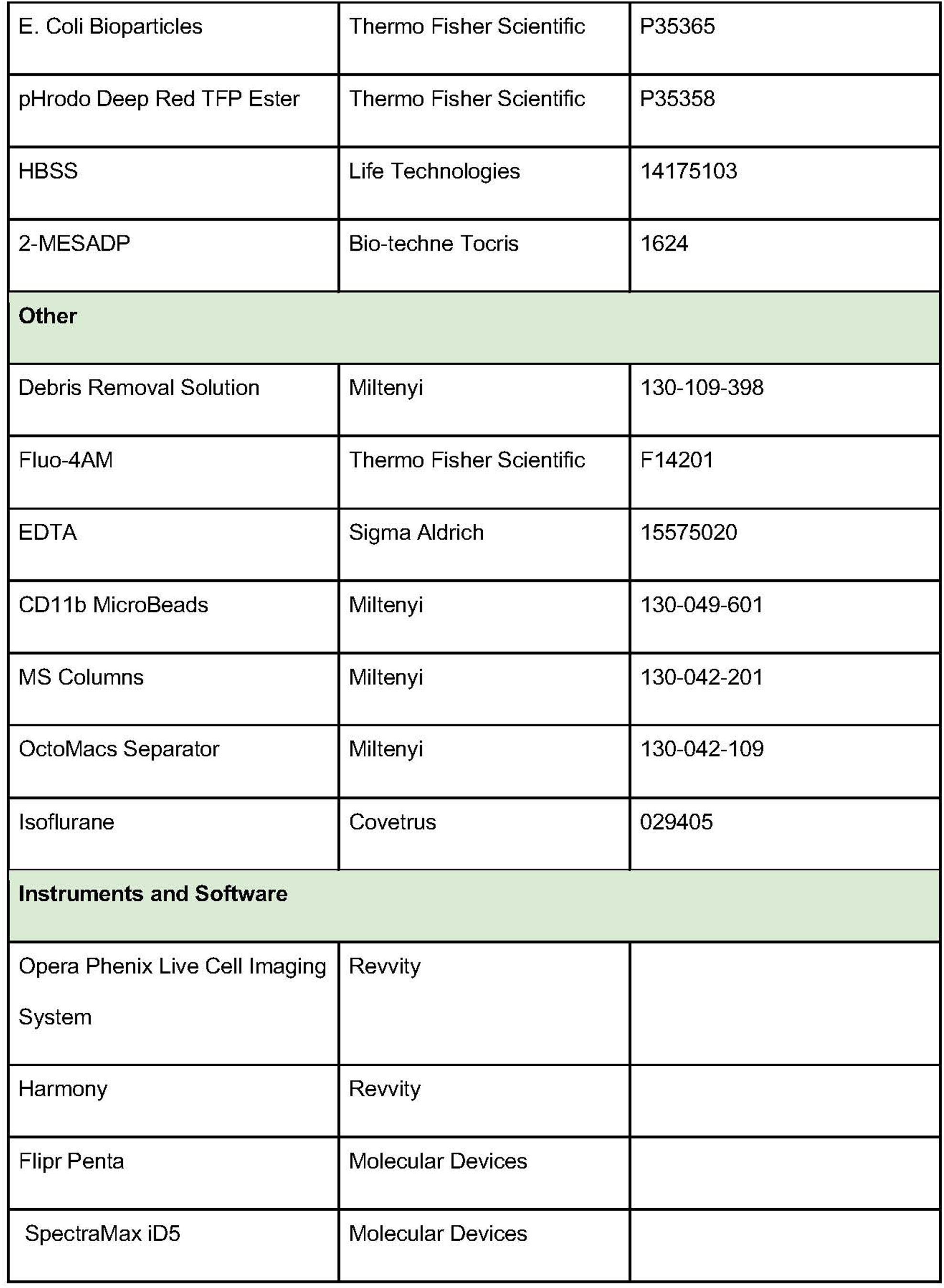
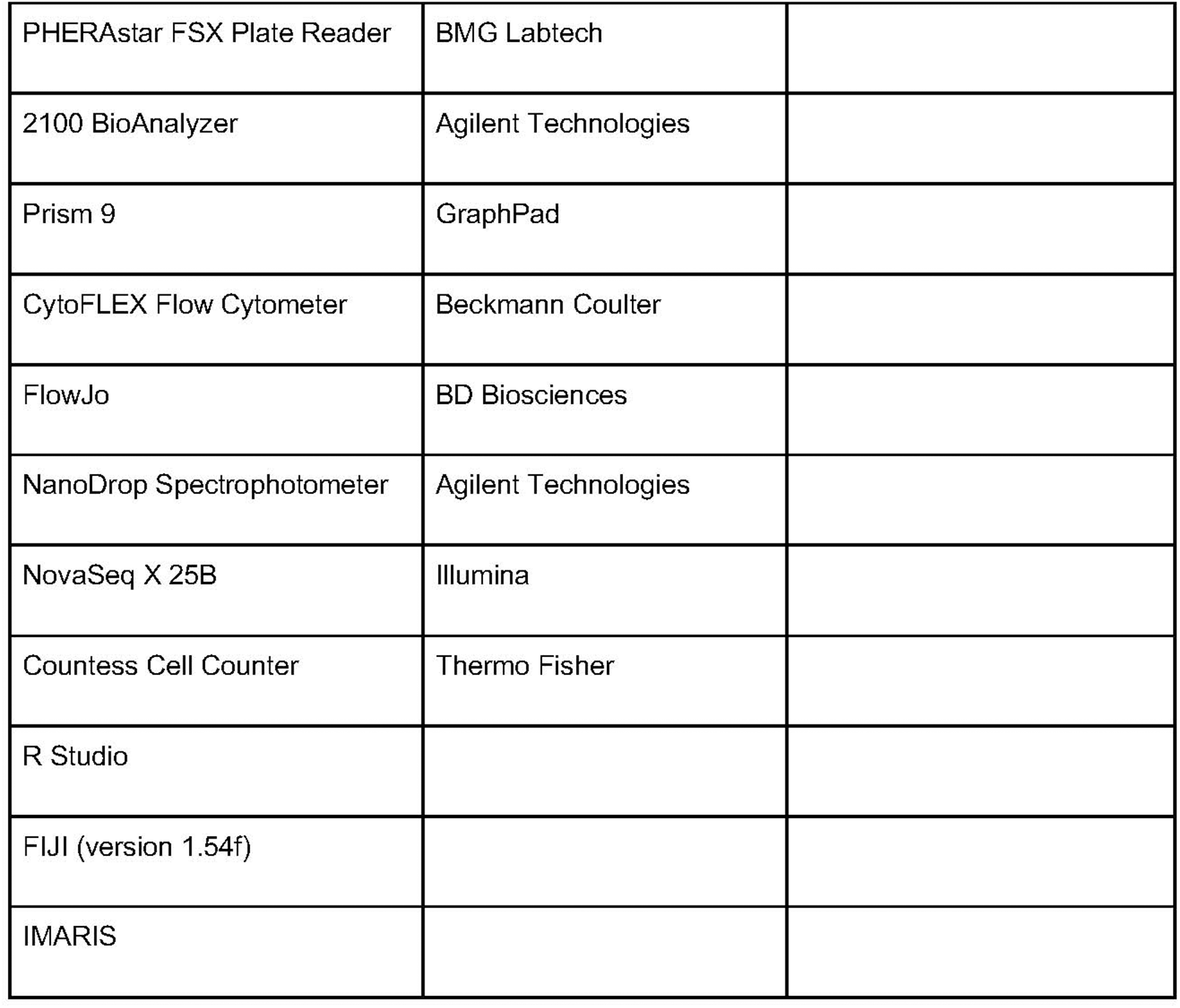

## Supplemental Figure Legends

**Supplemental Figure 1:**
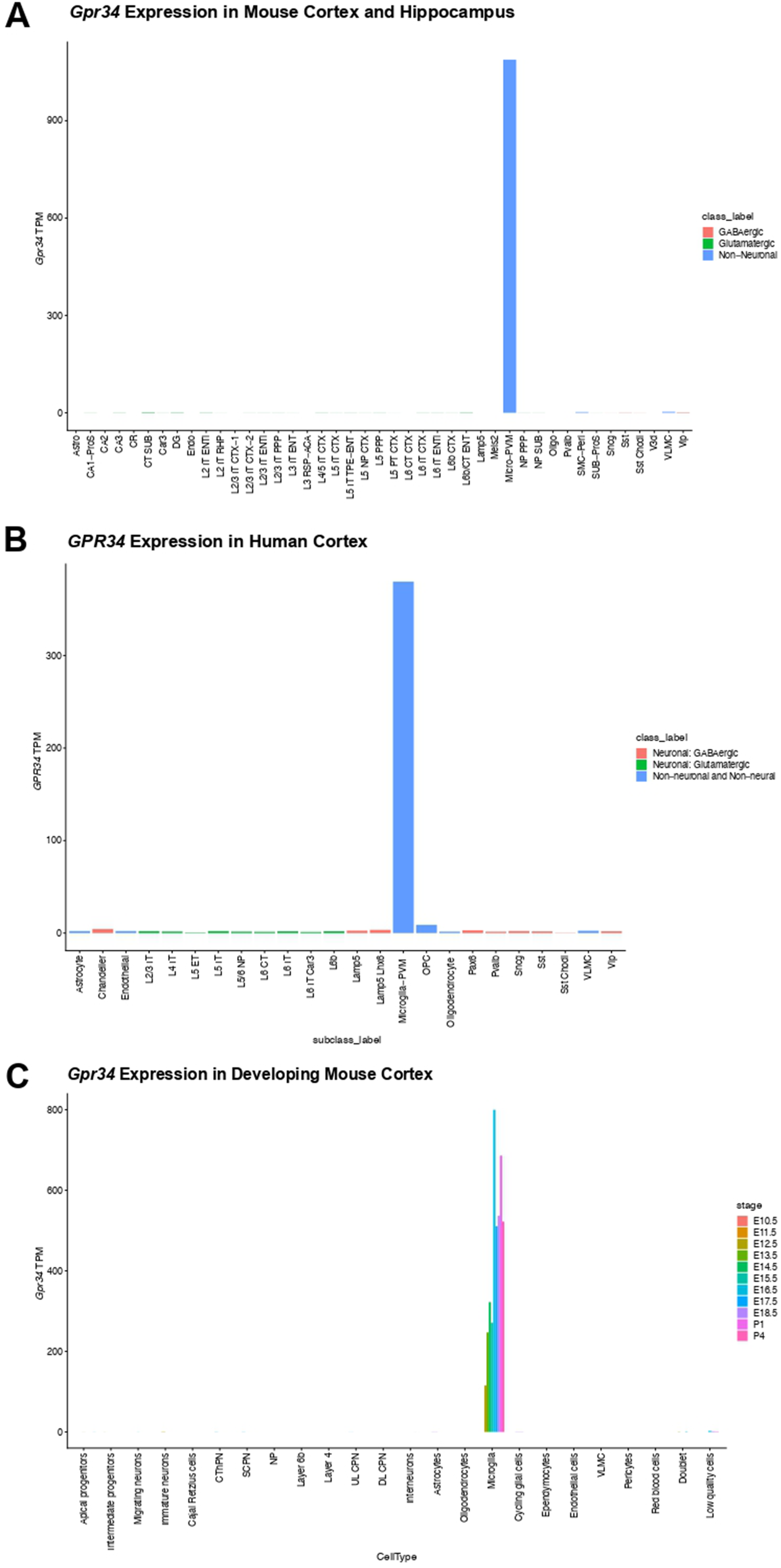
*Gpr34/GPR34* expression in mouse and human brain cell types. **A:** *Gpr34* expression (y-axis, TPM) in the mouse cortex and hippocampus based on data from the Allen Brain Cell Atlas by cell type (x-axis), colored by the class of that cell type. **B:** *GPR34* expression (y-axis, TPM) in the human cortex based on data from the Allen Brain Cell Atlas by cell type (x-axis), colored by the class of that cell type. **C:** *Gpr34* expression (y-axis, TPM) in the developing mouse cortex by cell type (x-axis) from the Allen Brain Cell Atlas, colored by age. For cell type abbreviations, please follow the source: https://celltypes.brain-map.org/

**Supplemental Figure 2:**
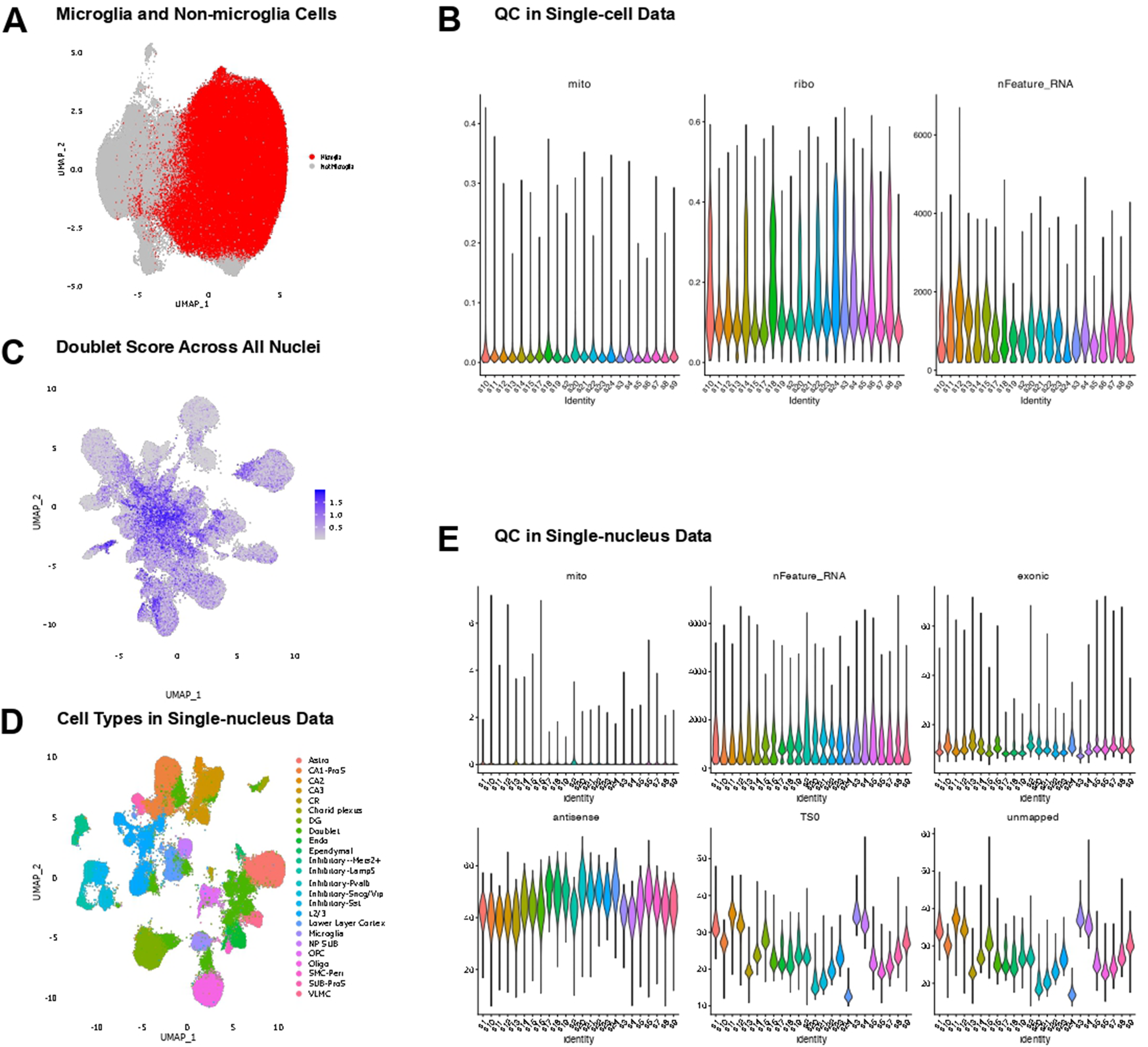
Quality control and annotation of single-cell and single-nucleus RNA-seq. **A:** UMAP plot of single-cell data, with microglia vs non-microglia cells labeled. **B:** Violin plots of different QC metrics (one per panel) for each sample (x-axis) in the single-cell data, using only the microglial cells from panel A. The QC metrics are percent mitochondrial UMI (mito), percent UMI mapping to ribosomal proteins (ribo), and number of genes per cell (nFeature_RNA). There is sample to sample variability in many of these metrics. **C:** UMAP of all droplets returned by CellRanger as nuclei in the single-nucleus dataset, colored by scds doublet score. We removed the large “blob” in the middle in downstream analysis (see Methods). **D:** UMAP of the single-nucleus data, colored by cell type. Includes doublets and cortical contamination. **E:** Violin plots of different QC metrics (one per panel) for each sample (x-axis) in the single-nucleus data after removing the central blob and doublets. The QC metrics are percent mitochondrial UMI (mito), percent UMI mapping to ribosomal proteins (ribo), and number of genes per cell (nFeature_RNA), percent reads mapping to antisense to genes (antisense), percent of reads with TSO trimmed by the Cell Ranger pipeline (TSO), and percent unmapped reads (unmapped).

**Supplemental Figure 3:**
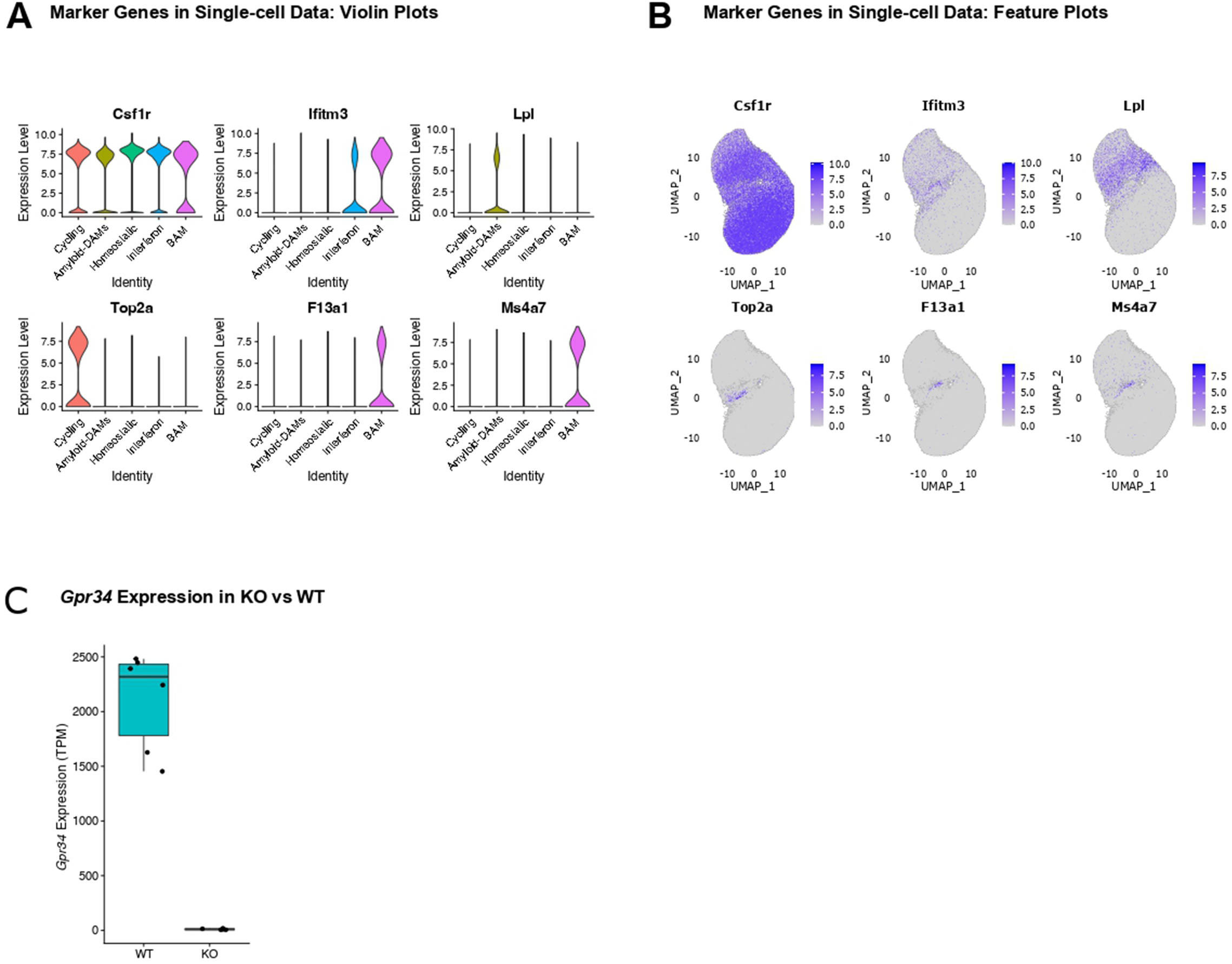
Marker expression, cell states, and *Gpr34* knockout validation in the single-cell microglia dataset. **A:** Violin plots of the expression (y-axis. CPM) in the single-cell data of various markers (different panels) in different cell types (x-axis). **B:** Feature plots of the markers from panel A, plotted on the same UMAP as in Figure 1B. **C:** Bar plot showing *Gpr34* expression in WT and *Gpr34* KO microglia

**Supplemental Figure 4:**
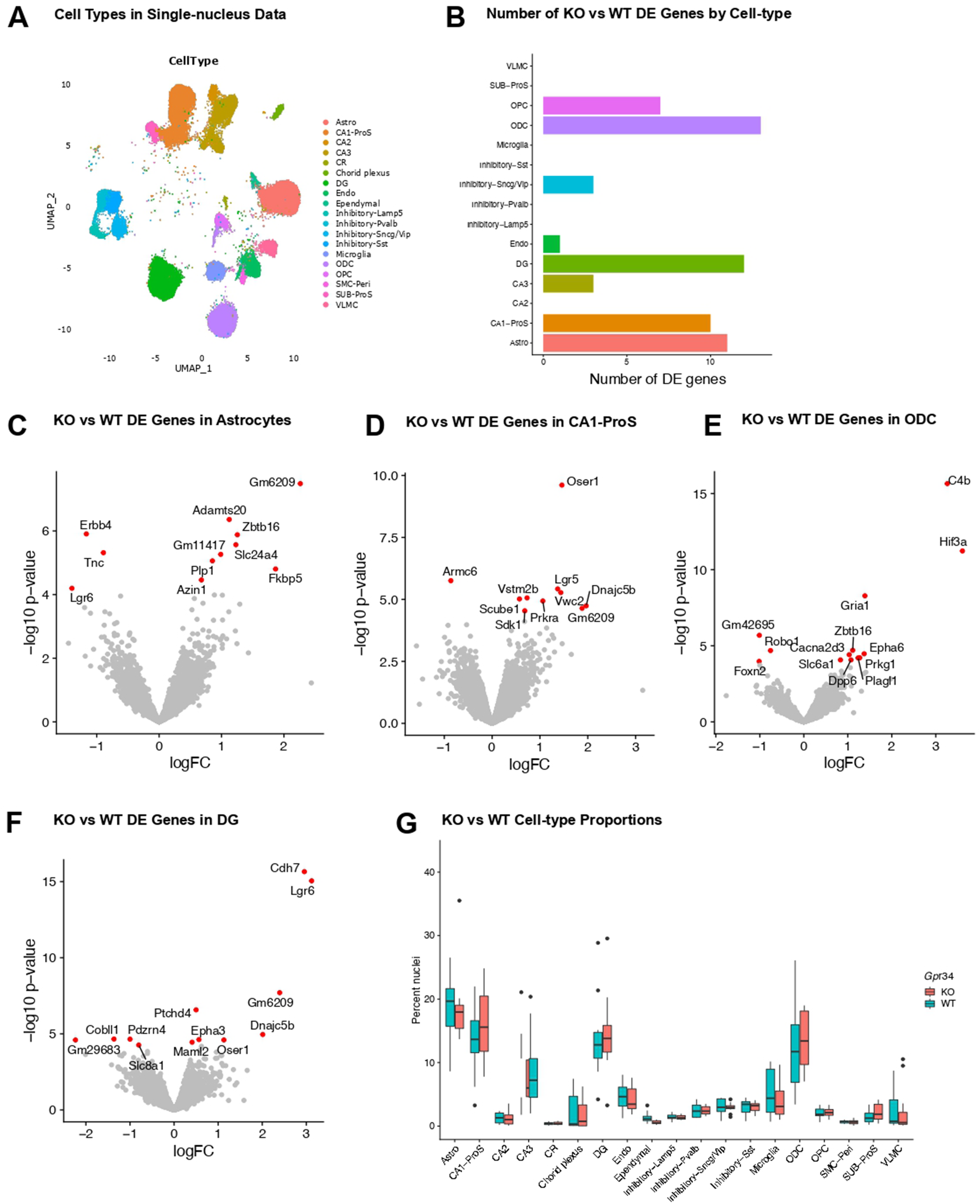
Differential gene expression and cell-type proportions in the single-nucleus dataset. **A:** UMAP of the single-nucleus data, colored by cell type. Astro is short for astrocytes and Endo is short for endothelial. **B:** Bar plot of number of DE genes (x-axis) for each cell type (y-axis) in the single-nucleus data, for *Gpr34* KO vs WT, calculated with pseudobulk analysis. **C:** Volcano plot of DE genes in *Gpr34* KO and WT mice in astrocytes, calculated with pseudobulk analysis, those colored red are significant with FDR<.05. Significant genes are labelled. The logFC is log base 2, as it is in all plots D-F. **D:** Volcano plot of DE genes in *Gpr34* KO and WT mice in CA1-ProS neurons, calculated with pseudobulk analysis, those colored red are significant. Significant genes are labelled. **E:** Volcano plot of DE genes in *Gpr34* KO and WT mice in ODCs, calculated with pseudobulk analysis, those colored red are significant. Significant genes are labelled. **F:** Volcano plot of DE genes in *Gpr34* KO and WT mice in DG neurons, calculated with pseudobulk analysis, those colored red are significant. Significant genes are labelled. **G:** Percent of nuclei in each mouse (y-axis) for each assigned cell type (x-axis) in the single-nucleus data, stratified by genotype (color). None reach significance (see Methods). In the box plots the center line is the median, the boxes span from 25% to 75%, the length of the whiskers are based off the smallest/largest value within 1.5 of the interquartile range from the median, and everything outside that range is plotted as an outlier.

**Supplemental Figure 5:**
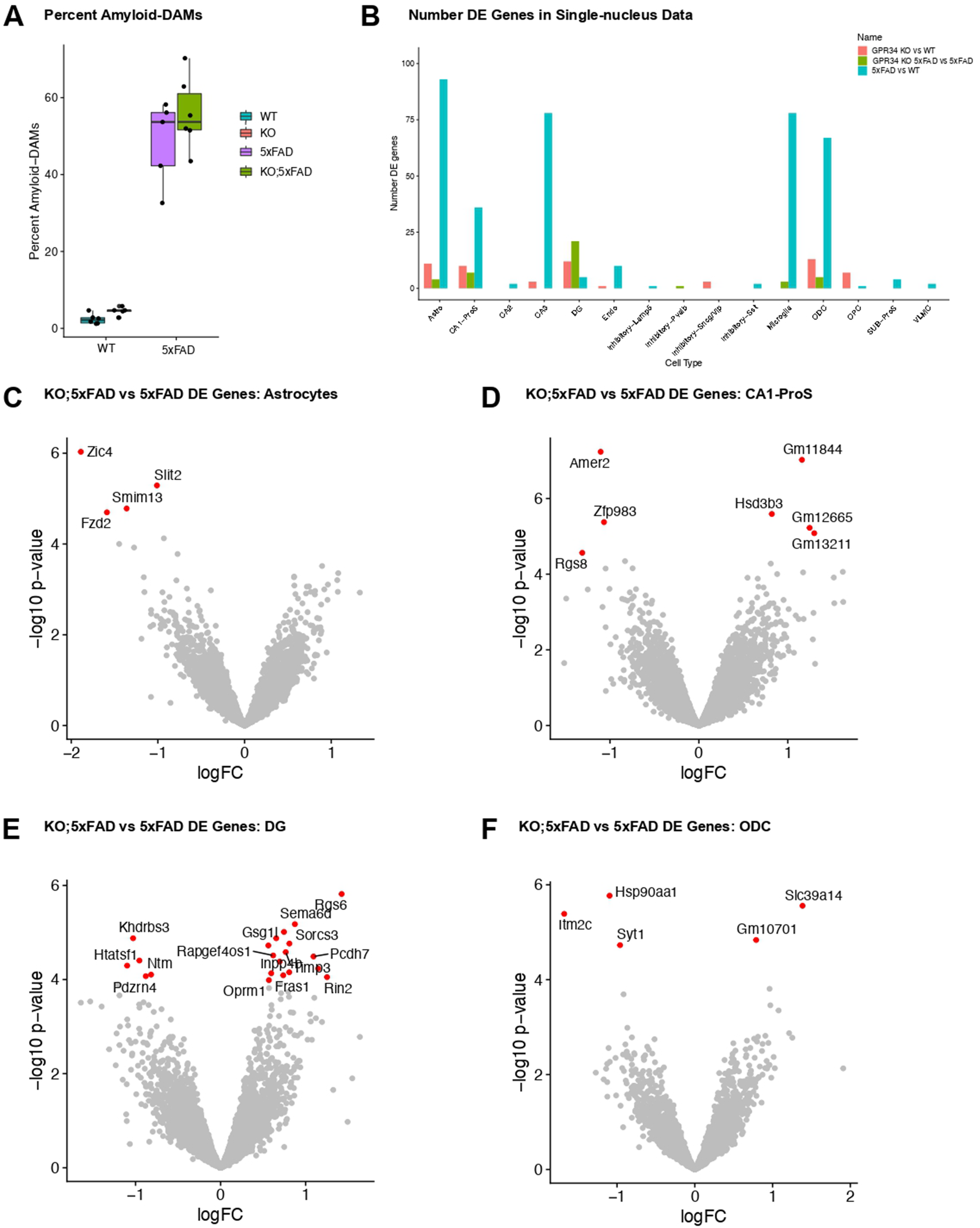
**Amyloid-DAM proportions and differential gene expression in *Gpr34*- and 5xFAD-related comparisons**. **A:** Percent of DAM cells (y-axis) in 5xFAD and non-5xFAD mice (x-axis) in each *Gpr34* KO and WT mouse (color), based on the single-cell microglia data. There is a significant difference (FDR<.05) between KO and WT as well as between 5xFAD and WT and between KO;5xFAD and KO. **B:** The number of DE genes (y-axis, based on FDR<.05) for each cell type in the single-nucleus data (x-axis) for each comparison (color). **C:** Volcano plot of DE genes for astrocytes in the single-nucleus data for *Gpr34* KO;5xFAD versus 5xFAD mice. Those colored red and with labels are significant (FDR<.05). The logFC is log base 2, as it is in the volcano plots in D-F. **D:** Volcano plot of DE genes for CA1-ProS neurons in the single-nucleus data for *Gpr34* KO;5xFAD versus 5xFAD mice. Those in red and with labels are significant (FDR<.05). **E:** Volcano plot of DE genes for DG neurons in the single-nucleus data for *Gpr34* KO;5xFAD versus 5xFAD mice. Those colored red and with labels are significant (FDR<.05). **F:** Volcano plot for ODCs in the single-nucleus data for KO;5xFAD versus 5xFAD mice. Those colored red and with labels are significant (FDR<.05).

**Supplemental Figure 6:**
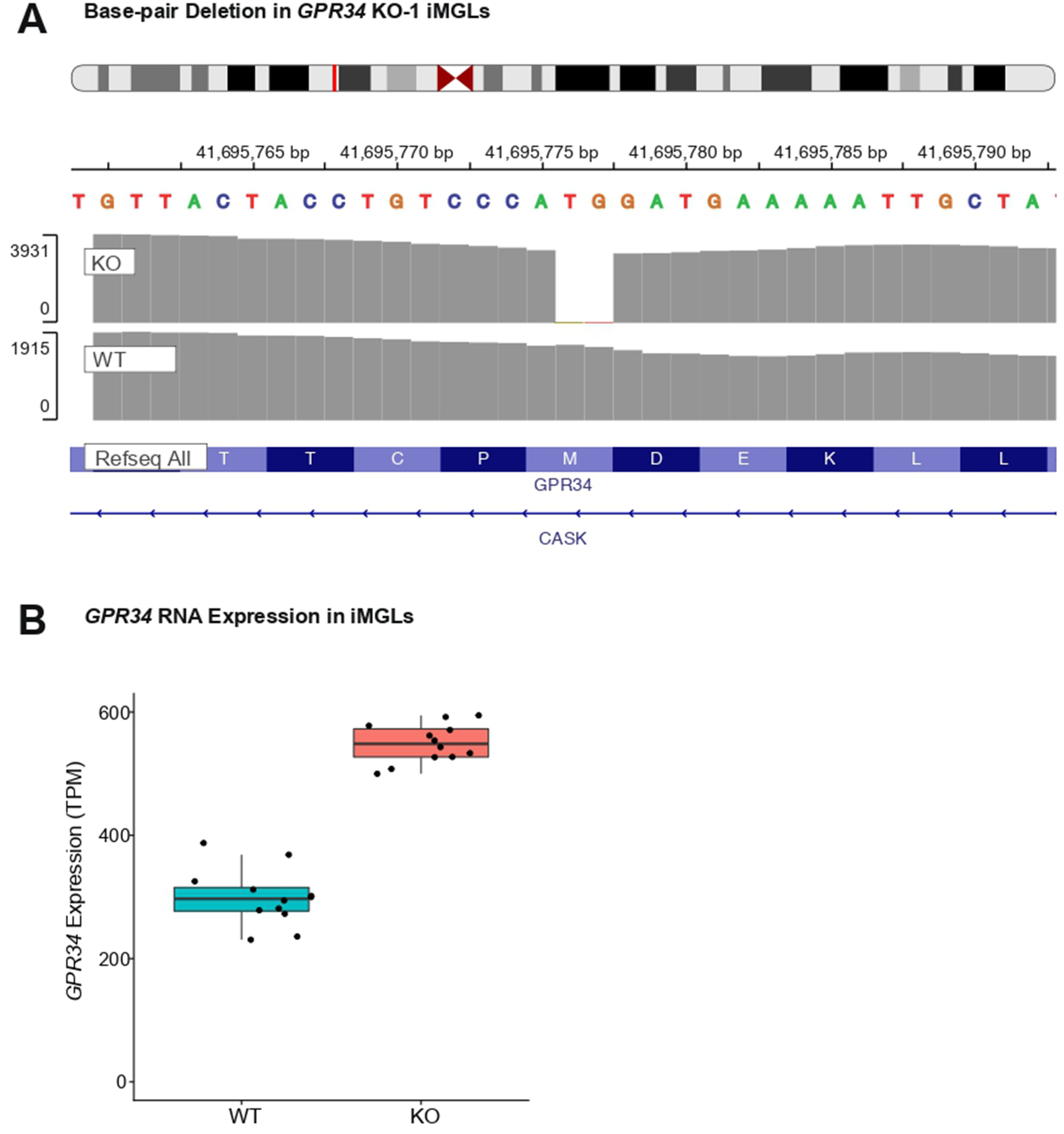
*GPR34* Knockout Validation in Human iMGLs. **A:** Integrative Genomics Viewer (IGV) shows the exact base-pair deletion in the *GPR34* KO-1 iMGL line. Y-axis shows the # of reads at each position. **B:** Expression of *GPR34* RNA (y-axis, TPM) in *GPR34* KO and WT iMGLs (x-axis).

**Supplemental Figure 7:**
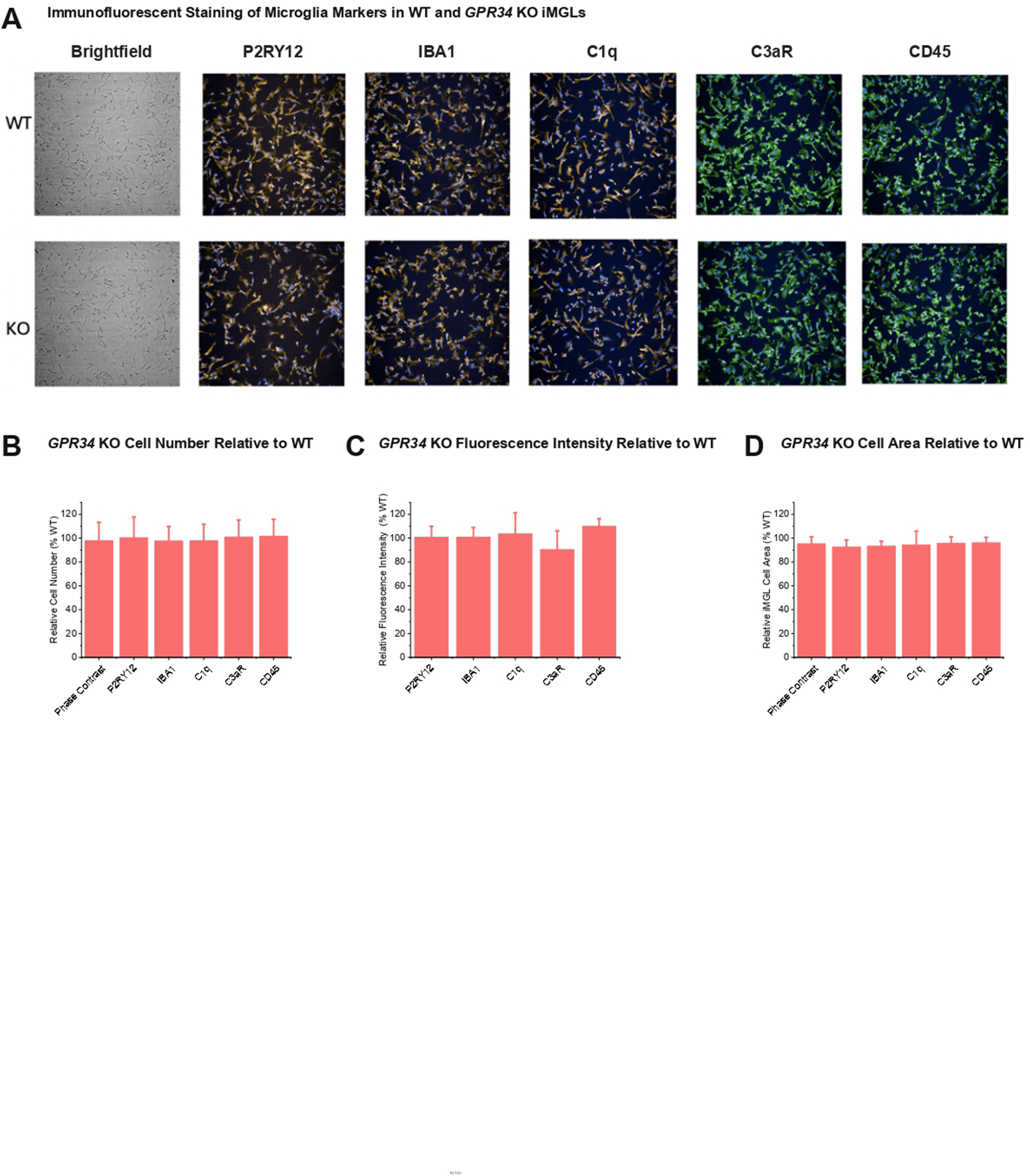
Immunofluorescent validation of microglial marker expression in WT and *GPR34 KO* iMGLs. **A:** Representative brightfield and immunofluorescence staining images of microglia markers (P2RY12, IBA1, C1q, C3aR, CD45) in WT and *GPR34* KO iMGLs. Nuclei are shown in blue; marker staining is shown in green or yellow/orange; brightfield images are grayscale. **B:** Quantification of cell number in *GPR34* KO iMGLs relative to WT iMGLs across markers. Values are shown as percent WT. **C:** Quantification of fluorescence intensity in *GPR34* KO iMGLs relative to WT across markers. Values are shown as percent WT. **D:** Quantification of cell area in *GPR34* KO iMGLs relative to WT across markers. Values are shown as percent WT. Values represent the mean ratio(+/-S.E.), calculated from at least 6 replicated wells.

**Supplemental Figure 8:**
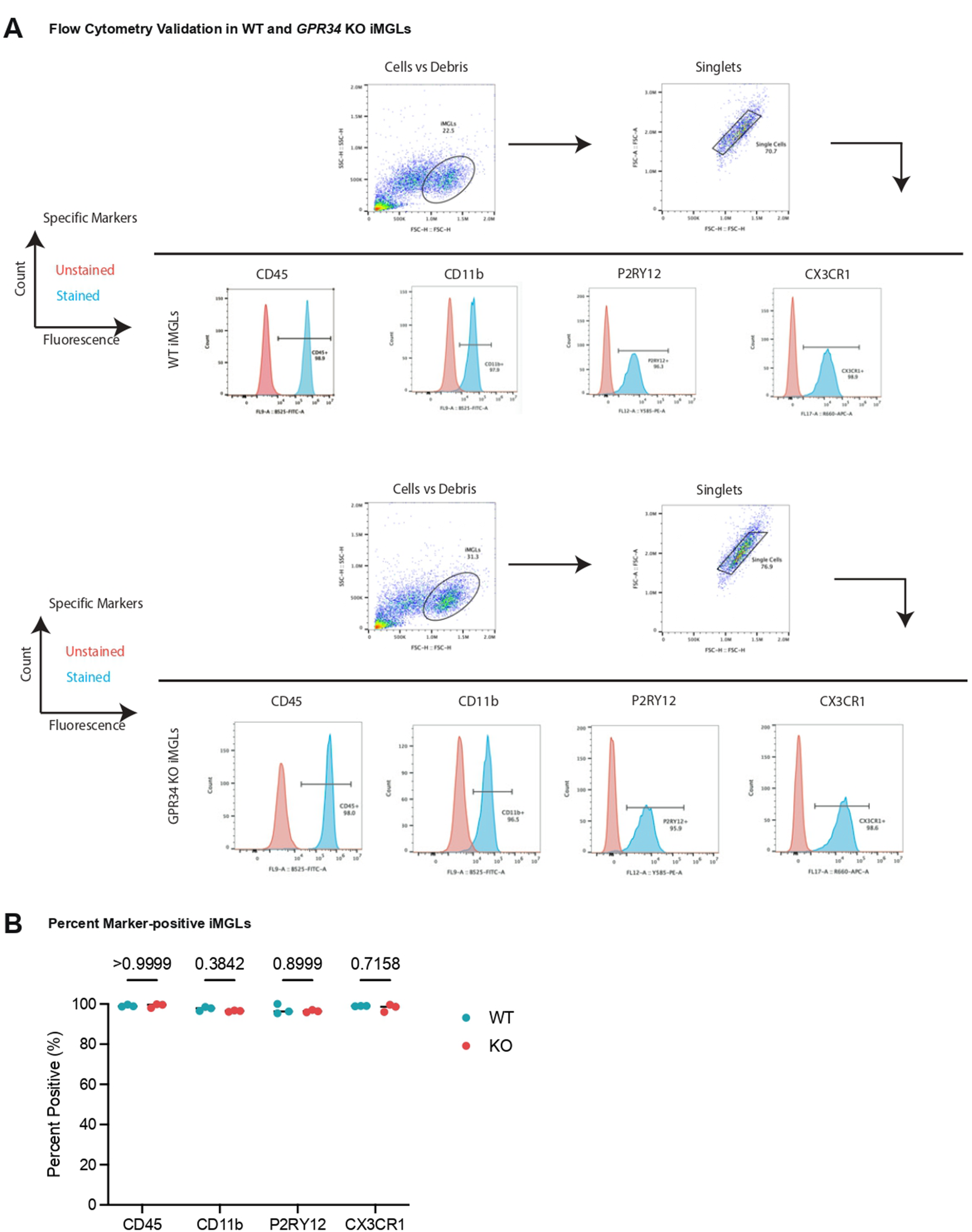
Flow cytometry validation of microglial marker expression in WT and *GPR34* KO iMGLs. **A:** Flow cytometry gating strategy and representative histograms illustrating expression of microglial markers (CD45, CD11b, P2RY12, and CX3CR1) in WT and *GPR34* KO Day 40 iMGLs. The x-axis denotes fluorescence intensity in the indicated detection channels (FL9-B525 FITC, FL12-Y585 PE, or FL17-A R660 APC-A), and the y-axis denotes cell count. Unstained controls were used to define background fluorescence and gating thresholds. Horizontal lines indicate the gated marker-positive populations. **B:** Quantification of the percentage of marker-positive iMGLs for CD45, CD11b, P2RY12, and CX3CR1 across three independent biological replicates. No significant differences were detected between WT and *GPR34* KO iMGLs by two-way ANOVA. P-values shown correspond to genotype effects.

**Supplemental Figure 9:**
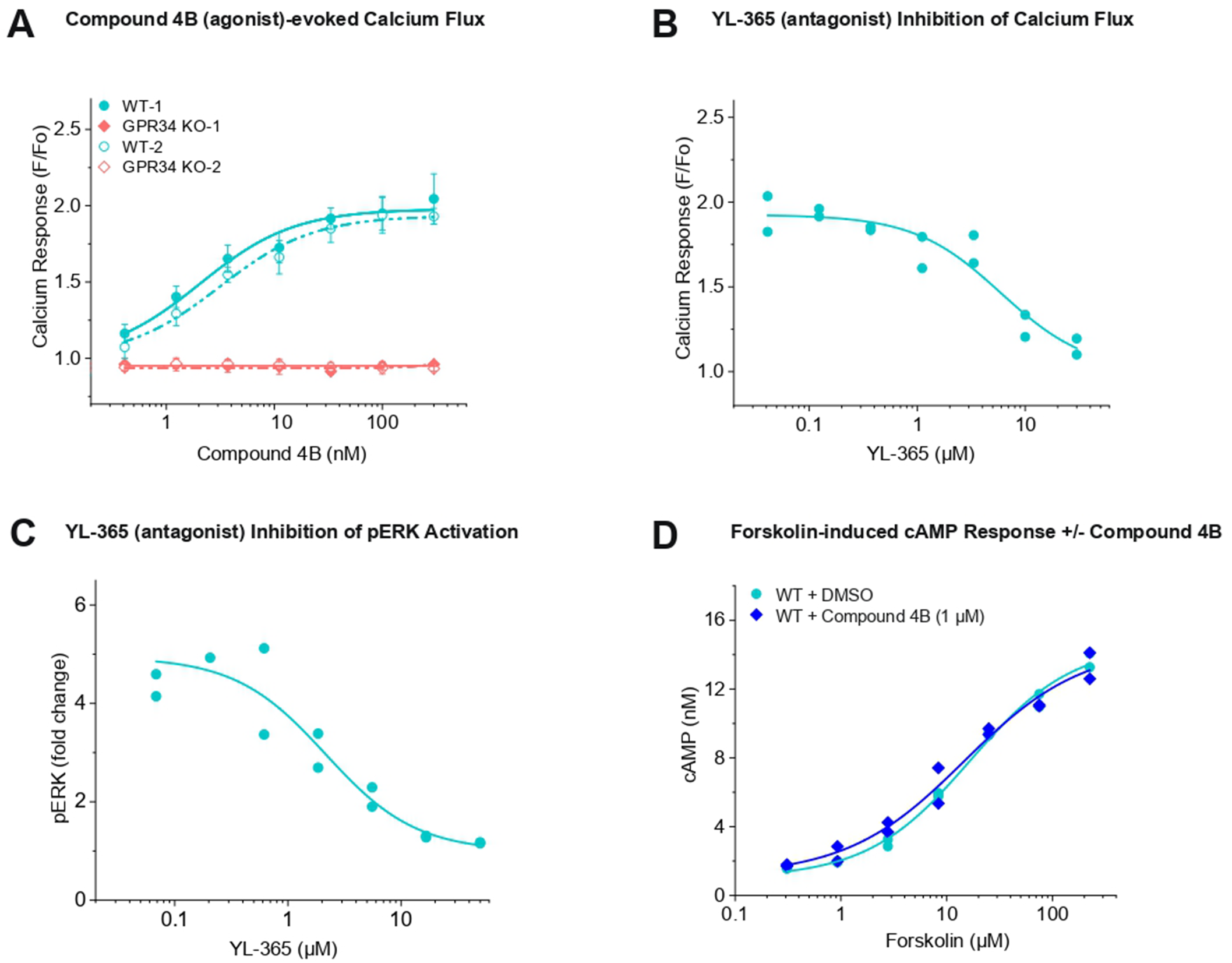
Compound 4B–evoked GPR34 signaling and YL-365 antagonism in iMGLs. **A:** Dose-dependent calcium response to the GPR34 agonist Compound 4B in WT and two independent *GPR34* KO lines, KO-1 and KO-2. **B:** The GPR34 antagonist YL-365 inhibits Compound-4B-evoked calcium flux in WT iMGLs. Duplicates **C:** YL-365 dose-response inhibition of Compound-4B-evoked pERK activation in WT iMGLs. Duplicates **D:** Forskolin-induced cAMP production in WT iMGLs treated with Compound 4B and DMSO. Sample sizes and stats: *GPR34* KO line 1 (KO-1) and its control (WT-1) were used for B-D. The representative curve for A was shown from the average data are from triplicates and for B-D is from duplicates.

**Supplemental Figure 10:**
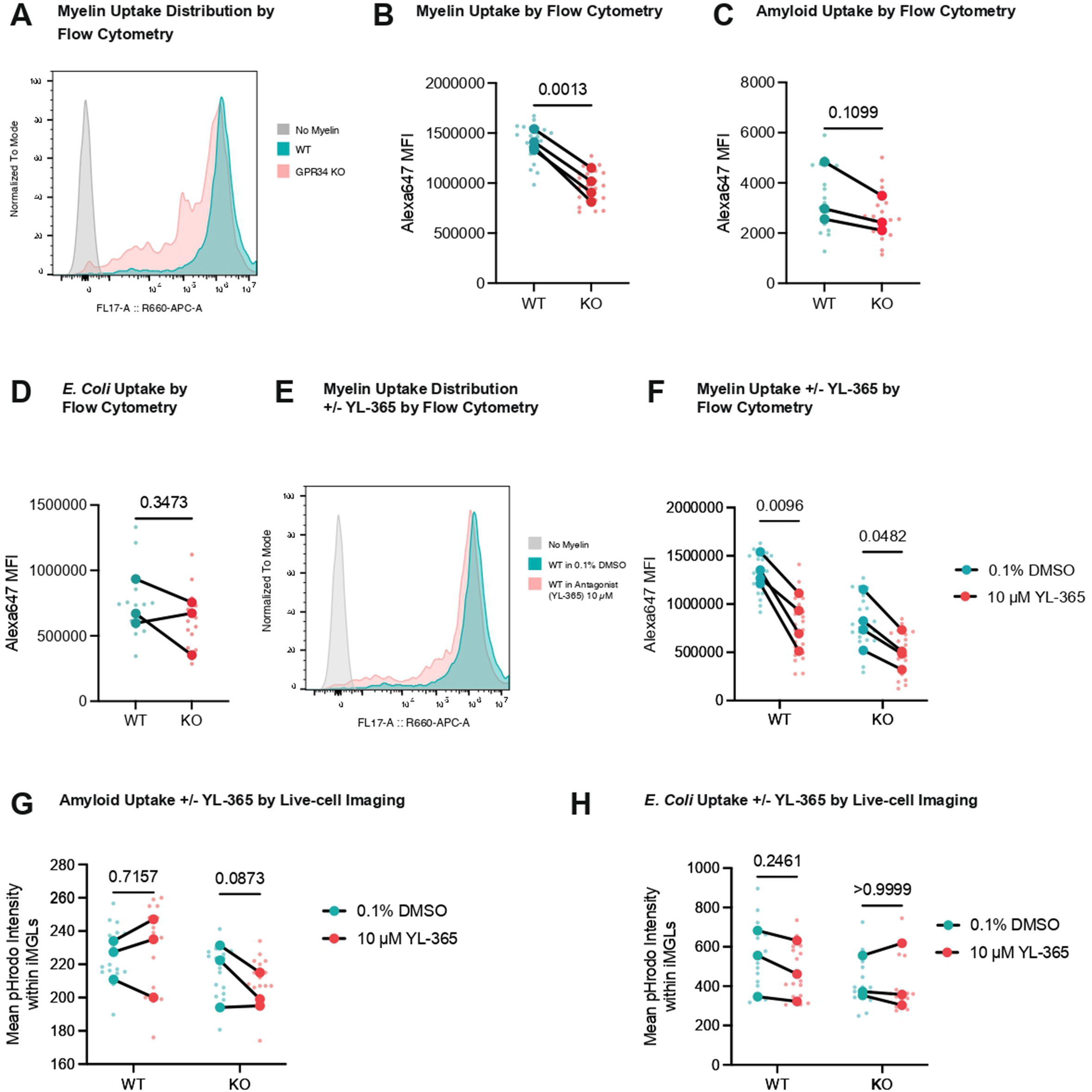
GPR34 loss or antagonism selectively reduces myelin uptake in iMGLs. **A:** Flow cytometry histogram traces normalized to mode from WT and *GPR34* KO iMGLs exposed to pHrodo AF647-labeled myelin for 25 h. FL17-A R660 APC-A denotes the APC-compatible fluorescence detection channel used to measure pHrodo AF647 signal. **B:** Quantification of median fluorescence intensity (MFI) upon myelin treatment by flow cytometry in WT and *GPR34* KO iMGLs. **C:** Quantification of MFI upon amyloid fibrils treatment by flow cytometry in WT and *GPR34* KO iMGLs. **D:** Quantification of MFI upon *E. coli* bioparticles treatment by flow cytometry in WT and *GPR34* KO iMGLs. **E:** Flow cytometry histogram traces normalized to mode from WT iMGLs treated with the GPR34 antagonist YL-365 (10 µM) or vehicle control (0.1% DMSO) and exposed to pHrodo AF647-labeled myelin for 25 h. FL17-A R660 APC-A denotes the APC-compatible fluorescence detection channel used to measure pHrodo AF647 signal. **F:** Quantification of MFI in WT and *GPR34* KO iMGLs treated with vehicle or YL-365 after 25-hour myelin treatment. **G-H:** Live-cell imaging quantification of pHrodo signal in iMGLs treated with vehicle or YL-365 after 24-hour exposure to pHrodo-labeled amyloid fibrils (**G**) or *E. coli* bioparticles **(H).** Data are represented as mean values with +/- SEM. Smaller circles represent technical replicates, and larger circles represent biological replicates. Statistical tests were undertaken at 5h, 24h, and 25h (n=2-4 independent biological replicates with 5-6 technical replicates per condition). Two-way ANOVA with Bonferroni correction, multiple comparisons test, paired t-test with Welch’s correction, as appropriate. Lines connect paired WT and KO biological replicates derived from the same experimental batch.

**Supplemental Figure 11:**
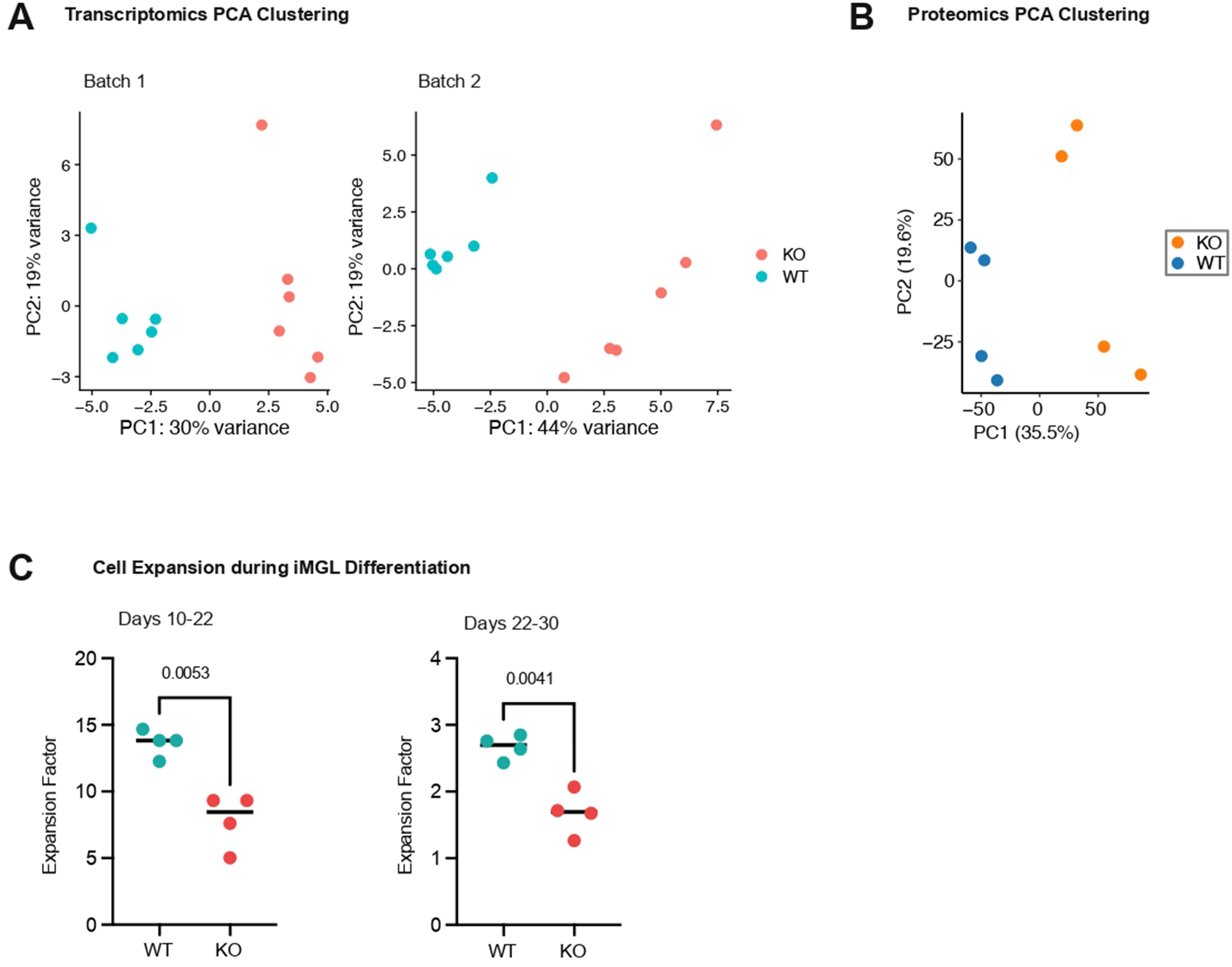
Transcriptomic, proteomic, and proliferation characterization of WT and *GPR34 KO* IMGLs. **A:** PCA plots of iMGL bulk RNA-seq samples from two different batches, colored by genotype. **B:** PCA plot of quantitative MS/MS proteomics showing distinct clustering of WT and *GPR34* KO samples **C:** Quantification of cell expansion during iMGL differentiation, showing reduced expansion of GPR34 KO cells compared to WT controls (left: days 10–22 and right: days 22–30). Expansion factor was calculated as the cell count at the time of replating divided by the initial number of cells seeded for differentiation. Paired t-test with Welch’s correction.

**Supplemental Figure 12:**
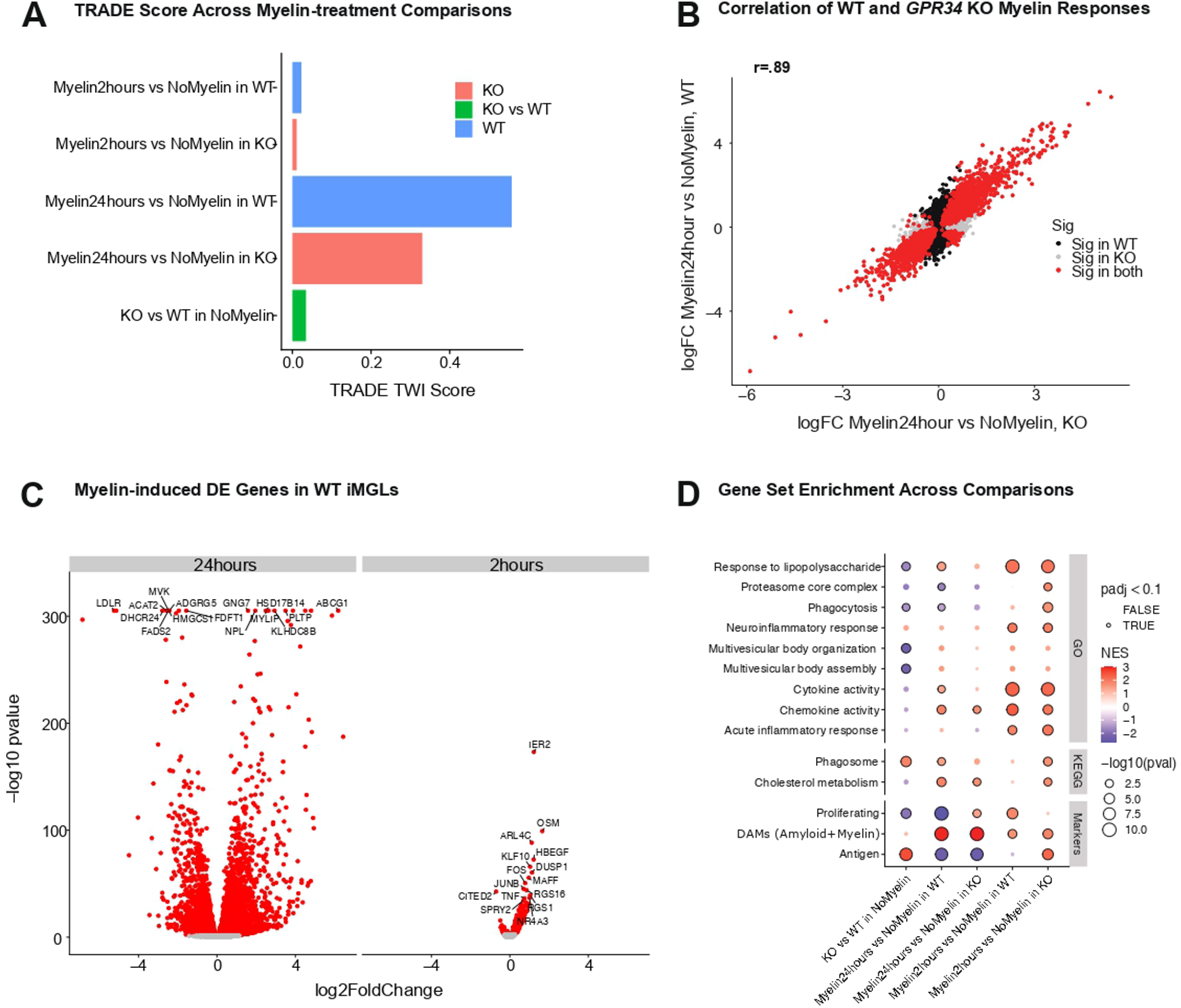
Transcriptional responses to myelin treatment in WT and *GPR34* KO iMGLs. **A:** Bar plots of TRADE score (x-axis) for each comparison (y-axis). **B:** Scatter plot comparing the logFC values (log base 2) for 24-hour myelin vs no-myelin in *GPR34* KO (x-axis) and WT (y-axis) iMGLs. Only genes that are significant in at least one of the two are shown. Colored based on which comparison is significant. Pearson correlation is reported. **C:** Volcano plots of DE genes in WT iMGLs following 24-hour or 2-hour myelin treatment vs no myelin control. Significant hits (FDR<.05) are colored red. Some top genes (chosen based off p-value) are labelled. The logFC is log base 2. **D:** Gene set enrichment analysis (y-axis) across WT and *GPR34* KO iMGL comparisons (x-axis). Color is NES, size of -log_10_ p-value, those circled in black are significant.

## References

1. Depp, C. et al. Myelin dysfunction drives amyloid-β deposition in models of Alzheimer’s disease. Nature 618, 349–357 (2023).

2. Bohlen, C. J., Friedman, B. A., Dejanovic, B. & Sheng, M. Microglia in Brain Development, Homeostasis, and Neurodegeneration. Annu. Rev. Genet. 53, 1–26 (2019).

3. Badimon, A. et al. Negative feedback control of neuronal activity by microglia. Nature 586, 417–423 (2020).

4. Paolicelli, R. C. et al. Microglia states and nomenclature: A field at its crossroads. Neuron 110, 3458–3483 (2022).

5. Hansen, D. V., Hanson, J. E. & Sheng, M. Microglia in Alzheimer’s disease. J. Cell Biol. 217, 459–472 (2018).

6. Salter, M. W. & Stevens, B. Microglia emerge as central players in brain disease. Nat. Med. 23, 1018–1027 (2017).

7. Depp, C., Doman, J. L., Hingerl, M., Xia, J. & Stevens, B. Microglia transcriptional states and their functional significance: Context drives diversity. Immunity 58, 1052–1067 (2025).

8. Deczkowska, A. et al. Disease-Associated Microglia: A Universal Immune Sensor of Neurodegeneration. Cell 173, 1073–1081 (2018).

9. Keren-Shaul, H. et al. A Unique Microglia Type Associated with Restricting Development of Alzheimer’s Disease. Cell 169, 1276–1290.e17 (2017).

10. Bellenguez, C. et al. New insights into the genetic etiology of Alzheimer’s disease and related dementias. Nat. Genet. 54, 412–436 (2022).

11. (ADGC), A. D. G. C., et al. Genetic meta-analysis of diagnosed Alzheimer’s disease identifies new risk loci and implicates Aβ, tau, immunity and lipid processing. Nat. Genet. 51, 414–430 (2019).

12. Haque, Md. E., Kim, I.-S., Jakaria, Md., Akther, M. & Choi, D.-K. Importance of GPCR-Mediated Microglial Activation in Alzheimer’s Disease. Front. Cell. Neurosci. 12, 258 (2018).

13. Hsiao, C.-C. et al. GPCRomics of Homeostatic and Disease-Associated Human Microglia. Front. Immunol. 12, 674189 (2021).

14. Schöneberg, T., Meister, J., Knierim, A. B. & Schulz, A. The G protein-coupled receptor GPR34 – The past 20 years of a grownup. Pharmacol. Ther. 189, 71–88 (2018).

15. Krasemann, S. et al. The TREM2-APOE Pathway Drives the Transcriptional Phenotype of Dysfunctional Microglia in Neurodegenerative Diseases. Immunity 47, 566–581.e9 (2017).

16. Lin, L.-L. et al. GPR34 Knockdown Relieves Cognitive Deficits and Suppresses Neuroinflammation in Alzheimer’s Disease via the ERK/NF-κB Signal. Neuroscience 528, 129–139 (2023).

17. Liebscher, I. et al. Altered Immune Response in Mice Deficient for the G Protein-coupled Receptor GPR34*. J. Biol. Chem. 286, 2101–2110 (2011).

18. Lin, B. et al. GPR34 senses demyelination to promote neuroinflammation and pathologies. Cell. Mol. Immunol. 21, 1131–1144 (2024).

19. Preissler, J. et al. Altered microglial phagocytosis in GPR34 - deficient mice. Glia 63, 206–215 (2015).

20. Raju, K. et al. GPR34 regulates microglia state and loss-of-function rescues TREM2 metabolic dysfunction. bioRxiv 2025.03.28.646038 (2025) doi:10.1101/2025.03.28.646038.

21. Zhou, Y. et al. Demyelination-derived lysophosphatidylserine promotes microglial dysfunction and neuropathology in a mouse model of Alzheimer’s disease. Cell. Mol. Immunol. 22, 134–149 (2025).

22. Bohannon, D. G. et al. Impaired removal of dying brain cells by microglia in Gpr34 deficient mice. J. Neuroinflammation 10.1186/s12974-026-03777-4 (2026) doi:10.1186/s12974-026-03777-4.

23. Etani, H. et al. Selective agonism of GPR34 stimulates microglial uptake and clearance of amyloid β fibrils. Alzheimer’s Res. Ther. 17, 248 (2025).

24. Iida, Y. et al. Lysophosphatidylserine stimulates chemotactic migration of colorectal cancer cells through GPR34 and PI3K/Akt pathway. Anticancer Res. 34, 5465–72 (2014).

25. Omi, J., Kano, K. & Aoki, J. Current Knowledge on the Biology of Lysophosphatidylserine as an Emerging Bioactive Lipid. Cell Biochem. Biophys. 79, 497–508 (2021).

26. Makide, K. & Aoki, J. GPR34 as a lysophosphatidylserine receptor. J. Biochem. 153, 327–329 (2013).

27. Ritscher, L. et al. The ligand specificity of the G-protein-coupled receptor GPR34. Biochem. J. 443, 841–850 (2012).

28. Wang, X. et al. GPR34-mediated sensing of lysophosphatidylserine released by apoptotic neutrophils activates type 3 innate lymphoid cells to mediate tissue repair. Immunity 54, 1123–1136.e8 (2021).

29. Sayo, A. et al. GPR34 in spinal microglia exacerbates neuropathic pain in mice. J. Neuroinflammation 16, 82 (2019).

30. Forner, S. et al. Systematic phenotyping and characterization of the 5xFAD mouse model of Alzheimer’s disease. Sci. Data 8, 270 (2021).

31. Oakley, H. et al. Intraneuronal β-Amyloid Aggregates, Neurodegeneration, and Neuron Loss in Transgenic Mice with Five Familial Alzheimer’s Disease Mutations: Potential Factors in Amyloid Plaque Formation. J. Neurosci. 26, 10129–10140 (2006).

32. Dolan, M.-J. et al. Exposure of iPSC-derived human microglia to brain substrates enables the generation and manipulation of diverse transcriptional states in vitro. Nat. Immunol. 24, 1382–1390 (2023).

33. Habib, N. et al. Disease-associated astrocytes in Alzheimer’s disease and aging. Nat. Neurosci. 23, 701–706 (2020).

34. Kenigsbuch, M. et al. A shared disease-associated oligodendrocyte signature among multiple CNS pathologies. Nat. Neurosci. 25, 876–886 (2022).

35. Pandey, S. et al. Disease-associated oligodendrocyte responses across neurodegenerative diseases. Cell Rep. 40, 111189 (2022).

36. Thrupp, N. et al. Single-Nucleus RNA-Seq Is Not Suitable for Detection of Microglial Activation Genes in Humans. Cell Rep. 32, 108189 (2020).

37. Robinson, M. D., McCarthy, D. J. & Smyth, G. K. edgeR: a Bioconductor package for differential expression analysis of digital gene expression data. Bioinformatics 26, 139–140 (2009).

38. Strimmer, K. fdrtool: a versatile R package for estimating local and tail area-based false discovery rates. Bioinformatics 24, 1461–1462 (2008).

39. Gosselin, D. et al. An environment-dependent transcriptional network specifies human microglia identity. Science 356, (2017).

40. Abud, E. M. et al. iPSC-Derived Human Microglia-like Cells to Study Neurological Diseases. Neuron 94, 278–293.e9 (2017).

41. Sayama, M. et al. Probing the Hydrophobic Binding Pocket of G-Protein-Coupled Lysophosphatidylserine Receptor GPR34/LPS1 by Docking-Aided Structure–Activity Analysis. J. Med. Chem. 60, 6384–6399 (2017).

42. Gedam, M. & Zheng, H. Complement C3aR signaling: Immune and metabolic modulation and its impact on Alzheimer’s disease. Eur. J. Immunol. 54, e2350815 (2024).

43. Ramaswami, G. et al. Transcriptional characterization of iPSC-derived microglia as a model for therapeutic development in neurodegeneration. Sci. Rep. 14, 2153 (2024).

44. Schulz, A. & Schöneberg, T. The Structural Evolution of a P2Y-like G-protein-coupled Receptor*. J. Biol. Chem. 278, 35531–35541 (2003).

45. Xia, A., et al. Cryo-EM structures of human GPR34 enable the identification of selective antagonists. Proc. Natl. Acad. Sci. 120, e2308435120 (2023).

46. Du, C. et al. Synaptotagmin - 11 inhibits cytokine secretion and phagocytosis in microglia. Glia 65, 1656–1667 (2017).

47. Grajchen, E. et al. CD36-mediated uptake of myelin debris by macrophages and microglia reduces neuroinflammation. J. Neuroinflammation 17, 224 (2020).

48. Loix, M. et al. Perilipin-2 limits remyelination by preventing lipid droplet degradation. Cell. Mol. Life Sci. 79, 515 (2022).

49. Quick, J. D. et al. Lysosomal acidification dysfunction in microglia: an emerging pathogenic mechanism of neuroinflammation and neurodegeneration. J. Neuroinflammation 20, 185 (2023).

50. Flowers, A., Bell-Temin, H., Jalloh, A., Stevens, S. M. & Bickford, P. C. Proteomic analysis of aged microglia: shifts in transcription, bioenergetics, and nutrient response. J. Neuroinflammation 14, 96 (2017).

51. Dräger, N. M. et al. A CRISPRi/a platform in human iPSC-derived microglia uncovers regulators of disease states. Nat. Neurosci. 25, 1149–1162 (2022).

52. Stogsdill, J. A. et al. Pyramidal neuron subtype diversity governs microglia states in the neocortex. Nature 608, 750–756 (2022).

53. Vietri, M., Radulovic, M. & Stenmark, H. The many functions of ESCRTs. Nat. Rev. Mol. Cell Biol. 21, 25–42 (2020).

54. Moloney, E. B., Moskites, A., Ferrari, E. J., Isacson, O. & Hallett, P. J. The glycoprotein GPNMB is selectively elevated in the substantia nigra of Parkinson’s disease patients and increases after lysosomal stress. Neurobiol. Dis. 120, 1–11 (2018).

55. Bogacki, E. C., Longmore, G., Lewis, P. A. & Herbst, S. GPNMB is a biomarker for lysosomal dysfunction and is secreted via LRRK2-modulated lysosomal exocytosis. bioRxiv 2025.01.01.630988 (2025) doi:10.1101/2025.01.01.630988.

56. Almolda, B., Gonzalez, B. & Castellano, B. Antigen presentation in EAE: role of microglia, macrophages and dendritic cells. Front. Biosci. 16, 1157 (2011).

57. Perlmutter, L. S., Scott, S. A., Barrón, E. & Chui, H. C. MHC class II - positive microglia in human brain: Association with alzheimer lesions. J. Neurosci. Res. 33, 549–558 (1992).

58. Sheffield, L. G. & Berman, N. E. J. Microglial Expression of MHC Class II Increases in Normal Aging of Nonhuman Primates. Neurobiol. Aging 19, 47–55 (1998).

59. Dusi, S. et al. Exome Sequence Reveals Mutations in CoA Synthase as a Cause of Neurodegeneration with Brain Iron Accumulation. Am. J. Hum. Genet. 94, 11–22 (2014).

60. Nadig, A. et al. Transcriptome-wide analysis of differential expression in perturbation atlases. Nat. Genet. 57, 1228–1237 (2025).

61. Marsh, S. E. et al. Dissection of artifactual and confounding glial signatures by single-cell sequencing of mouse and human brain. Nat. Neurosci. 25, 306–316 (2022).

62. Kaye, S. et al. Hypercholesterolemia drives microglial dysfunction and weakens response to amyloid plaques. Exp. Neurol. 390, 115272 (2025).

63. Lindholm, D., Bornhauser, B. C. & Korhonen, L. Mylip makes an Idol turn into regulation of LDL receptor. Cell. Mol. Life Sci. 66, 3399–3402 (2009).

64. Manavalan, A. P. C. et al. Phospholipid Transfer Protein Is Expressed in Cerebrovascular Endothelial Cells and Involved in High Density Lipoprotein Biogenesis and Remodeling at the Blood-Brain Barrier*. J. Biol. Chem. 289, 4683–4698 (2014).

65. Villa, M., Wu, J., Hansen, S. & Pahnke, J. Emerging Role of ABC Transporters in Glia Cells in Health and Diseases of the Central Nervous System. Cells 13, 740 (2024).

66. Lin, L.-L. et al. GPR34 Knockdown Relieves Cognitive Deficits and Suppresses Neuroinflammation in Alzheimer’s Disease via the ERK/NF-κB Signal. Neuroscience 528, 129–139 (2023).

67. Lin, B. et al. GPR34 senses demyelination to promote neuroinflammation and pathologies. Cell. Mol. Immunol. 21, 1131–1144 (2024).

68. Izume, T. et al. Structural basis for lysophosphatidylserine recognition by GPR34. bioRxiv 2023.02.15.528751 (2023) doi:10.1101/2023.02.15.528751.

69. Depp, C. White matters arising: Rethinking Alzheimer’s disease through the lens of myelin. Science 390, 483–483 (2025).

70. Papuć, E. & Rejdak, K. The role of myelin damage in Alzheimer’s disease pathology. Arch. Méd. Sci. : AMS 16, 345–351 (2018).

71. Gong, Z. et al. Lower myelin content is associated with more rapid cognitive decline among cognitively unimpaired individuals. Alzheimer’s Dement. 19, 3098–3107 (2023).

72. Chen, J.-F. et al. Enhancing myelin renewal reverses cognitive dysfunction in a murine model of Alzheimer’s disease. Neuron 109, 2292–2307.e5 (2021).

73. Zhan, X. et al. Myelin injury and degraded myelin vesicles in Alzheimer’s disease. Curr. Alzheimer Res. 11, 232–238 (2014).

74. Zhan, X. et al. Myelin Basic Protein Associates with AβPP, Aβ1-42, and Amyloid Plaques in Cortex of Alzheimer’s Disease Brain. J. Alzheimer’s Dis. 44, 1213–1229 (2015).

75. Diaz, C. et al. A Strategy Combining Differential Low - Throughput Screening and Virtual Screening (DLS - VS) Accelerating the Discovery of new Modulators for the Orphan GPR34 Receptor. Mol. Inform. 32, 213–229 (2013).

76. Ikubo, M. et al. Structure–Activity Relationships of Lysophosphatidylserine Analogs as Agonists of G-Protein-Coupled Receptors GPR34, P2Y10, and GPR174. J. Med. Chem. 58, 4204–4219 (2015).

77. Liu, G., Li, X., Wang, Y., Zhang, X. & Gong, W. Structural basis for ligand recognition and signaling of the lysophosphatidylserine receptors GPR34 and GPR174. PLOS Biol. 21, e3002387 (2023).

78. Cheng, Q. et al. TREM2-activating antibodies abrogate the negative pleiotropic effects of the Alzheimer’s disease variant Trem2 R47H on murine myeloid cell function. J. Biol. Chem. 293, 12620–12633 (2018).

79. Ikezu, T. et al. P2RX7 regulates tauopathy progression via tau and mitochondria loading in extracellular vesicles. Res. Sq. rs.3.rs-6771517 (2025) doi:10.21203/rs.3.rs-6771517/v1.

80. McQuade, A. et al. Gene expression and functional deficits underlie TREM2-knockout microglia responses in human models of Alzheimer’s disease. Nat. Commun. 11, 5370 (2020).

81. Wang, S. et al. Anti-human TREM2 induces microglia proliferation and reduces pathology in an Alzheimer’s disease model. J. Exp. Med. 217, e20200785 (2020).

82. Wang, Y. et al. TPL2 kinase activity regulates microglial inflammatory responses and promotes neurodegeneration in tauopathy mice. eLife 12, e83451 (2023).

## References

1. Picelli, S. et al. Full-length RNA-seq from single cells using Smart-seq2. Nat. Protoc. 9, 171–181 (2014).

2. Zheng, G. X. Y. et al. Massively parallel digital transcriptional profiling of single cells. Nat. Commun. 8, 14049 (2017).

3. Lun, A. T. L. et al. EmptyDrops: distinguishing cells from empty droplets in droplet-based single-cell RNA sequencing data. Genome Biol. 20, 63 (2019).

4. Hao, Y. et al. Dictionary learning for integrative, multimodal and scalable single-cell analysis. Nat. Biotechnol. 42, 293–304 (2024).

5. Nicol, P. B. & Miller, J. W. Model-based dimensionality reduction for single-cell RNA-seq using generalized bilinear models. Biostat. (Oxf., Engl.) 26, kxaf024 (2025).

6. Bais, A. S. & Kostka, D. scds: Computational Annotation of Doublets in Single Cell RNA Sequencing Data. bioRxiv 564021 (2019) doi:10.1101/564021.

7. Depp, C. et al. Myelin dysfunction drives amyloid-β deposition in models of Alzheimer’s disease. Nature 618, 349–357 (2023).

8. Phipson, B. et al. propeller: testing for differences in cell type proportions in single cell data. Bioinformatics 38, 4720–4726 (2022).

9. Robinson, M. D., McCarthy, D. J. & Smyth, G. K. edgeR: a Bioconductor package for differential expression analysis of digital gene expression data. Bioinformatics 26, 139–140 (2009).

10. Korsunsky, I., Nathan, A., Millard, N. & Raychaudhuri, S. Presto scales Wilcoxon and auROC analyses to millions of observations. bioRxiv 653253 (2019) doi:10.1101/653253.

11. Keren-Shaul, H. et al. A Unique Microglia Type Associated with Restricting Development of Alzheimer’s Disease. Cell 169, 1276–1290.e17 (2017).

12. Mandell, K. A. P. et al. Meta-analysis of the brain transcriptomes of multiple genetic mouse models of schizophrenia highlights dysregulation in striatum and thalamus. Transl. Psychiatry 15, 345 (2025).

13. Dobin, A. et al. STAR: ultrafast universal RNA-seq aligner. Bioinformatics 29, 15–21 (2012).

14. Patro, R., Duggal, G., Love, M. I., Irizarry, R. A. & Kingsford, C. Salmon provides fast and bias-aware quantification of transcript expression. Nat. Methods 14, 417–419 (2017).

15. Soneson, C., Love, M. I. & Robinson, M. D. Differential analyses for RNA-seq: transcript-level estimates improve gene-level inferences. F1000Research 4, 1521 (2016).

16. Love, M. I., Huber, W. & Anders, S. Moderated estimation of fold change and dispersion for RNA-Seq data with DESeq2. bioRxiv 002832 (2014) doi:10.1101/002832.

17. Nadig, A. et al. Transcriptome-wide analysis of differential expression in perturbation atlases. Nat. Genet. 57, 1228–1237 (2025).

18. Ritchie, M. E. et al. limma powers differential expression analyses for RNA-sequencing and microarray studies. Nucleic Acids Res. 43, e47–e47 (2015).

19. Korotkevich, G. et al. Fast gene set enrichment analysis. bioRxiv 060012 (2021) doi:10.1101/060012.

20. Viechtbauer, W. Conducting Meta-Analyses in R with the metafor Package. J. Stat. Softw. 36, (2010).

21. Kreitzer, F. R. et al. A robust method to derive functional neural crest cells from human pluripotent stem cells. Am. J. stem cells 2, 119–31 (2013).

22. Dolan, M.-J. et al. Exposure of iPSC-derived human microglia to brain substrates enables the generation and manipulation of diverse transcriptional states in vitro. Nat. Immunol. 24, 1382–1390 (2023).

23. Subramanian, A. et al. Gene set enrichment analysis: A knowledge-based approach for interpreting genome-wide expression profiles. Proc. Natl. Acad. Sci. 102, 15545–15550 (2005).

24. Liberzon, A. et al. Molecular signatures database (MSigDB) 3.0. Bioinform. (Oxf., Engl.) 27, 1739–40 (2011).

25. Erwig, M. S. et al. Myelin: Methods for Purification and Proteome Analysis. Methods Mol. Biol. (Clifton, NJ) 1936, 37–63 (2019).

